# Highly efficient homology-directed repair using transient CRISPR/Cpf1-geminiviral replicon in tomato

**DOI:** 10.1101/521419

**Authors:** Tien Van Vu, Velu Sivankalyani, Eun-Jung Kim, Duong Thi Hai Doan, Mil Thi Tran, Jihae Kim, Yeon Woo Sung, Minwoo Park, Yang Jae Kang, Jae-Yean Kim

## Abstract

Genome editing via the homology-directed repair (HDR) pathway in somatic plant cells is very inefficient compared to error-prone repair by nonhomologous end joining (NHEJ). Here, we increased HDR-based genome editing efficiency approximately 3-fold compared to a Cas9-based single-replicon system via the use of *de novo* multi-replicon systems equipped with CRISPR/LbCpf1 in tomato and obtained replicon-free but stable HDR alleles. The efficiency of CRISPR/LbCpf1-based HDR was significantly modulated by physical culture conditions such as temperature and light. Ten days of incubation at 31°C under a light/dark cycle after *Agrobacterium*-mediated transformation resulted in the best performance among the tested conditions. Furthermore, we developed our single-replicon system into a multi-replicon system that effectively increased HDR efficiency. Although this approach is still challenging, we showed the feasibility of HDR-based genome editing of a salt-tolerant SlHKT1;2 allele without genomic integration of antibiotic markers or any phenotypic selection. Self-pollinated offspring plants carrying the HKT1;2 HDR allele showed stable inheritance and germination tolerance in the presence of 100 mM NaCl. Our work may pave the way for transgene-free editing of alleles of interest in asexually as well as sexually reproducing plants.

## INTRODUCTION

In plant somatic cells, double-strand DNA breaks (DSBs) are efficiently repaired by a nonhomologous end joining (NHEJ) mechanism, which dominates over the homology-directed repair (HDR) pathway (Jiang et al., 2013; Puchta, 2005). NHEJ repair usually leads to various types of mutations including DNA sequence insertions, deletions (Hsu et al., 2014; Zetsche et al., 2015), chromosome rearrangement, or chromosome relocation (Ferguson and Alt, 2001; Richardson et al., 1998; Varga and Aplan, 2005). Early in the 1990s, a transgenic approach using yeast mitochondrial I-Sce I endonuclease as a DSB inducer was adopted in attempts to investigate the mechanisms of DSB repair in plants, especially gene targeting via the HDR pathway in plant somatic cells (Fauser et al., 2012; Puchta et al., 1993), which have been the main targets of recent plant genome engineering approaches (Baltes et al., 2014; Belhaj et al., 2013; Čermák et al., 2015; Nekrasov et al., 2013). In plant somatic cells, the HDR pathway employs homologous DNA templates to precisely repair damaged DNA, mainly via the synthesis-dependent strand annealing (SDSA) mechanism, with an extremely low efficiency (Puchta et al., 1996; Szostak et al., 1983), leading to difficulties in practical applications. Therefore, research on plant gene targeting has continued to focus on improving HDR efficacy. Previously reported data have indicated two most important factors affecting HDR efficiency in plant somatic cells: DSB formation and the amount of homologous DNA templates available at sites of breakage (Baltes et al., 2014; Endo et al., 2016; Puchta, 2005; Puchta et al., 1993; Townsend et al., 2009).

The recent development of the clustered regularly interspaced short palindromic repeats (CRISPR)/CRISPR-associated (Cas) protein system has provided excellent molecular scissors for the generation of DSBs. *Streptococcus pyogenes* Cas 9 (SpCas9) (Sapranauskas et al., 2011) and *Lachnospiraceae bacterium* Cas12a (LbCas12a or LbCpf1) (Zetsche et al., 2015) have been adapted for wide use in genome engineering studies in various kingdoms including Plantae (Barrangou and Doudna, 2016; Hsu et al., 2014; Jinek et al., 2012). The former system generally generates blunt ends (Jinek et al., 2012) at DSBs, while the latter cuts in a cohesive end configuration (Zetsche et al., 2015). As a consequence of DSB repair by NHEJ, the two types of CRISPR complexes exhibit comparably high indel mutation rates under *in vivo* conditions, thus proving to be ideal tools for DSB formation for initiating targeted HDR in plants. Furthermore, it has been suggested that the Cpf1 complex might present an advantage in HDR-based genome editing compared to the Cas9 complex because the cutting site of Cpf1 is located distal to the core target sequence and the protospacer-adjacent motif (PAM), potentially allowing recutting even after indel mutations are introduced during NHEJ-mediated repair (Zetsche et al., 2015; Lowder et al., 2016). CRISPR/Cpf1 complexes were recently successfully applied for gene targeting in plants (Li et al., 2018), providing alternative options for T-rich target site selection.

Because of the highly efficient replication of geminivirus genomes and their single-stranded DNA nature, these genomes have been used as an ideal DNA template carrier for gene targeting in plants. Geminiviral genomic DNAs have been reconstructed to exogenously overexpress foreign proteins in plants at up to 80-fold higher levels compared to those of conventional T-DNA systems (Mor et al., 2003; Needham et al., 1998; Zhang and Mason, 2006), due to their highly autonomous replication inside host nuclei and the ability to reprogram cells (Gutierrez, 1999; Hanley-Bowdoin et al., 2013). Furthermore, Rep/RepA has been reported to promote a cell environment that is permissive for homologous recombination to stimulate the replication of viral DNA. Interestingly, it has been reported that somatic homologous recombination is promoted by geminiviral infection (Richter et al., 2014). The above characteristics of geminiviral replicons have been shown to make them ideal delivery tools for introducing large amounts of homologous donor templates to plant nuclei. Likewise, a bean yellow dwarf virus (BeYDV)– based replicon was developed by replacing its movement protein and coat protein genes with Cas9 or TALEN to improve gene targeting in plants (Baltes et al., 2014; Butler et al., 2016; Čermák et al., 2015; Dahan-Meir et al., 2018; Gil-Humanes et al., 2017; Hummel et al., 2018). The LbCpf1 complex, which was subsequently discovered and adapted for genome editing in 2015, has not been tested in combination with geminiviral replicon systems for plant gene targeting.

Despite higher success rates in gene targeting in plants using the geminiviral replicon system, most of the reported cases have required selection markers associated with the edited alleles, indicating that plant gene targeting without the use of selection markers is still challenging (Butler et al., 2016; Gil-Humanes et al., 2017; Hummel et al., 2018). In addition, the effective application of replicon cargos in plant gene targeting has been shown to be limited by their size (Baltes et al., 2014; Suarez-Lopez and Gutierrez, 1997). Therefore, plant gene targeting, especially in cases of selection marker-free alleles, still requires improvement. Here, we report significant improvement of homology-directed repair using CRISPR/LbCpf1-geminiviral multi-replicons in tomato and the successful application of the system to target a marker-free salt-tolerant HKT1;2 allele.

## RESULTS AND DISCUSSION

### The CRISPR/LbCpf1-based geminiviral replicon system is feasible for performing HDR in tomato

To test the hypothesis above, we re-engineered a BeYDV replicon to supply a high dose of homologous donor templates and used the CRISPR/LbCpf1 system (Zetsche et al., 2015) for DSB formation (Figure 1A and 1B). Two long intergenic regions (LIR) of BeYDV (pLSLR) (Baltes et al., 2014) were cloned in the same orientation with a short intergenic region (SIR) inserted between them, generating an LIR-SIR-LIR amplicon unit (Data S1). To support the autonomous replication of the replicon, the Rep/RepA coding sequence was also introduced in *cis* (in the center of the 3’ side, SIR-LIR) and transcriptionally driven by the bidirectional promoter activity of the LIR. This cloning strategy interrupted a possible upstream ORF of Rep/RepA and added an AAA Kozak consensus sequence (Kozak, 1981) upstream of the major ATG of Rep (Supplemental Figure 1A and 1B; Data S1), thus potentially contributing to increasing the translation of the Rep protein (Barbosa et al., 2013; Zhang et al., 2018). The selection of HDR events was performed with a double selection/screening system based on kanamycin resistance and anthocyanin overproduction (Figure 1A; Supplemental Figure 2A and 2B; Data S1).

**Figure 1.**
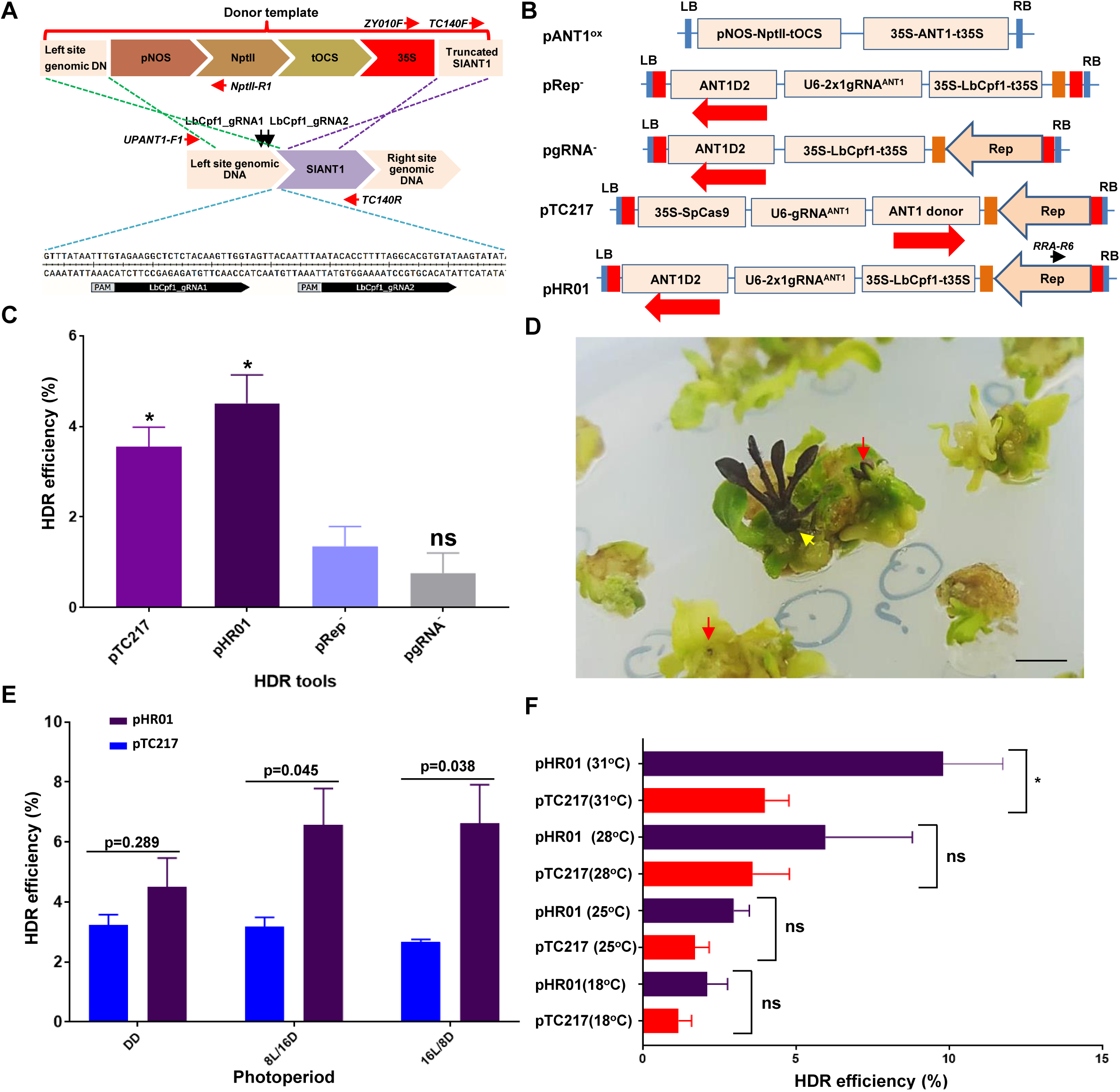
HDR-based genome editing of the ANT1 locus. (**A**) Representatives of ANT1 targeting sites and homologous DNA donor template construction. The upstream sequence of the ANT1 locus (middle panel) was selected for targeting by HDR. The kanamycin expression cassette (pNOS-NptII-tOCS) and CaMV 35S promoter were designed to be inserted at a position 142 bp upstream of the ANT1 start codon. The cutting sites of the two guide RNAs used in this study are indicated by two black arrows. The sequences of the gRNAs are shown in the bottom panel. The red arrows show the relative binding sites and orientations of the primers used for analyses of HDR events. (**B**) T-DNA constructs used for HDR improvement experiments. The dual-guide RNA scaffold (2×1 gRNA^ANT1^) was driven by the *Arabidopsis* U6 promoter core element (75 bp). The LbCpf1 expression cassette was re-engineered to contain the *Arabidopsis* Ubiquitin 1 intron I downstream of the CaMV 35S promoter and upstream of LbCpf1 and to be terminated by the CaMV 35S terminator (35S-LbCpf1I-t35S). Red and orange boxes indicate long intergenic regions (LIR) and short intergenic regions (SIR) of geminivirus DNA, respectively. The black arrow indicates the relevant binding site and orientation of the RRA-R6 primer for subsequent analyses. The red arrows show the orientation of the ANT1 donor templates (ANT1D2). (**C**) Comparison of HDR efficiency between different constructs. Transformed tomato cotyledon fragments were incubated under continuous darkness at 28°C for the first 10 days after washing. Representative photographs of HDR-edited T0 events indicated by purple calli (red arrows) or direct HDR shoot formation (yellow arrow). (**E**) Impact of photoperiod on HDR. The transformed tomato cotyledon fragments were incubated under different lighting regimes at 28°C for the first 10 days after washing. DD: continuous darkness; 8 L/16 D: 8 h light/16 h darkness; 16 L/8 D: 16 h light/8 h darkness. (**F**) HDR efficiencies of the pTC217 and pHR01 constructs obtained at various temperatures. HDR efficiencies were recorded in at least triplicate and were calculated and plotted using PRISM 7.01 software (details of the statistical analyses are described in the Methods section). *: significantly different (p<0.05); ns: not significantly different; p values are shown on the top of the bars of (**E**) for comparison. The data in (**C**), (**E**) and (**F**) are represented as the mean ± SEM.

To validate our system, the LbCpf1 expression cassette driven by the CaMV 35S promoter and 5’UTR with AtUBI10 intron I (to suppress silencing effects (Christie et al., 2011)), guide RNA scaffolds driven by the AtU6 promoter (Data S1) (Belhaj et al., 2013) and donor templates were cloned into the *de novo*-engineered geminiviral DNA replicon (Figure 1B) and transformed via *Agrobacterium*-mediated transformation into tomato cotyledon explants. The *de novo*-engineered geminiviral DNA replicon system exhibited efficient and durable maintenance of circularized DNAs in mature tomato leaves (Supplemental Figure 3). The LbCpf1 system using two guide RNAs for targeting the ANT1 gene, a key transcription factor controlling the anthocyanin pathway, showed a much higher HDR efficiency, of 4.51±0.63% (normalized to an overexpression construct (pANT1^ox^, Figure 1B)), than the other control constructs, including the “minus Rep” (pRep^-^) and “minus gRNA” (pgRNA^-^) constructs (Figure 1C; Supplemental Table 1A). LbCpf1 system-based HDR events was visualized by the presence of purple calli and/or shoots (Figure 1C and 1D), and its efficiency was similar to that of a CRISPR/SpCas9-based construct (pTC217) (Čermák et al., 2015) included in the same experiment (Figure 1C; Supplemental Table 1A) or used in hexaploid wheat with the same scoring method (Gil-Humanes et al., 2017). It is worth noting that the normalized HDR efficiencies reported from this study (see Materials and Methods section) using transformed cell-based efficiency are calculated differently from those reported in the initial work by Čermák and coworkers (2015). The data obtained from this experiment revealed that functional geminiviral replicons were crucial for increasing HDR efficiencies of the Cpf1 complex, as shown for Cas9 system (Čermák et al., 2015). This result shows the feasibility of highly efficient HDR in plants using Cpf1 expressed from a geminiviral replicon, thus expanding the choices of molecular scissor system for gene targeting in plants.

### Favorable physical conditions significantly increase the HDR efficiency of the CRISPR/LbCpf1-based geminiviral replicon system

In seeking suitable physical conditions for *Agrobacterium*-mediated delivery and DSB repair using our HDR tool in tomato somatic cells, we investigated various incubation regimes at early stages post-transformation. Short-day conditions have been shown to have strong impacts on intrachromosomal recombination repair (ICR) in *Arabidopsis* (Boyko et al., 2005). We tested whether the same could be true for the gene targeting approach in tomato. Using various lighting regimes, including complete darkness (DD), short (8 hours light/16 hours dark; 8 L/16 D)- and long (16 L/8 D)-day conditions, we found that the HDR efficiencies achieved under short- and long-day conditions were higher than those under DD conditions in the case of LbCpf1 but not SpCas9 and reached 6.62±1.29% (p<0.05, Figure 1E; Supplemental Table 1B). Considering the similar repair activities observed after DSBs were generated by either of the CRISPR/Cas systems, it was quite difficult to explain why the light conditions only affected LbCpf1-based HDR in this experiment compared to the dark treatment. There must be unknown mechanism(s) that facilitate LbCpf1-mediated HDR in a light-dependent manner.

Temperature is an important factor controlling ICR (Boyko et al., 2005), CRISPR/Cas9-based targeted mutagenesis in plants (LeBlanc et al., 2018), and CRISPR/Cpf1-based HDR in zebrafish and *Xenopus* by controlling genome accessibility (Moreno-Mateos et al., 2017). In addition, Malzahn and coworkers recently reported dependency of Cpf1 cleavage activity on temperature (Malzahn et al., 2019). Pursuing the approach for the improvement of HDR, we compared the HDR efficiencies of the pHR01 and pTC217 systems subjected to various temperature treatments under an 8 L/16 D photoperiod, since the two nucleases (SpCas9 and LbCpf1) may respond differently. Our data revealed that within a temperature range of 19-31°C, the somatic HDR efficiency increased with increasing temperature (Figure 1F; Supplemental Table 1C). Notably, at 31°C, LbCpf1 showed an HDR efficiency (9.80±1.12%) that was more than 2-fold higher than that of SpCas9 (p<0.05) and was nearly twice that of a similar system in hexaploid wheat (Gil-Humanes et al., 2017) as well as an LbCpf1-based T-DNA tool in rice (Li et al., 2018). The results supported the principle of heat stress-stimulated HDR in plants reported by Boyko and coworkers (2005). The ease of LbCpf1 at genome accessibility at high temperatures (Moreno-Mateos et al., 2017) in combination with the ability to repeatedly cut at the target sites (Zetsche et al., 2015) may explain the higher HDR efficiency of LbCpf1 compared to that of SpCas9. The claims are supported by the pHR01 and pTC217 guide RNA activity analyses. The number of transformed events carrying indel mutations was almost similar, but the average adjusted levels of mutation rate obtained from the pHR01-based LbCpf1 (Supplemental Table 2B; Data S2) was nearly three-fold higher than that of the SpCas9-based pTC217 events (Supplemental Table 2A; Data S2). Interestingly, the LbCpf1 complex was shown to be highly active only at high temperatures (i.e., more than 29°C) (Malzahn et al., 2019), which partially explains the higher HDR efficiencies observed at high temperatures in this experiment. It is notable that a highly efficient CRISPR/LbCpf1 mutant in low temperature was reported for plant gene editing (Schindele and Puchta, 2019). Even the LbCpf1_gRNA1 appeared to be highly active at the on-target site; no modification at two potential off-targeting sites was observed (Data S3). Briefly, a comparison of data on plant HDR between Cas9- and Cpf1-based systems at different temperatures and under short-day conditions is presented to reveal the best conditions for plant HDR improvement.

### A multi-replicon system outperformed the single-replicon system in HDR-based GE

The size of viral replicons has been shown to be inversely correlated with their copy numbers (Baltes et al., 2014; Suarez-Lopez and Gutierrez, 1997). In an approach to overcome the replicon size limitation, we designed and tested the novel idea of using a T-DNA system that potentially produces multiple replicons (Figure 2A, and Supplemental Figure 4). Compared to pHR01, a multi-replicon system designed to release donor templates from replicon 2 (MR02) but not replicon 1 (MR01) showed a significant increase in the HDR efficiency by 30% and reached up to 12.79±0.37% (Figure 2B and Supplemental Table 3). Temporal evaluation of donor template levels between the HDR tools showed significantly higher levels of MR02 at 3 days post-transformation (dpt) compared to those of pHR01 and MR01 (Figure 2C). The highest donor template levels in multi-replicons tested were available while CRISPR/Cas was generating DSBs at early times after transformation (3 dpt, Figure 2C), except for MR01 showing a peak at 6 dpt (Figure 2C). Under the same conditions and calculation methods, the combination of our multi-replicons with LbCpf1 significantly increased HDR efficiencies by 3-4-fold compared to those of the Cas9-based replicon systems. We also confirmed the release of three circularized replicons from the single vector used in this work (Figure 2D) by PCR amplification using circularized replicon-specific primers (Supplemental Table 4).

**Figure 2.**
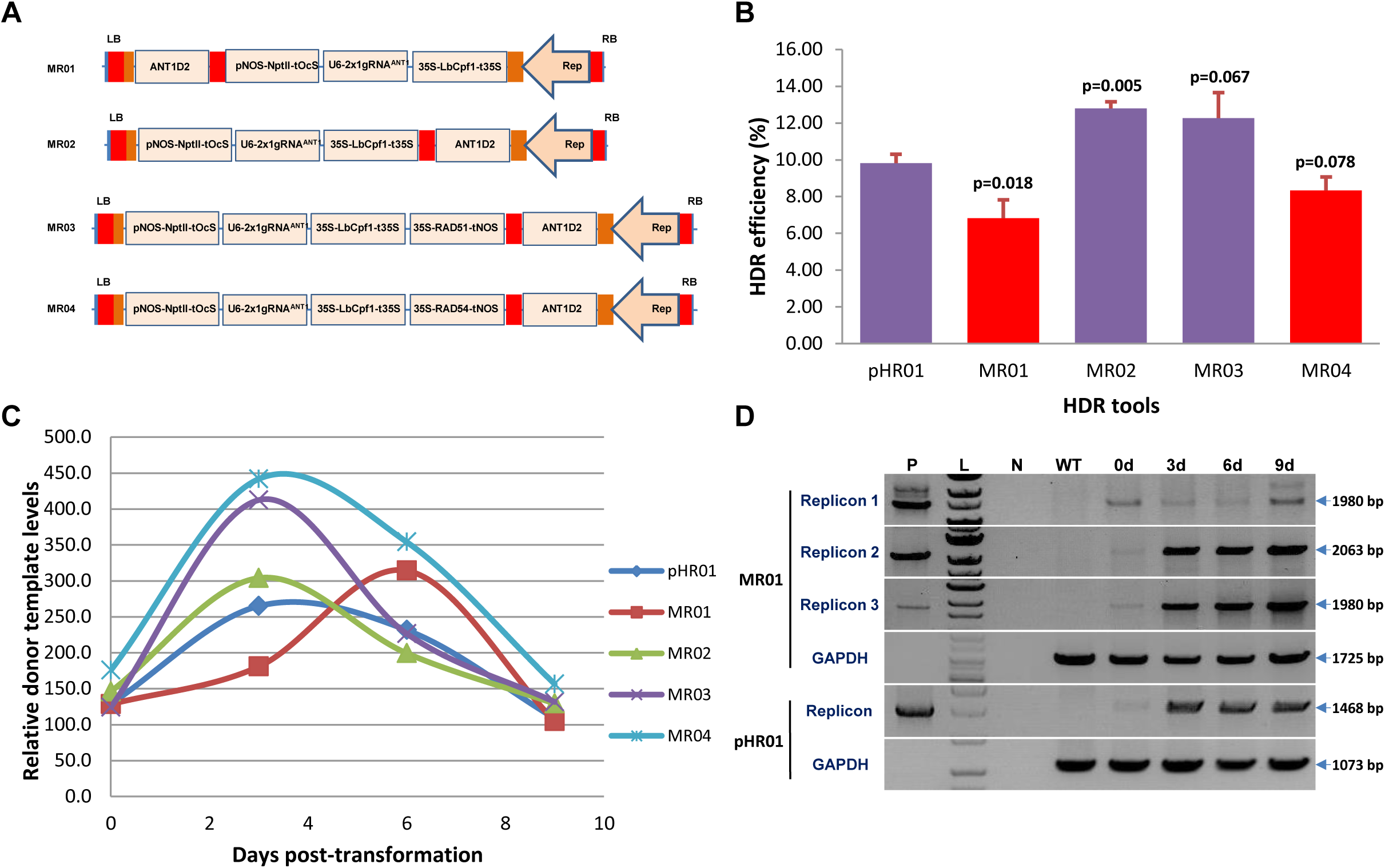
Multi-replicon tools for HDR improvement. (**A**) Multi-replicon constructs tested for the improvement of HDR over NHEJ. Red and orange boxes indicate LIRs and SIRs of geminiviral DNA, respectively. ANT1D2, ANT1 donor templates. (**B**) HDR efficiencies obtained using multi-replicons as cargos for the HDR tools. HDR efficiencies were recorded four times and were calculated and plotted using PRISM 7.01 software (details of the statistical analyses are described in the Materials and Methods section). p values (pairwise comparisons to pHR01 using Student’s test) are shown on the top of the bars. Data are represented as the mean ± SEM. (**C**) Relatively quantified donor template levels at different time points post-transformation by qPCR using ANT1D2 template-specific primers normalized to SlPDS. (**D**) PCR detection of circularized replicons simultaneously released from the MR01 vector. 0d, 3d, 6d and 9d: samples collected at 0, 3, 6 and 9 days post-transformation with *Agrobacterium* carrying MR01. P: pHR01 plasmid isolated from Agrobacteria; L: 1 kb ladder; N: water control; WT: wild-type tomato Hongkwang. The primer pairs used in PCR to detect circularized replicons are shown in Supplemental Figure 4, bottom panel, and Supplemental Table 4.

In another test of the multi-replicon system, we overexpressed two key proteins involved in the plant HDR pathway from the replicon 1 site. Either SlRAD51 (Solyc07g017540.2) or SlRAD54 (Solyc04g056400.2) was overexpressed with the multi-replicon tools (MR03 and MR04) (Figure 2A; Data S1). Surprisingly, even when the donor template level of MR03 or MR04 was nearly twice that of MR01 (Figure 2C), the HDR efficiency was not significantly different in the case of MR03 and was even significantly lower for MR04 (Figure 2B and Supplemental Table 3).The assessment of mRNA levels of SlRAD51 (Supplemental Figure 5A; Data S4) or SlRAD54 (Supplemental Figure 5B; Data S4) of transformed events of MR03 or MR04, respectively, showed higher relative transcript levels (up to 522.18 folds of SlRAD51 and 83.68 folds of SlRAD54 transcripts) compared to multi-replicon control (MR02) (Supplemental Figure 5; Data S4). Overexpression of SlRAD54 might increase the displacement of SlRAD51 from SlRAD51-bound dsDNAs at the early stage of HDR initiation (Petukhova et al., 1999), thereby suppressing HDR to some extent in the case of MR04 (Figure 2B). Overexpression of either SlRAD51 (MR03) or SlRAD54 (MR04) increased the 3-day peaks of geminiviral replicons (replicon 2 and 3) at 30-50 % compared to the control (MR02) (Figure 2C), confirming the positive roles of these proteins in geminivirus replication in a homologous recombination manner, as reported elsewhere (Kaliappan et al., 2012; Richter et al., 2016; Suyal et al., 2013). The data also revealed a temporal difference in the maximal peaks of replicon 1 and 2 because replicon 1 was not accompanied by a Rep/RepA expression cassette.

The multi-replicon system may provide more flexible choices for expressing multiple donor templates/genes/genetic tools in plant cells with temporally controllable copy levels without incurring an expression penalty from excess replicon sizes up to 18 kb (size of replicon 3 released by MR03). The validation of the multi-replicon system provides an excellent alternative for genetic engineering in plants in addition to applications in plant genome editing. If we carefully design and clone multiple donor templates or gene expression cassettes into the multi-replicons, we can control donor templates/gene doses without incurring penalties from excessing replicon size limitations.

### True ANT1 HDR events occurred at high frequency

An important step in plant genome editing is the regeneration of edited calli into shoots. We used kanamycin in our study to select edited calli and plants. Since we put a fully functional NptII expression cassette into the ANT1 donor, we observed many WT-like calli and green shoots arose from our plates. In the case, the purple marker was so much useful for us to select HDR events. Our observation recorded a significantly higher number of both purple spots per cotyledon and purple plants per cotyledon obtained from pHR01 and MR02 compared to that of pTC217 (Supplemental Table 5). However, the regeneration of the purple calli into plants was not completely proportional probably due to pleiotropic impacts of the new replicon systems. To verify HDR repair events, PCR analyses were conducted using primers specific for the left (UPANT1-F1/NptII-R1) and right (ZY010F/TC140R) (Figure 1A; Supplemental Table 6 and 7) junctions employing genomic DNAs extracted from derived HDR events (independently regenerated purple plants or genome-edited generation 0 (GE0)) (Figure 3A, Supplemental Figure 6 and 7). For pHR01, all (16/16) of the analyzed independent events showed the expected band for right junction integration, and 10/16 independent events showed the expected band for left junction repair (Figure 3B). The PCR products were sequenced to identify junction sequences. A majority of the events (11/16) showed sequences corresponding to perfect right arm integration through HDR repair, and 5/16 events showed a combination of HDR and NHEJ repair with an NHEJ fingerprint at the 5’ terminus of the pNOS sequence (Supplemental Figure 7A, with event C1.8 highlighted in blue) or even an integration of the right board of T-DNA at the left junction boundary (Supplemental Figure 8). All of the sequences amplified from the left junctions showed perfected DNA sequence exchange via the HDR pathway (Supplemental Figure 7B). The results obtained in these analyses revealed the common features of products repaired via HDR pathways in plant somatic cells reported elsewhere in dicots (Butler et al., 2016; Čermák et al., 2015; Dahan-Meir et al., 2018) and monocots (Gil-Humanes et al., 2017; Li et al., 2018), regardless of whether a T-DNA or geminiviral replicon system was involved. More importantly, 15 out of 16 events showed no amplification of circularized forms of the DNA replicon, and even the replicon-carrying events lost this replicon after long-term growth in greenhouse conditions (data not shown), indicating that these plants were free of the replicon (Figure 3B). The absence of the replicon might be hypothetically explained by reverse construction of the donor template (Figure 1B), leading to the opposite arrangement of the LIR forward promoter sequence against a 35S promoter sequence (LIR-p35S orientation interference), which triggers a silencing mechanism in plant cells in later stages. This possibility was later supported by the appearance of replicons in the majority of plants regenerated using other replicon systems without LIR-p35S orientation interference.

**Figure 3.**
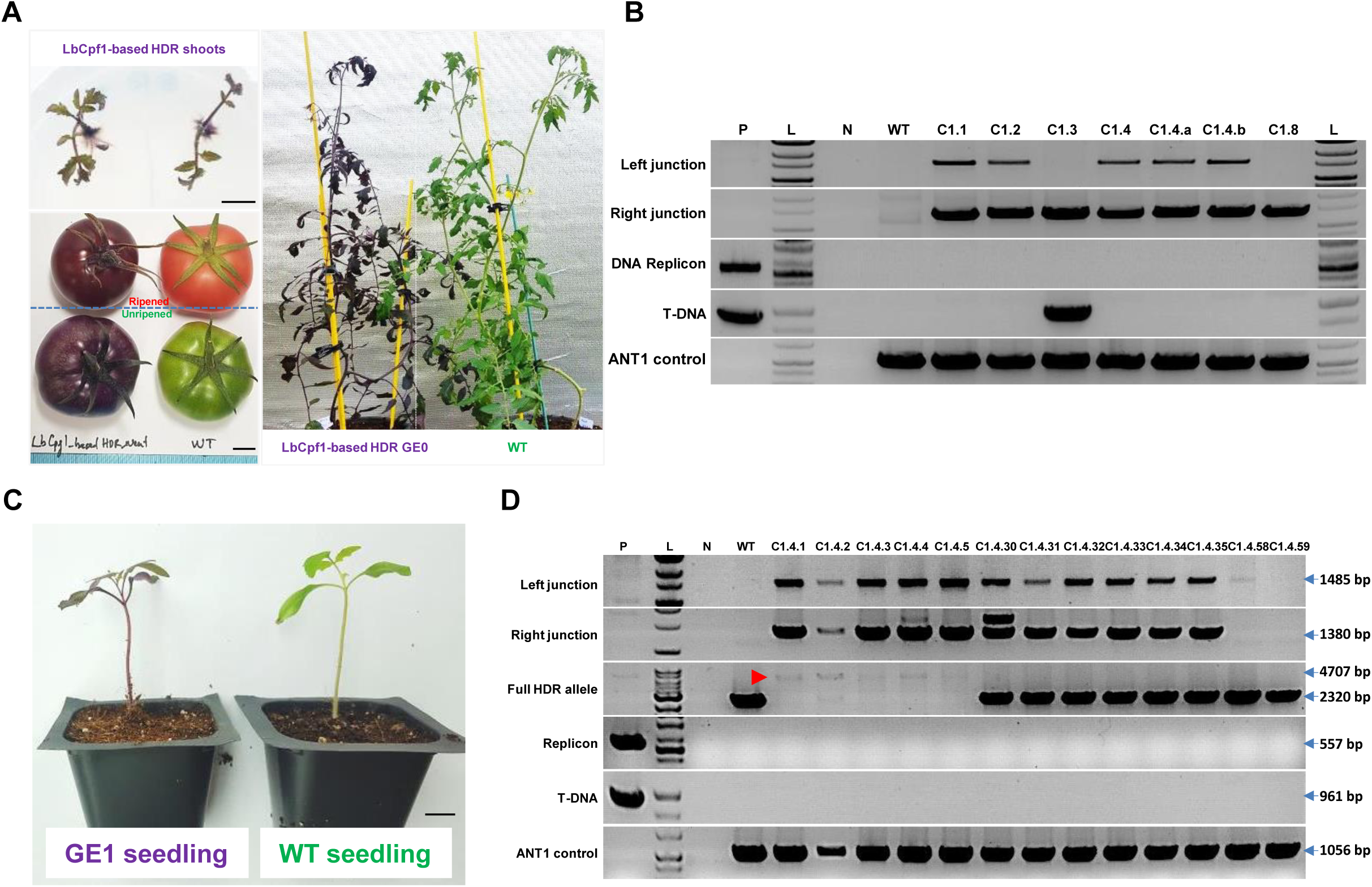
Analyses of HDR-edited plants. (**A**) Representative HDR-edited plants in greenhouse conditions and their fruits. Scale bars = 1 cm. (**B**) PCR analysis data of representative HDR-independent events. P: pHR01 plasmid isolated from Agrobacteria; L: 1 kb ladder; N: water control; WT: wild-type tomato Hongkwang; C1.1, C1.2, C1.3, C1.8: independent LbCpf1-based HDR GE0 events. ANT1 control products were PCR amplified using the TC140F and TC140R primers (Figure 1A) flanking the upstream region of the ANT1 gene. (**C**) Generation 1 of HDR-edited events (GE1). GE1 plants (left) germinated in soil in pots in comparison with wild-type plants (right). Scale bar = 1 cm. (**D**) PCR analysis data of GE1 offspring resulting from C1.4 events. P: pHR01 plasmid isolated from Agrobacteria; L: 1 kb ladder; N: water control; WT: wild-type tomato Hongkwang; C1.4.1, C1.4.2, C1.4.3, C1.4.4 and C1.4.5: GE1 plants showing dark purple color obtained from the self-pollination of plants from the C1.4 event. ANT1 control products were PCR amplified using the TC140F and TC140R primers (Figure 1A) flanking the upstream region of the ANT1 gene. Red arrowhead indicates HDR allele.

### The HDR allele was stably inherited in offspring by self-pollination as well as backcrossing

To validate stable heritable edits, we grew genome-edited generation 1 (GE1) plants (Figure 3C) obtained from the self-pollination of LbCpf1-based HDR GE0 events and identified a segregating population with a purple phenotype (Supplemental Table 8) similar to the segregating profiles shown by Čermák and coworkers (2015). PCR analyses of the segregating plants showed inheritance of the edited allele (Figure 3D and Supplemental Figure 9). The offspring segregated from the #C1.4 event were analyzed in detail. Five dark purple plants (C1.4.1-C1.4.5, homozygous for the ANT1 HDR-edited allele, Supplemental Figure 10), six pale purple plants (C1.4.30-C1.4.35, heterozygous for the ANT1 HDR-edited allele, Supplemental Figure 10), and two wild-type-like plants did not contain the HDR-edited allele, as expected (Figure 3D, predicted results correlated with phenotypes). The dark purple plants showed PCR amplification from the replaced allele but no amplification of the wild-type allele when PCR was performed using primers flanking the editing site (Figure 1A). In contrast, heterozygous and wild-type plants showed a band corresponding to the wild-type allele. Further assessment indicated that the GE2 offspring of the homozygous GE1 plants were all dark purple, and the back-crossed (to WT female as pollen acceptors) BC1F1 generation all showed the pale purple phenotype (Supplemental Figure 10), suggesting the feasibility of recovering the parental genetic background via backcrossing in cases of unexpected modification, including off-target effects. Sanger sequencing revealed perfect inheritance of the HDR-edited allele from the GE0 generation of event C1.4 (Supplemental Figure 11) to its homozygous offspring. We subsequently subjected gDNAs of several putative homozygous as well as heterozygous GE1 lines to Southern blot analysis. The data revealed and confirmed the existence and inheritance of the edited locus in GE1 lines as shown at expected sizes at single HDR band (homozygote) or in combination of HDR and WT bands (heterozygote) (Supplemental Figure 12; Data S5) These data also showed no amplification of circular forms of the DNA replicon (Figure 3D and Supplemental Figure 10), indicating that the GE1 plants were also free of the replicons.

### HDR-based GE using allele-associated marker-free approaches

To show the applicability of our HDR system to practical plant genome editing, we sought to use it to edit a potentially agronomic trait, and salinity tolerance was chosen as the target trait. High-affinity K^+^ Transporter 1;2 (HKT1;2) plays an important role in the maintenance of K+ uptake under salt stress (Ali et al., 2012). Salinity tolerance was determined by a single N/D variant (N217D in tomato) in the pore region of HKT1;2, which determines selectivity for Na+ and K+ (Ali et al., 2016). We succeeded in generating a heterozygous but perfect HDR GE0 event to produce the salt-tolerant allele (N217D) (Ali et al., 2016) (Figure 4A, Supplemental Table 9) according to the analysis of 150 events (∼0.66%) using our system with a *HKT1;2* gene donor template that included neither an allele-associated antibiotic selection marker nor an ANT1 color marker (Figure 4B; Data S1). The CRISPR/LbCpf1 system was very effective for NHEJ repair because it generated indel mutation rates of up to 72% in multiple mutation patterns decomposed by ICE Synthego software (Hsiau et al., 2019) (Supplemental Figure 13A and B), in which most of the events resulted in 47-97% cells carrying indel sequences (Supplemental Table 10). In comparison with the first report on the allele-associated marker-free gene targeting of the CRTISO allele (Dahan-Meir et al., 2018), the HDR frequency obtained with the HKT1;2 locus in this study was much lower, possibly due to (1) lower cutting activity (note the indel mutation rates in Supplemental Table 10), (2) a different target site context or (3) the use of a different strategy to express Rep/RepA (Dahan-Meir and coworkers used a replicon tool with Rep expression driven by a CaMV35S promoter from outside of LIR-SIR-LIR boundary), or to unknown reasons associated with the CRTISO alleles, as claimed by the authors, or all of the above-mentioned factors. We used a similar replicon tool to that reported by Dahan-Meir and coworkers (2018) for ANT1 targeting via the HDR pathway in this study but obtained significantly lower HDR efficiencies than were obtained with the pHR01 tool (data not shown).

**Figure 4.**
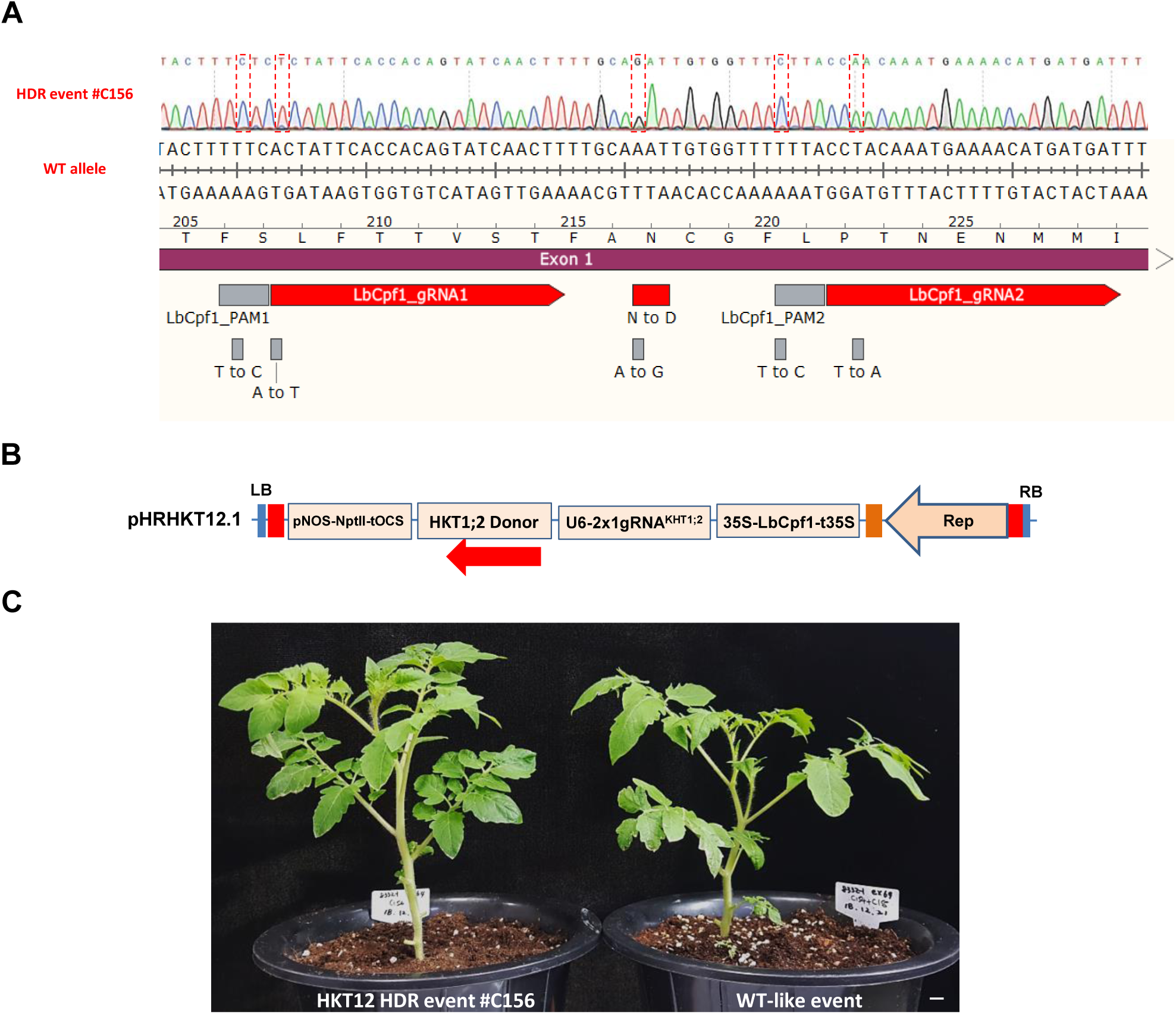
HKT1;2^N217D^ allele editing by HDR using the CRISPR/Cpf1-based replicon system. (**A**) Sanger sequencing of event #C156. Sequence alignment shows the perfectly edited HKT1;2 N217 to D217 allele with the WT allele as a reference. The nucleotides highlighted in the discontinuous red boxes correspond to intended modifications for N217D, PAM and core sequences (to avoid recutting). (**B**) HDR construct layout for HKT1;2 editing. There is neither selection nor a visible marker integrated into the donor sequence. The *Npt*II marker was used for the enrichment of transformed cells. (**C**) Morphology of the HKT1;2^N217D^ edited event compared to its WT-like event in greenhouse conditions. Scale bar = 1 cm.

The editing event involving the D217 allele resulted in a normal morphology (Figure 4C) and normally set fruits (Supplemental Figure 14) compared to WT. It should be noted that the mutated nucleotide (A to G) of *HKT1;2* is not accessible by any currently known base editor (BE), including xCas9-ABE (Hu et al., 2018), highlighting the significance of HDR-based genome editing. We tested the self-pollinated GE1 generation of the plants obtained from the event and observed up to 100 mM NaCl tolerance at the germination stage (Figure 5A) in both homozygous and heterozygous plants. The salt-tolerant plants showed a 3-4-day delay in germination compared to the mock controls but grew normally in NaCl-containing medium (Figure 5A) and later fully recovered in soil (Figure 5B). Screening for the presence of HDR allele(s) in the tested plants via the cleaved amplified polymorphic sequence (CAPS) method showed allele segregation following Mendelian rules (Figure 5C). The true HKT1;2^N217D^ HDR alleles in the GE1 plants were ultimately confirmed by Sanger sequencing. It is worth noting that most of the elite alleles in plants do not associate with any selection marker, and hence, a highly efficient HDR with allele-associated marker-free system is in high demand.

**Figure 5.**
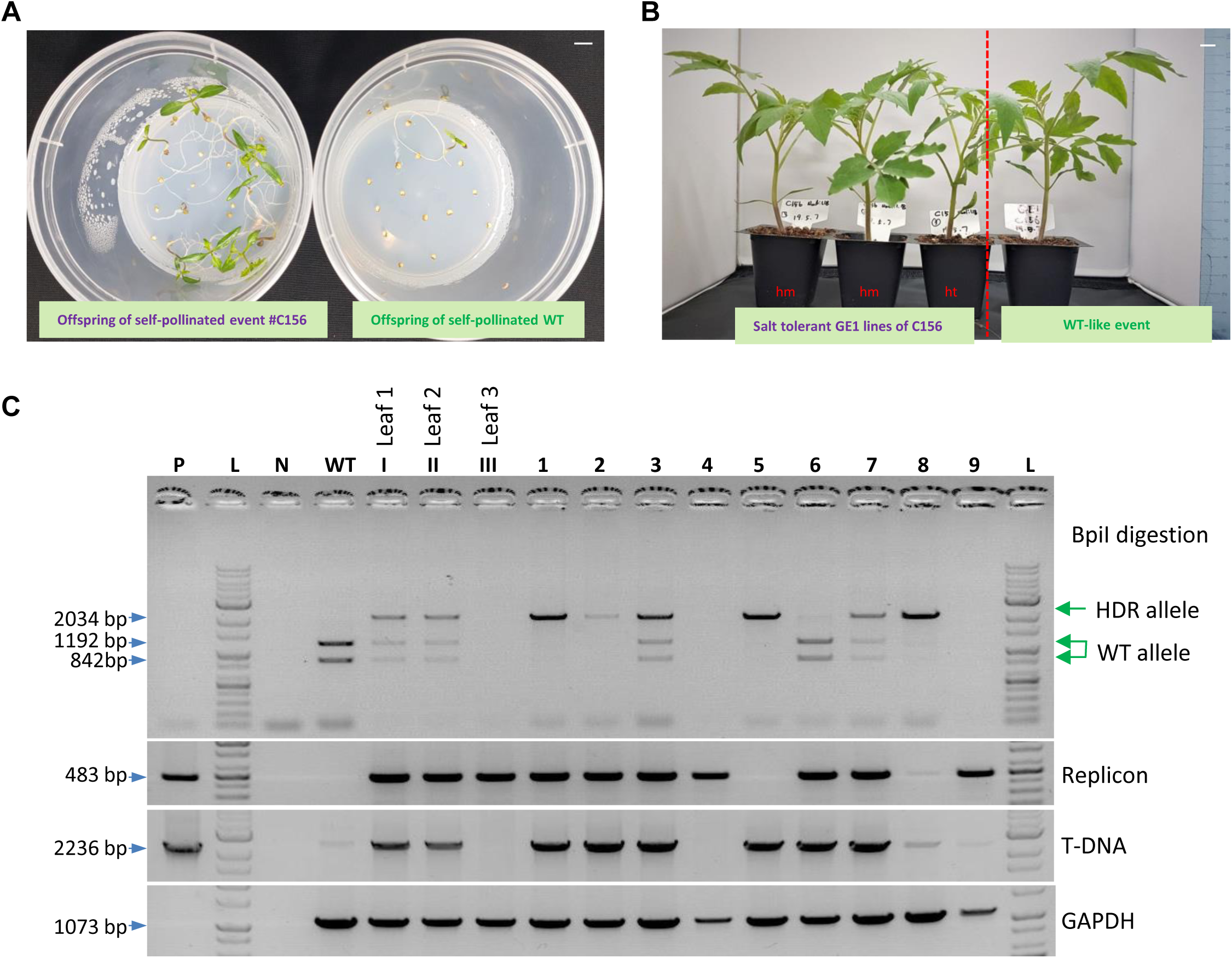
Evaluation of the GE1 offspring of the HKT1;2^N217D^ HDR event. (**A**) Salinity tolerance test at the germination stage using NaCl. Left panel: GE1 plants obtained from self-pollination of the plants obtained from event #C156; right panel: WT control. Bar=1 cm. (**B**) Salt-tolerant plants (left panel) growing in soil showed normal growth compared to WT-like event (right panel). hm=homozygous for the HKT1;2^N211D^ allele; ht=heter^ozygo^us for the HKT1;2^N217D^ allele. Bar=1 cm. (**C**) Screening for the presence of HDR allele(s) in the tested plants via the cleaved amplified polymorphic sequence (CAPS) method. PCR amplification using primers flanking the targeted region was conducted. The PCR products were digested with the BpiI enzyme and resolved in a 1% agarose gel. P: Plasmid control; L: 1 kb ladder; WT: wild-type sample; Leaf 1, Leaf 2 and Leaf 3: samples collected from three different positions (angles) on the C156 plants. 1-9: GE1 plants of C156.

Thus, through the application of various approaches, our study showed a significant improvement of HDR efficiency in tomato somatic cells. The HDR allele was stably inherited in subsequent generations obtained via self-pollination and backcrossing. The advancement of HDR in somatic cells, the generation of replicon-free HDR-edited plants in the GE0 generation and the invention of multi-replicon system open the door for practical applications of the technique to improve crop traits, with special interest for asexually reproducing crops.

## MATERIALS AND METHODS

### Construction and cloning of HDR testing systems

The entire design principle and all cloning procedures followed MoClo (Weber et al., 2011) and Golden Gate (Engler et al., 2014) protocols. pLSL.R.Ly was designed by amplifying the long intergenic region (LIR), short intergenic region (SIR) and lycopene marker from the pLSLR plasmid (Čermák et al., 2015) and was cloned following the order shown in Supplemental Figure 3A and Data S1. Level 2 Golden Gate BpiI restriction sites flanking the pink marker gene (lycopene) were also integrated within the replicon for the cloning of HDR expression cassettes. The release of circularized DNA replicons was validated in tomato leaves (Supplemental Figure 3B) as well as tomato cotyledon explants (data not shown). The pTC147 and pTC217 plasmids (Čermák et al., 2015) were obtained from Addgene and used as a reference. The LbCpf1-based HDR replicons were designed and cloned similarly to the SpCas9-based constructs, with two guide RNAs (LbCpf1_gRNA1 and LbCpf1_gRNA2, Figure 1A; Data S1). Donor DNAs (ANT1D2) were constructed for the integration of an antibiotic selection marker (NptII) and the insertion of a CaMV 35S promoter to drive overexpression of the ANT1 gene (pANT1^ox^, Figure 1A; Data S1). The dual-guide RNA construct was designed by multiplexing the LbCpf1 crRNAs as a tandem repeat of scaffold RNA followed by 23 nt guide RNA sequences. The crRNAs were driven by an AtU6 promoter (Kamoun Lab, Addgene #46968) and terminated by 7-T chain sequences (Data S1).

### Tomato transformation

Our study of HDR improvement was conducted using tomato (Hongkwang cultivar, a local variety) as a model plant. All the binary vectors were transformed into *Agrobacterium tumefaciens* GV3101 (pMP90) using electroporation. *Agrobacterium*-mediated transformation was used to deliver editing tools to tomato cotyledon fragments (Supplemental Figure 15). Explants for transformation were prepared from 7-day-old cotyledons. Sterilized seeds of the Hongkwang cultivar were grown in MSO medium (half-strength MS medium containing 30 g/L of sucrose, pH 5.8) at 25±2°C under 16 h/8 h light/dark conditions. Seven-day-old seedlings were collected, and their cotyledonary leaves were sliced into 0.2-0.3 cm fragments. The fragments (explants) were pretreated in PREMC medium [MS basal salts, Gamborg B5 vitamins, 2.0 mg/L of Zeatin trans isomer and 0.2 mg/L of indolyl acetic acid (IAA), 1 mM of putrescine and 30 g/L of glucose, pH 5.7] for 1 day. The precultured explants were then pricked and transformed using *A. tumefaciens* GV3101::pMP90 cells carrying HR construct(s).

*A. tumefaciens* GV3101::pMP90 cells were grown in primary culture overnight (LB containing suitable antibiotics) in a shaking incubator at 30°C. Agrobacteria were then collected from the culture (OD 0.6-0.8) by centrifugation. The cells were resuspended in liquid ABM-MS (pH 5.2) and 200 µM acetosyringone. Transformation was carried out for 25 min at RT. The explants were then transferred to cocultivation medium containing all of the components in the ABM-MS medium and 200 µM acetosyringone, pH 5.8. The cocultivation plates were kept in the darkness at 25°C for 2 days, and the explants were then shifted to nonselection medium (NSEL) for 5 days and subcultured in selection medium (SEL5). The nonselection and selection media contained all of the components of the preculture medium as well as 300 mg/L of timentin and 80 mg/L of kanamycin. Subculture of the explants was carried out at 14-day intervals to achieve the best regeneration efficiency. Explants containing purple calli or shoots were then transferred to SEL5R medium (similar to SEL5 but with the zeatin trans isomer concentration reduced to 1.0 mg/L) for further regeneration and/or elongation. When the shoots were sufficiently long (1.5-3.0 cm), they were transferred to rooting medium (containing all of the components of the elongation medium the except zeatin trans isomer plus 1.0 mg/L IBA) to generate intact plants. The intact plants from the rooting medium were transferred to vermiculite pots to allow them to harden before shifting them to soil pots in a greenhouse with a temperature of 26±2°C under a 16 h/8 h photoperiod. The experimental treatment of the physical conditions and data collection were conducted as described in Supplemental Figure 15.

### HDR efficiency calculation

In a previous report, the HDR efficiency calculated by dividing the number of explants containing at least one purple callus (appearing as a purple spot) by the total number of explants obtained from *Agrobacterium*-mediated transformation reached 12% with the replicon system (Čermák et al., 2015). In the present study, purple spots were scored at 21-day post-transformation and HDR efficiencies were calculated differently by normalization of the purple spot numbers per cotyledon explant obtained using genome editing constructs to the purple spot numbers per cotyledon explant counted in case of transformation of the SlANT1 overexpression cassette (pTC147 and pANT1^ox^, Figure 1B) in the same conditions.

### Plant genomic DNA isolation

Tomato genomic DNA isolation was performed using the DNeasy Plant Mini Kit (Qiagen, USA) according to the manufacturer’s protocol. Approximately 200 mg of leaf tissue was crushed in liquid nitrogen using a ceramic mortar and pestle and processed with the kit. Genomic DNA was eluted from the mini spin column with 50-80 µl of TE or nuclease-free water.

### HDR event evaluation

The assessment of gene targeting junctions was performed by conventional PCR using primers flanking the left (UPANT1-F1/NptII-R1) and right (ZY010F/TC140R (Čermák et al., 2015) (Supplemental Table 6 and 7) junctions and a high-fidelity Taq DNA polymerase (Phusion Taq, Thermo Fisher Scientific, USA) and Sanger sequencing (Solgent, Korea). DNA amplicons and related donor template levels were evaluated by semiquantitative PCR and qPCR (using KAPA SYBR FAST qPCR Kits, Sigma-Aldrich, USA), respectively, using primers specific to only circularized replicons and the donor template. Additionally, the qPCR assays were designed and conducted following MIQE’s guidelines, with SlPDS (Solyc03g123760) and SlEF1 (Solyc07g016150) as normalized controls. Analyses of the inherited behavior of the HDR-edited allele were performed with genome-edited generation 1 (GE1) by PCR and Sanger sequencing. Circularized replicons were detected using PCR with the corresponding primers for pHR01 (Supplemental Table 6), multi-replicons (Supplemental Table 4) or pTC217 (Supplemental Table 7).

### Statistical analyses

HDR efficiencies were recorded in at least three replicates and were statistically analyzed and plotted using PRISM 7.01 software. In Figure 1C, multiple comparisons of the HDR efficiencies of the other constructs with that of pRep^-^ were performed by one-way ANOVA (uncorrected Fisher LSD test, n=3, df=2, t=4.4; 4.4 and 1.5 for pTC217; pHR01 and pgRNA^-^, respectively). In Figure 1E, pairwise comparisons of the HDR efficiencies of pTC217 and pHR01 under the three lighting conditions were performed with Student’s t-test (DD: t=1.222, df=4; 8 L/16 D: t=2.424, df=7 and 16 L/8 D: t=3.059, df=4). In Figure 1F, comparisons of the HDR efficiencies of pTC217 and pHR01 in the various temperature conditions were performed with Student’s t-test (19°C: t=2.656, df=2; 25°C: t=3.346, df=2; 28°C: t=2.099, df=5; 31°C: t=4.551, df=2). In Figure 2B, comparisons of the HDR efficiencies of the other multi-replicon tools with pHR01 were performed with Student’s test (MR01: t=3.648, df=3; MR02: t=6.041, df=3; MR03: t=2.032, df=3; MR04: t=1.893, df=3).

## AUTHOR CONTRIBUTIONS

T.V.V., V.S. and J.Y.K. designed the experiments; T.V.V., V.S., E.J.K., M.T.T., J.K., Y.W.S., D.T.H.D and M.P. performed the experiments; T.V.V., Y.J.K. and J.Y.K. analyzed the results; T.V.V. and J.Y.K. wrote the manuscript.

## Acknowledgments

We wish to thank Mrs. Jeong Se Jeong and Mrs. Hyun Jeong Kim for their valuable technical support in this study.

---

**Supplemental Figure 1.**
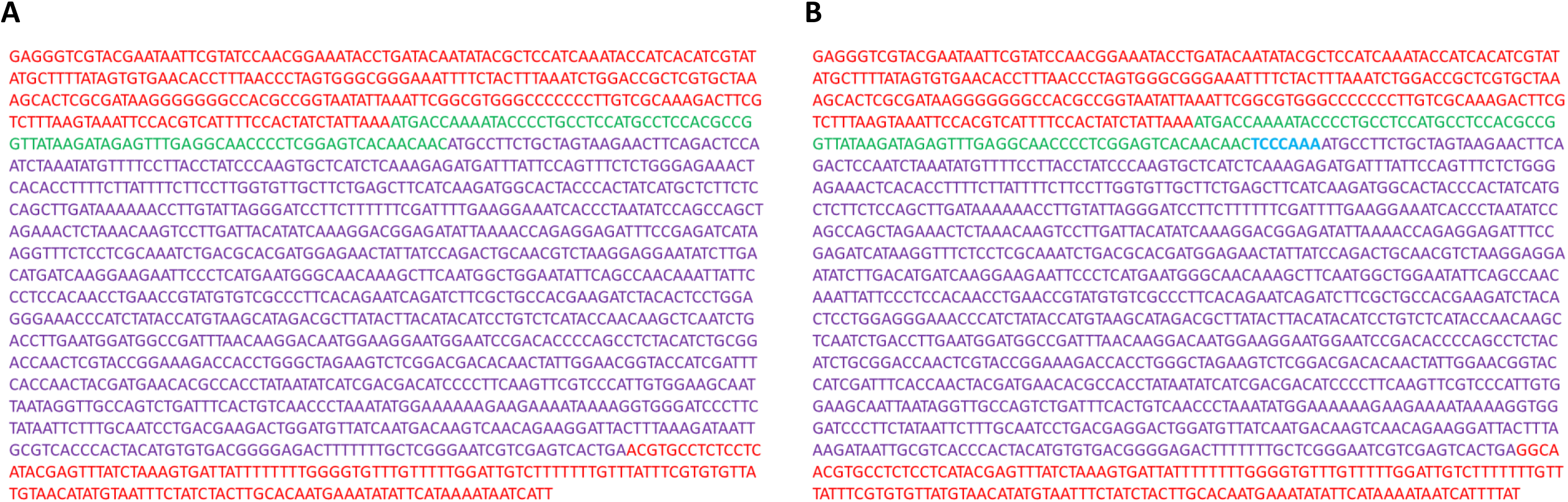
Reengineering of the BeYDV Rep coding sequence used in the study. (**A**) Reverse complement view of the LIR-Rep/RepA-SIR sequence isolated from pLSLR. (**B**) Reverse complement view of the LIR-Rep/RepA-SIR sequence in the *de novo*-engineered replicon used in this study. Upper red font sequences: LIR; bottom red font sequences: SIR; purple sequences: Rep/RepA; green font: upstream ORF sequence (uORF); the light blue sequence TCCCAAA was inserted by cloning to interrupt uORF and add the Kozak preference sequence.

**Supplemental Figure 2.**
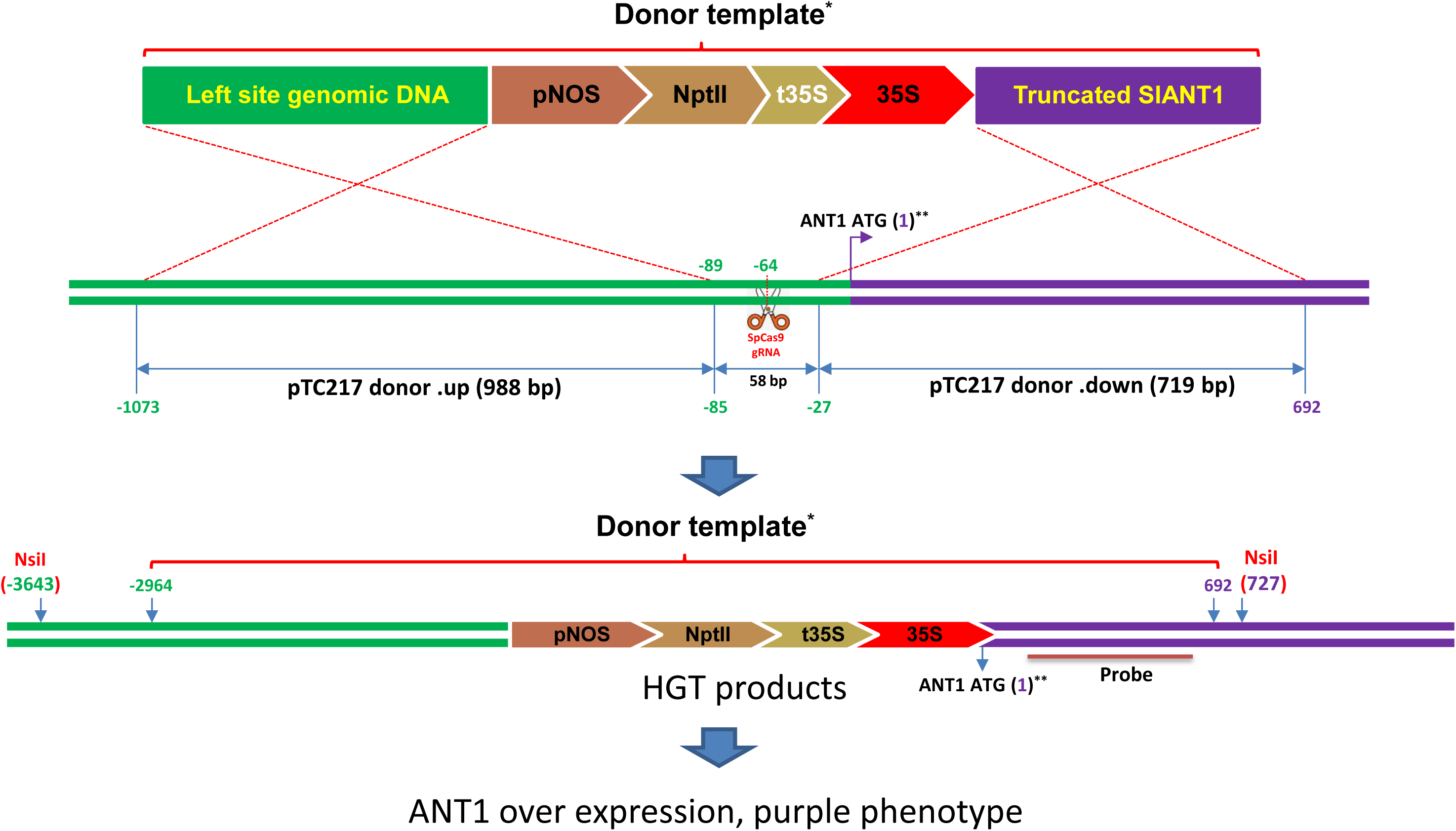
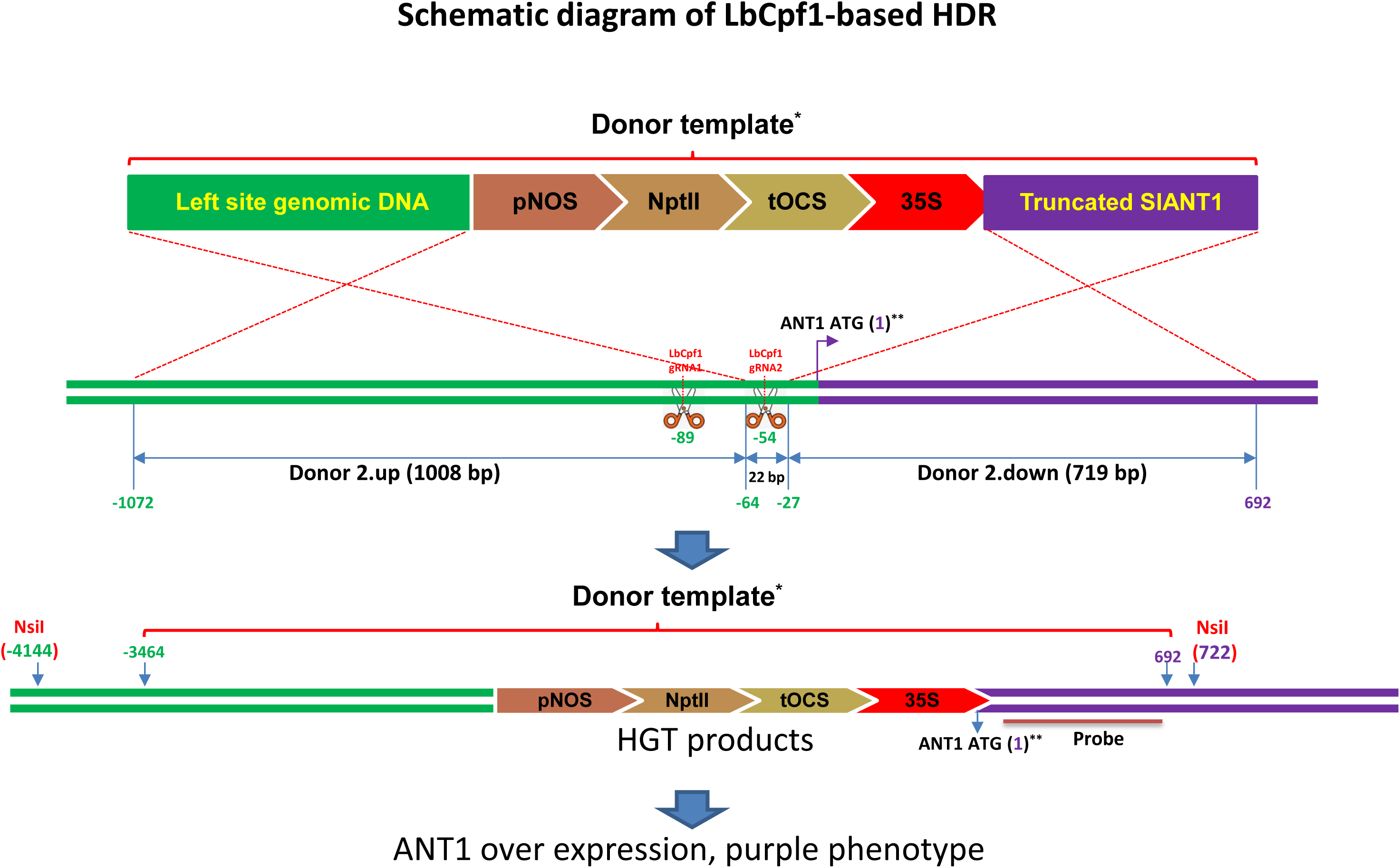
Schematic diagram of HDR-directed editing of ANT1 locus. (**A**) Schematic diagram of SpCas9-based HDR. *Donor length (left arm + insert + right arm): SpCas9 (pTC217): 988 + 1949 + 719 = 3656 bp. **The A nucleotide of start codon of ANT1 is set as +1 position. *Nsi*I sites and probe were used for Southern Blot analysis. (**B**) Schematic diagram of LbCpf1-based HDR. *Donor length (left arm + insert + right arm): LbCpf1: 1008 + 2429 + 719 = 4156 bp. **The A nucleotide of start codon of ANT1 is set as +1 position. *Nsi*I sites and probe were used for Southern Blot analysis.

**Supplemental Figure 3.**
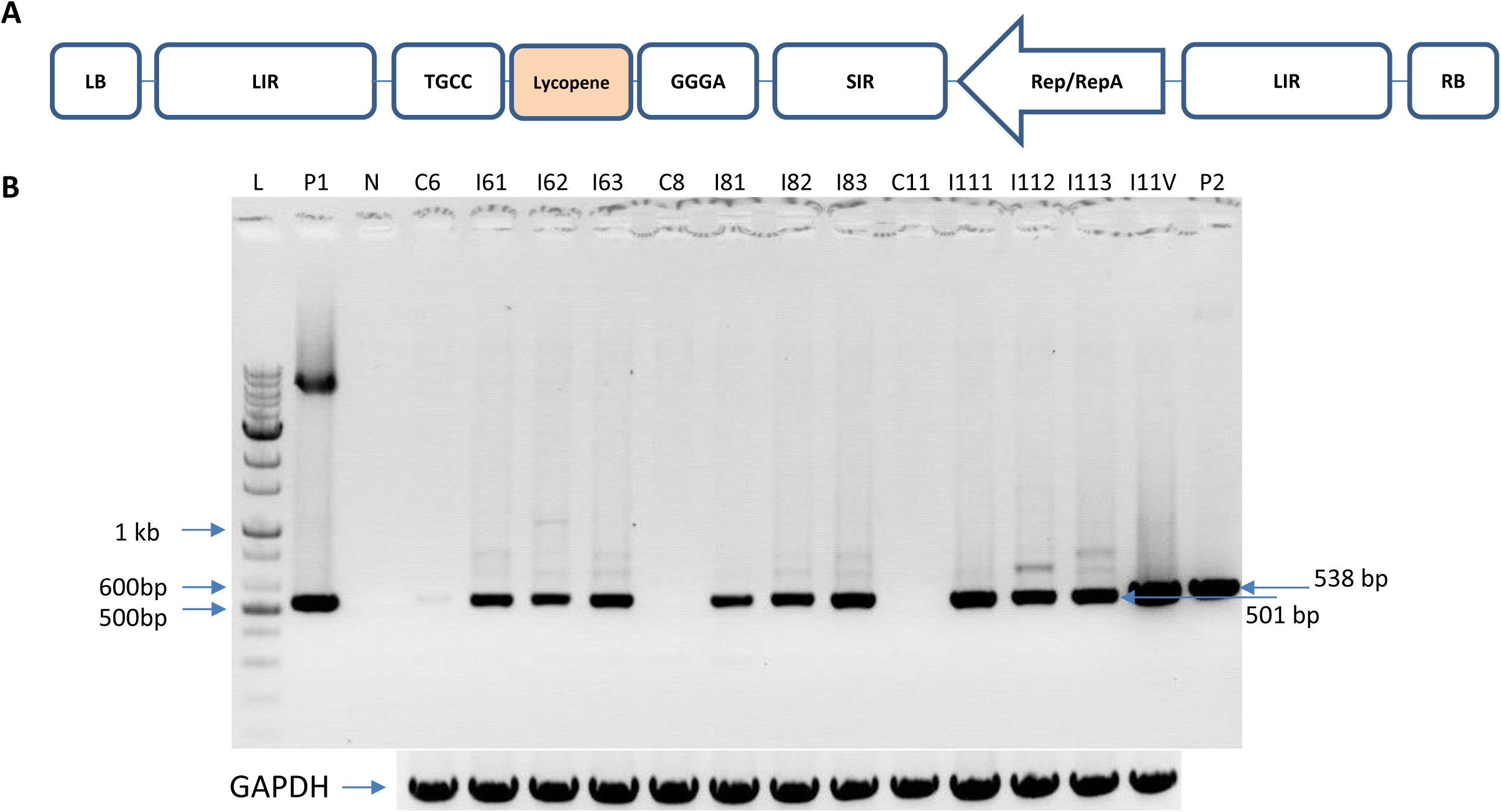
The *de novo-*engineered geminiviral amplicon (named pLSL.R. Ly) and its replication in tomato. (**A**) Map of pLSL.R.Ly. The DNA amplicon is defined by its boundary sequences (long intergenic region, LIR) and a terminated sequence (short intergenic region, SIR). The replication-associated protein (Rep/RepA) is expressed from the LIR promoter sequence. All of the expression cassettes of HDR tools were cloned into the vector by replacing the red marker (Lycopene) using a pair of type IIS restriction enzymes (BpiI, flanking ends are TGCC and GGGA). Left (LB) and right (RB) denote the borders of a T-DNA. (**B**) Circularized DNA detection in tomato leaves infiltrated with pLSL.R. Ly compared to those infiltrated with pLSLR. Agrobacteria containing the plasmids were infiltrated into tomato leaves (Hongkwang cultivar), and infiltrated leaves were collected at 6, 8 and 11 dpi and used for the detection of circularized DNAs. N: water; P1: positive control for pLSL.R. Ly; positive control for P2: pLSLR; Cx: Control samples collected at x dpi; Ixy: infiltrated sample number y collected at x dpi; I11 V: sample collected from leaves infiltrated with pLSLR at 11 dpi. PCR products obtained using primers specific to GAPDH were used as loading controls.

**Supplemental Figure 4.**
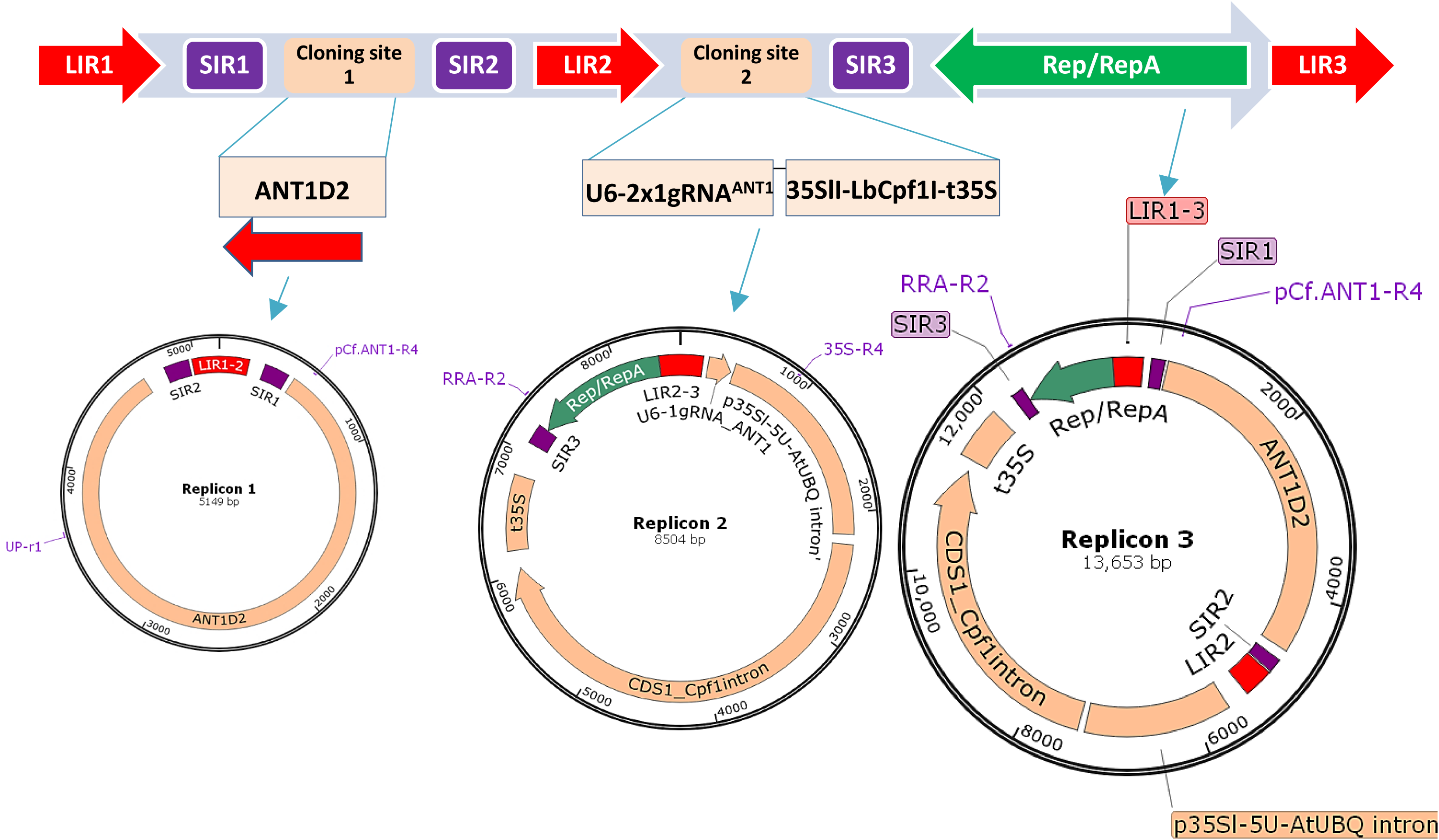
Schematic representation of the system and the released forms of the MR01 multi-replicon system. Upper panel: The design for the general construction of multi-replicon complexes included three LIR and three SIR sequences. Middle panel: Representative arrangement of the MR01 tool. The donor template was cloned in one replicon, and the other components for inducing DSBs were located the other replicon. Bottom panel: Three replicons would be formed from the MR01. Primer pairs for detecting circularized replicon 1 (Upr1/pCf. ANT1-R4), replicon 2 (RRA-R2/35S-R4), and replicon 3 (RRA-R2/pCf. ANT1-R4) are indicated in the map of each replicon.

**Supplemental Figure 5.**
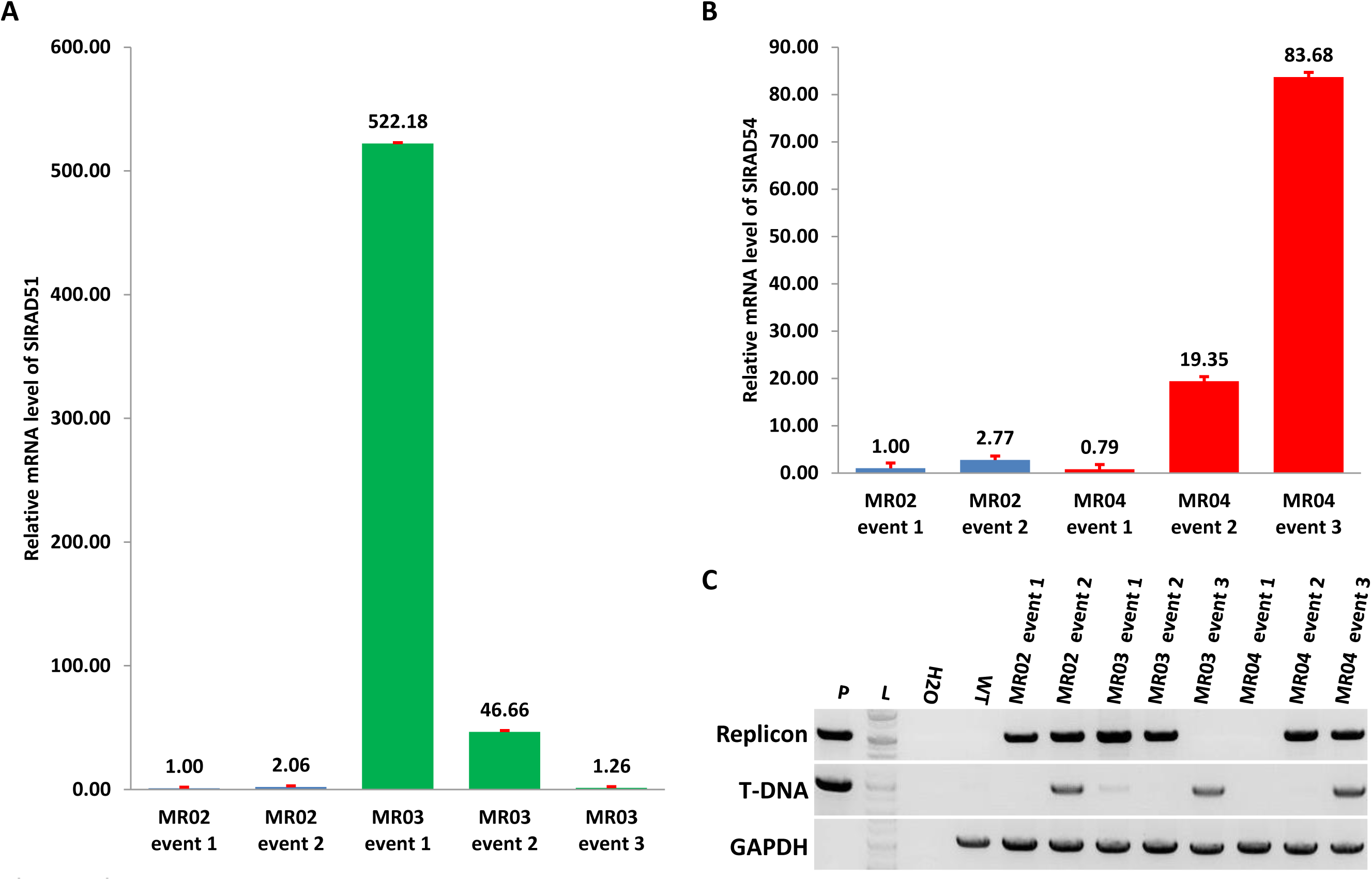
Relative levels of SlRAD51 or SlRAD54 transcripts expressed in transgenic events carrying MR03 or MR04, respectively. A. Relative mRNA levels of three independent events (event 1, 2 and 3) obtained from MR03 transformation. Event 3 was shown to be free of replicons in (C). B. Relative mRNA levels of three independent events (event 1, 2 and 3) obtained from MR04 transformation. Event 1 was shown to be free of replicons in (C). C. PCR analysis of replicon presence in the transformed events. P: plasmid MR02; L: BIOFACT 1kb ladder plus; H2O: water; WT: wild-type Hongkwang; MR02 event 1 and 2: transformed plants obtained using MR02 vector; MR03 event 1, 2 and 3: transformed plants obtained using MR03 vector; MR04 event 1, 2 and 3: transformed plants obtained using MR04 vector. MR03 event 3 and MR04 event 1 appear to be non-transformed events which might escaped from antibiotic selection. The error bars shown in (A) and (B) are standard deviation values.

**Supplemental Figure 6.**
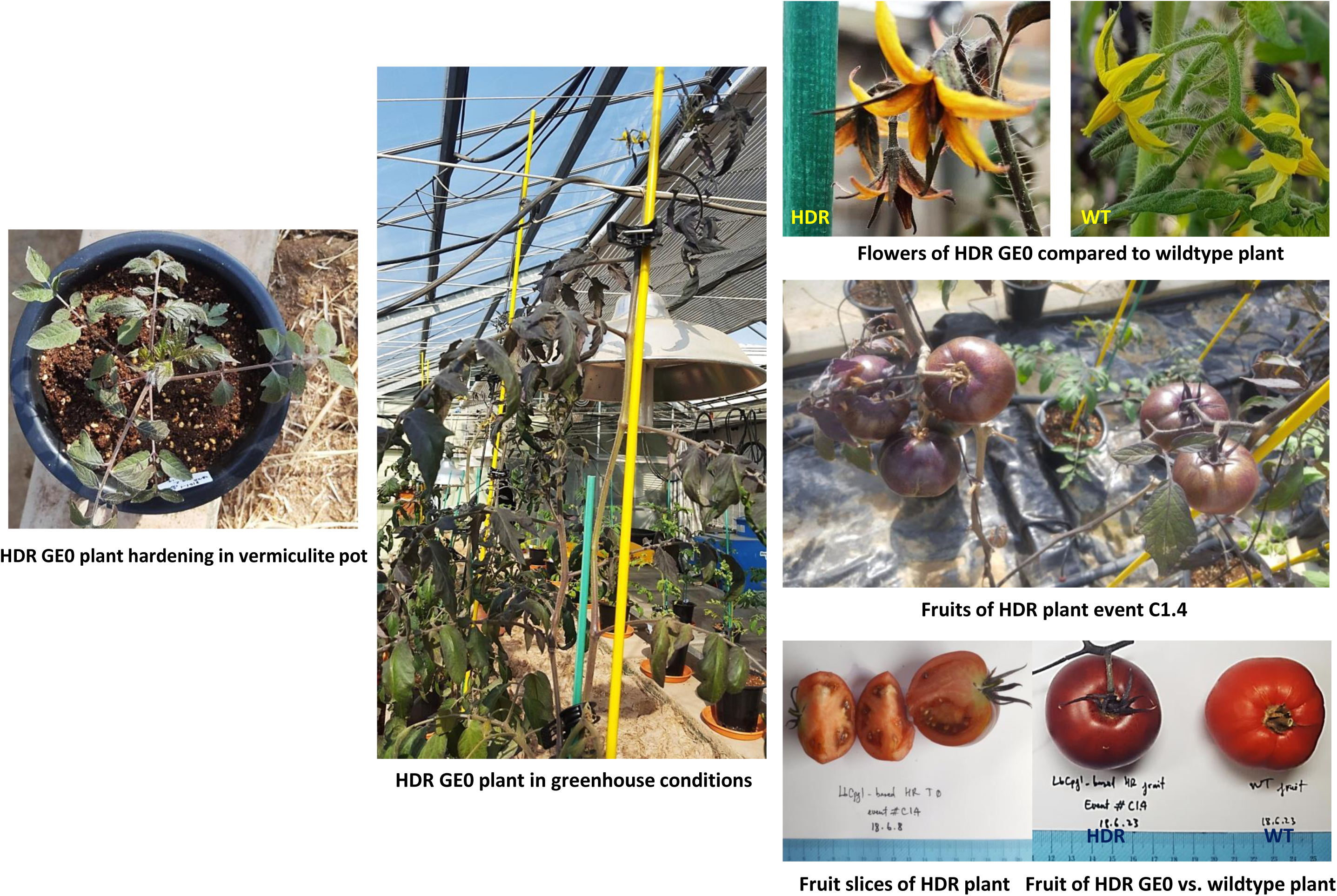
Morphological appearance of GE0 plants

**Supplemental Figure 7.**
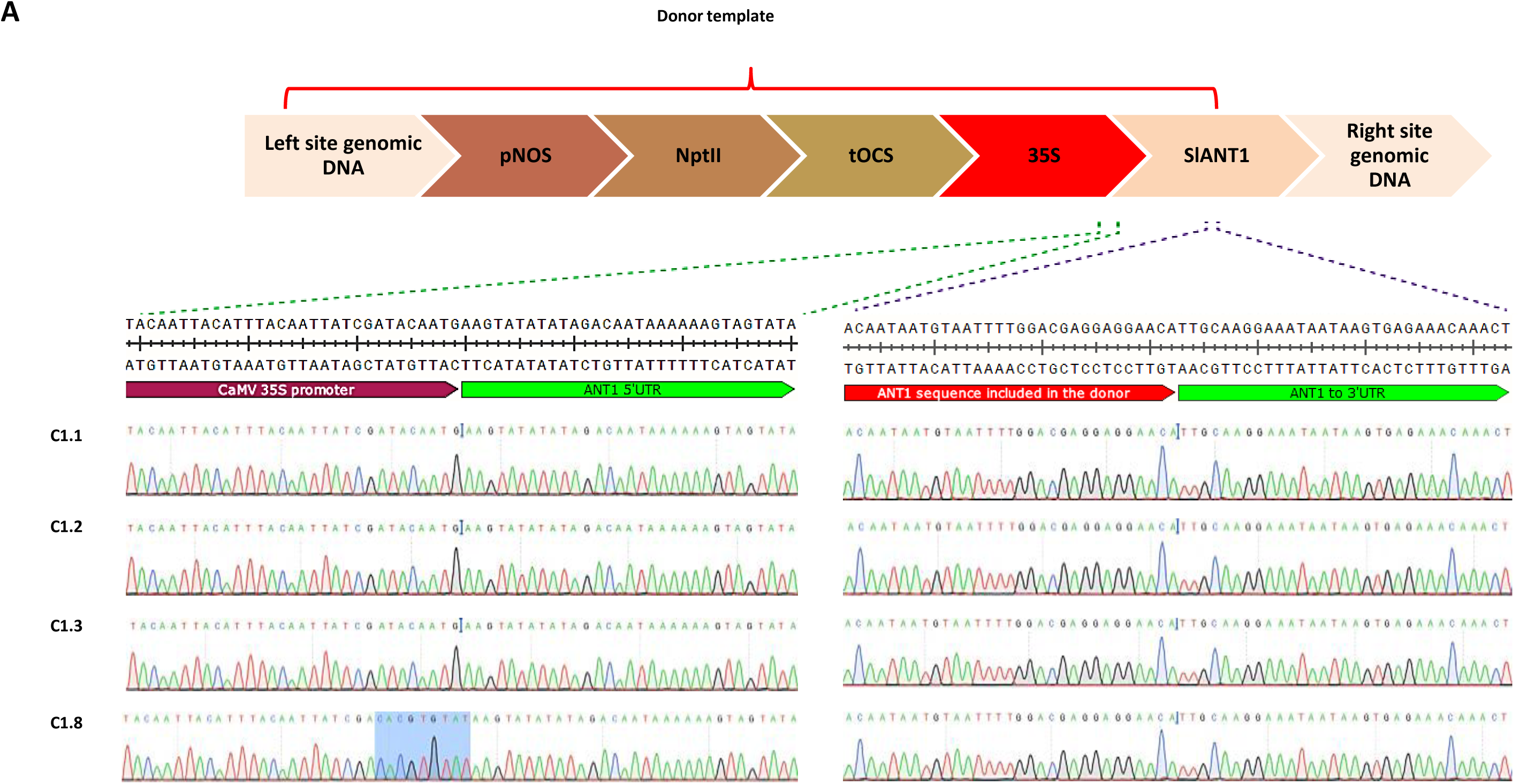
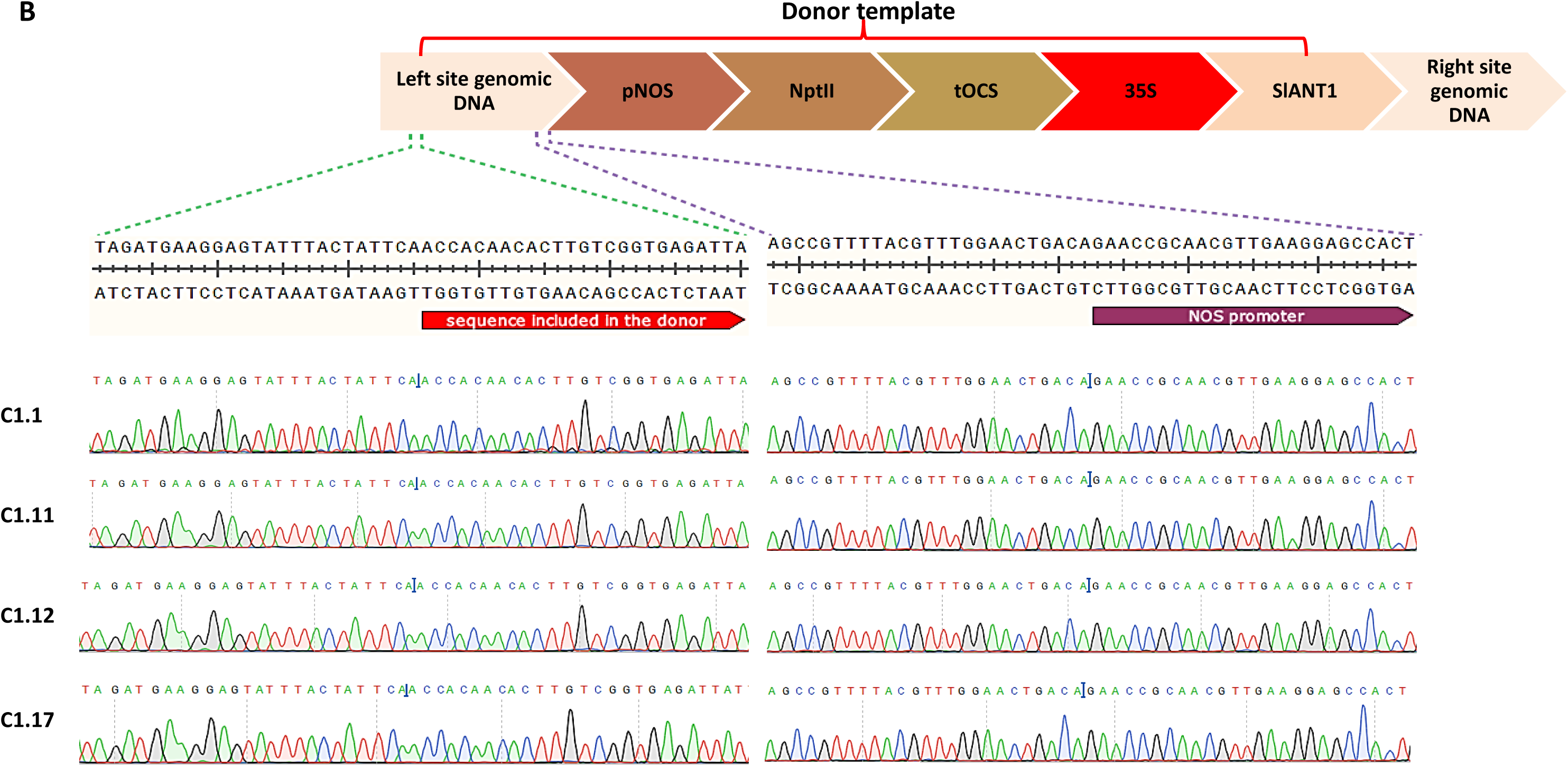
Sanger sequencing data to confirm donor exchanges. (**A**) Right junction. (**B**) Left junction. C1.1, C1.2, C1.3, C1.8, C1.11, C1.12, and C1.17: Independent LbCpf1-based HDR GE0 events

**Supplemental Figure 8.**
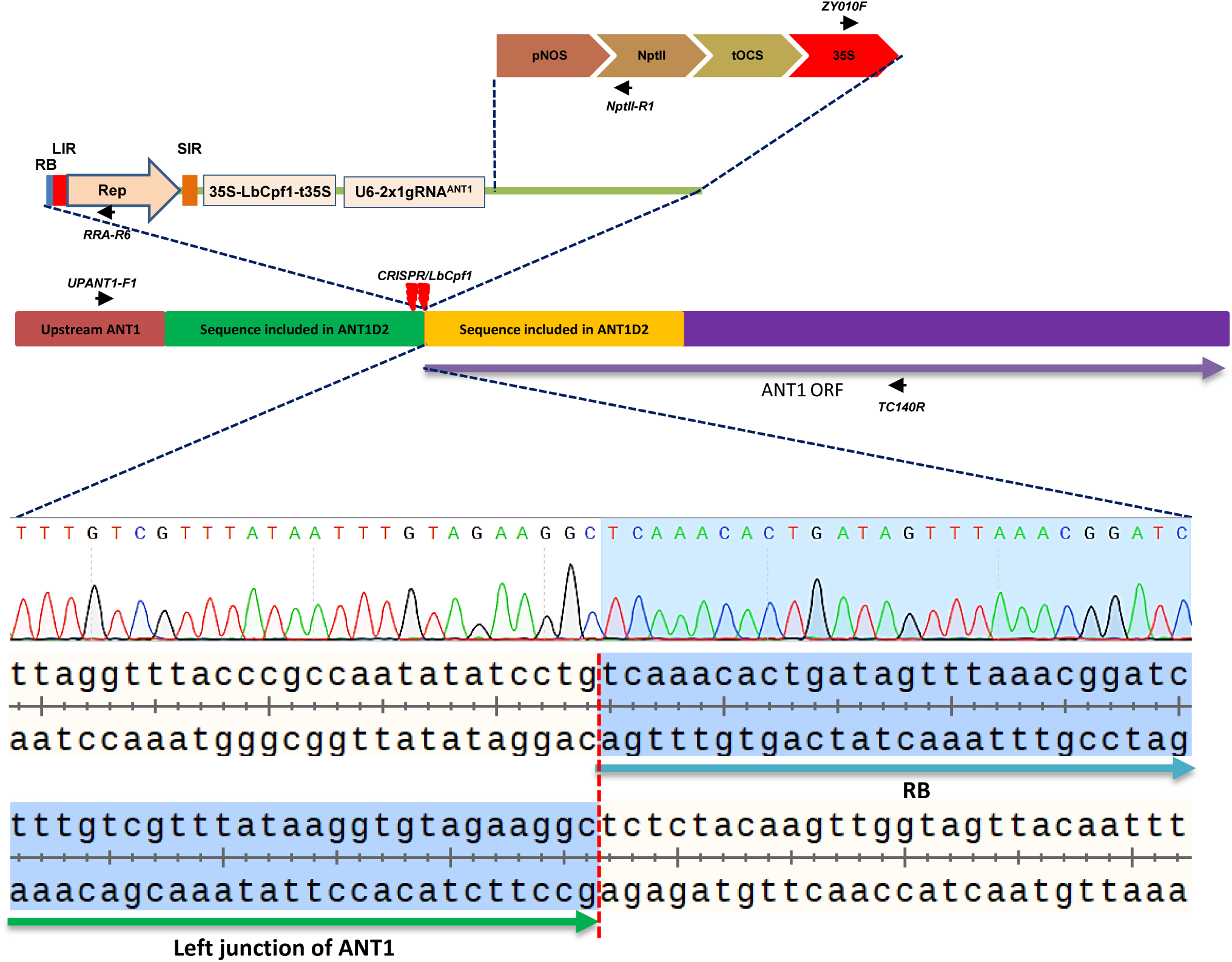
Error-prone repair combining HDR and NHEJ in event #C1.3. The right junctions (amplified by ZY010F/TC140R) of the events were confirmed to be perfectly adapted to HDR repair (Supplemental Figure 6A), but the left junction could not be amplified (using the UPANT1-F1/NptII-R1 primer pair, Figure 1A). Sequencing of the left junction region showed a ligation event between the RB of the T-DNA and the 3’ break in the upstream ANT1 promoter sequence via NHEJ. Red dotted line: ligation boundary.

**Supplemental Figure 9.**
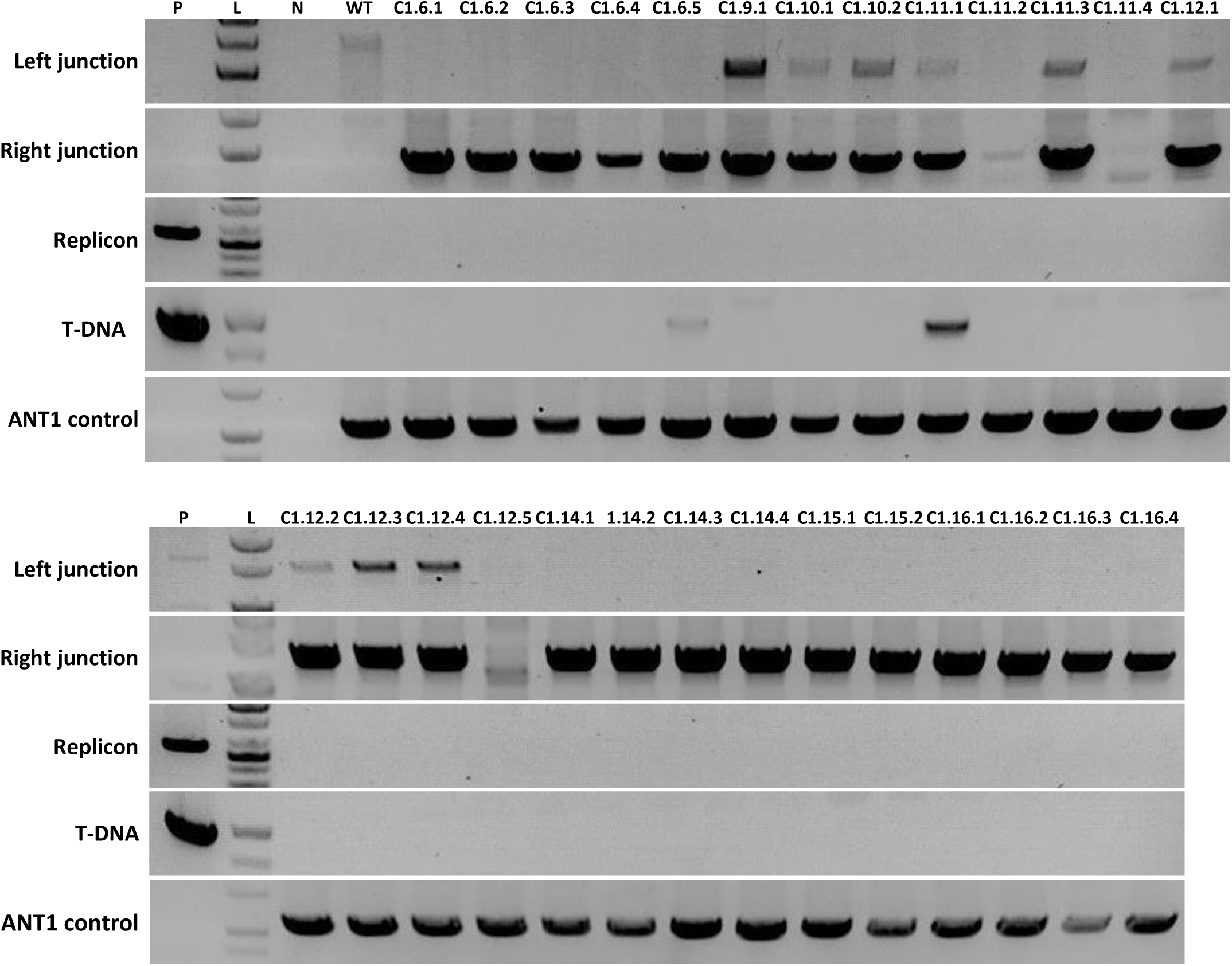
PCR analyses of GE1 plants obtained from GE0 LbCpf1-based HR events. P: pHR01 plasmid isolated from Agrobacteria; L: 1 kb ladder; N: water control; WT: wild-type Hongkwang; C1.6.1-C1.6.5: GE1 offspring of event C1.6.; C1.9.1: GE1 offspring of event C1.9; C1.10.1 and C1.10.2: GE1 offspring of event C1.10; C1.11.1-C1.11.4: GE1 offspring of event C1.11; C1.12.1-C1.12.5: GE1 offspring of event C1.12; C1.14.1-C1.14.4: GE1 offspring of event C1.14; C1.15.1 and C1.15.2: GE1 offspring of event C1.15; C1.16.1-C1.16.4: GE1 offspring of event C1.16.

**Supplemental Figure 10.**
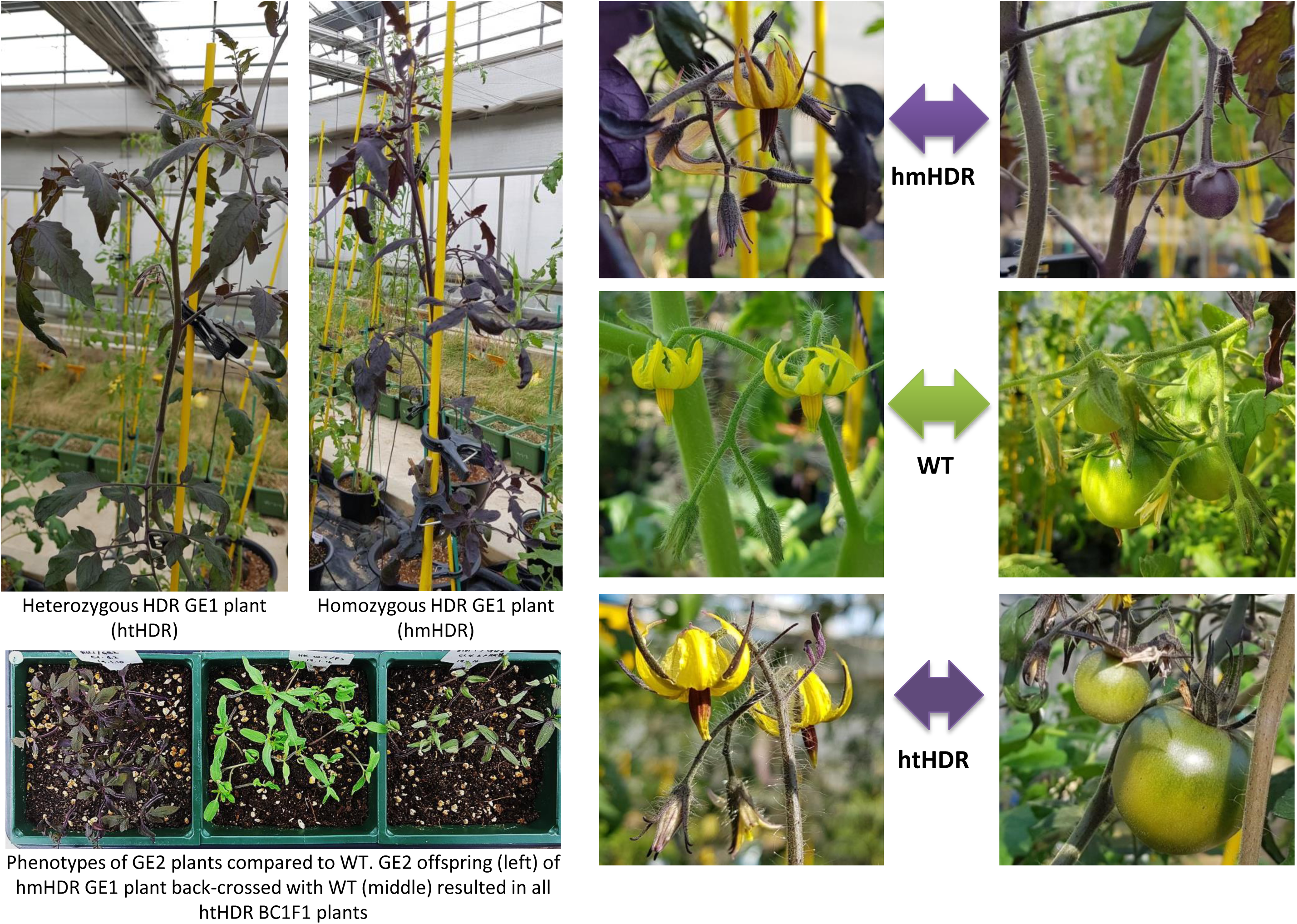
Morphological appearance of GE1 plants

**Supplemental Figure 11.**
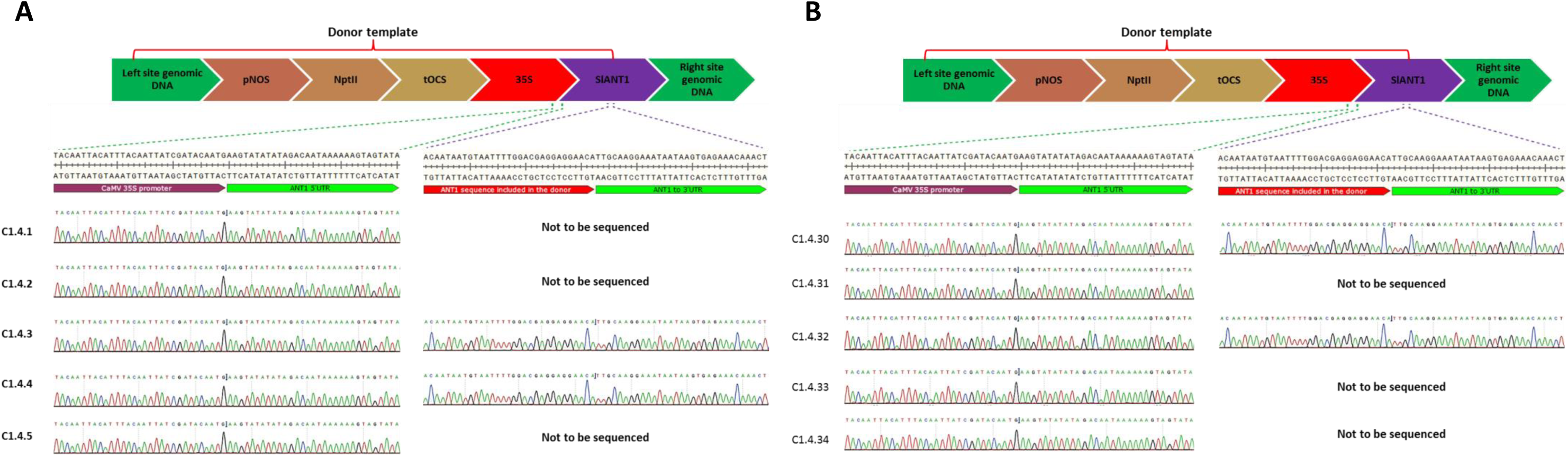
Analyses of left and right junction sequences of GE1 plants. Sanger sequencing data to confirm donor exchanges for the right (A) and left (B) junctions of the GE1 plants are presented.

**Supplemental Figure 12.**
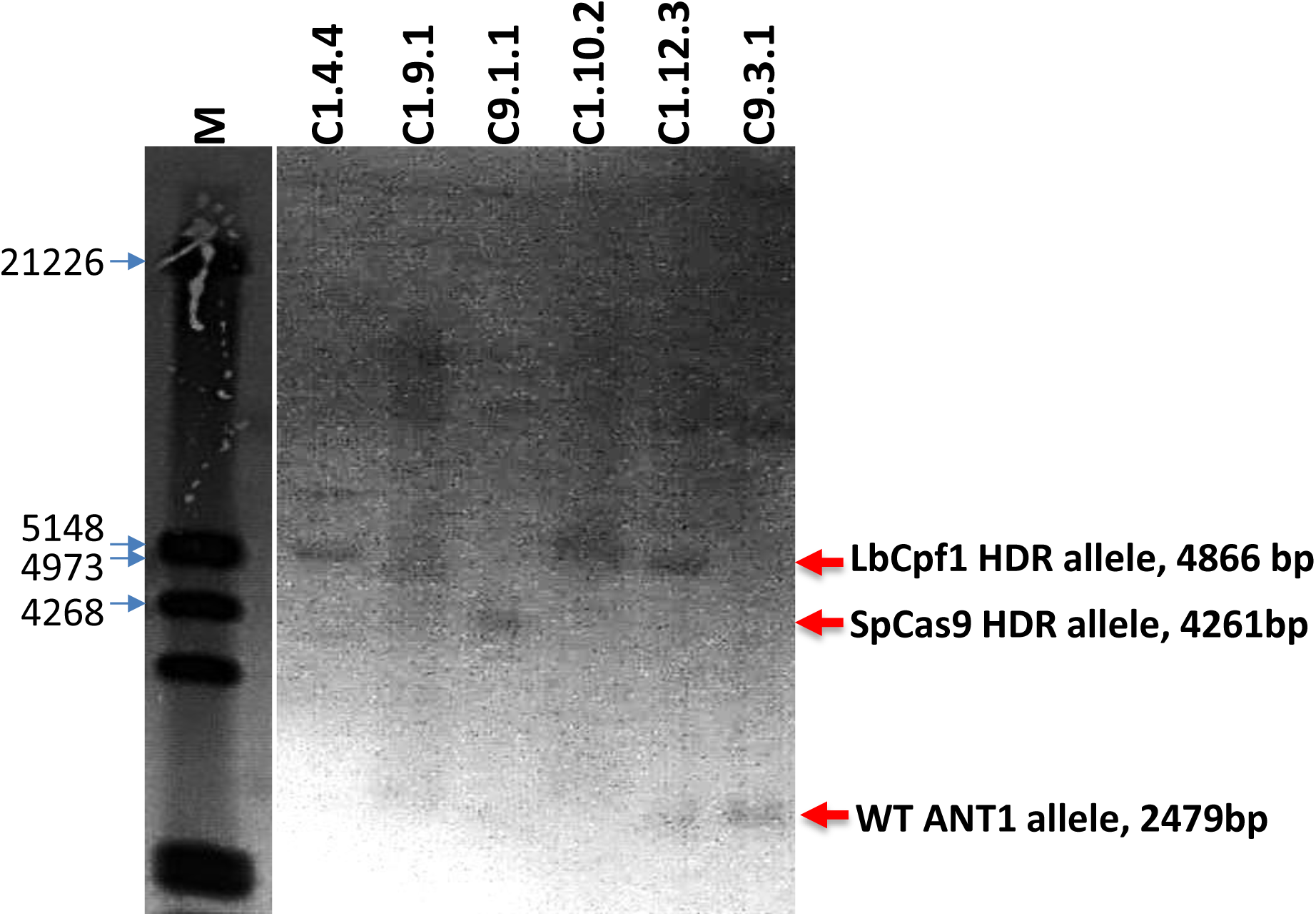
Southern Blot analysis of the ANT1 edited locus. Genomic DNAs were isolated from the leaves of the ANT1 HDR events and 20 µg of each gDNAs were digested with NsiI and resolved on 0.8% agrose gel. The resolved DNAs were then blotted onto Hybond N+ membrane and later detected using Dioxigenin labeled Probe (Supplemental Figure 2). The marker lane (M, DNA Molecular Weight Marker III, Digoxigenin-labeled, Roche) is separated for better illustration of the image. The expected bands with sizes are denoted beside the Southern blot panel. C1.4.4, C1.9.1, C1.10.2, C1.12.3: GE1 lines showed strong amplification of RJ and LJ in Supplemental Figure 9; C9.1.1 (RJ and LJ edited) and C9.3.1 (only RJ inserted): two GE1 line obtained by pTC217 tool.

**Supplemental Figure 13.**
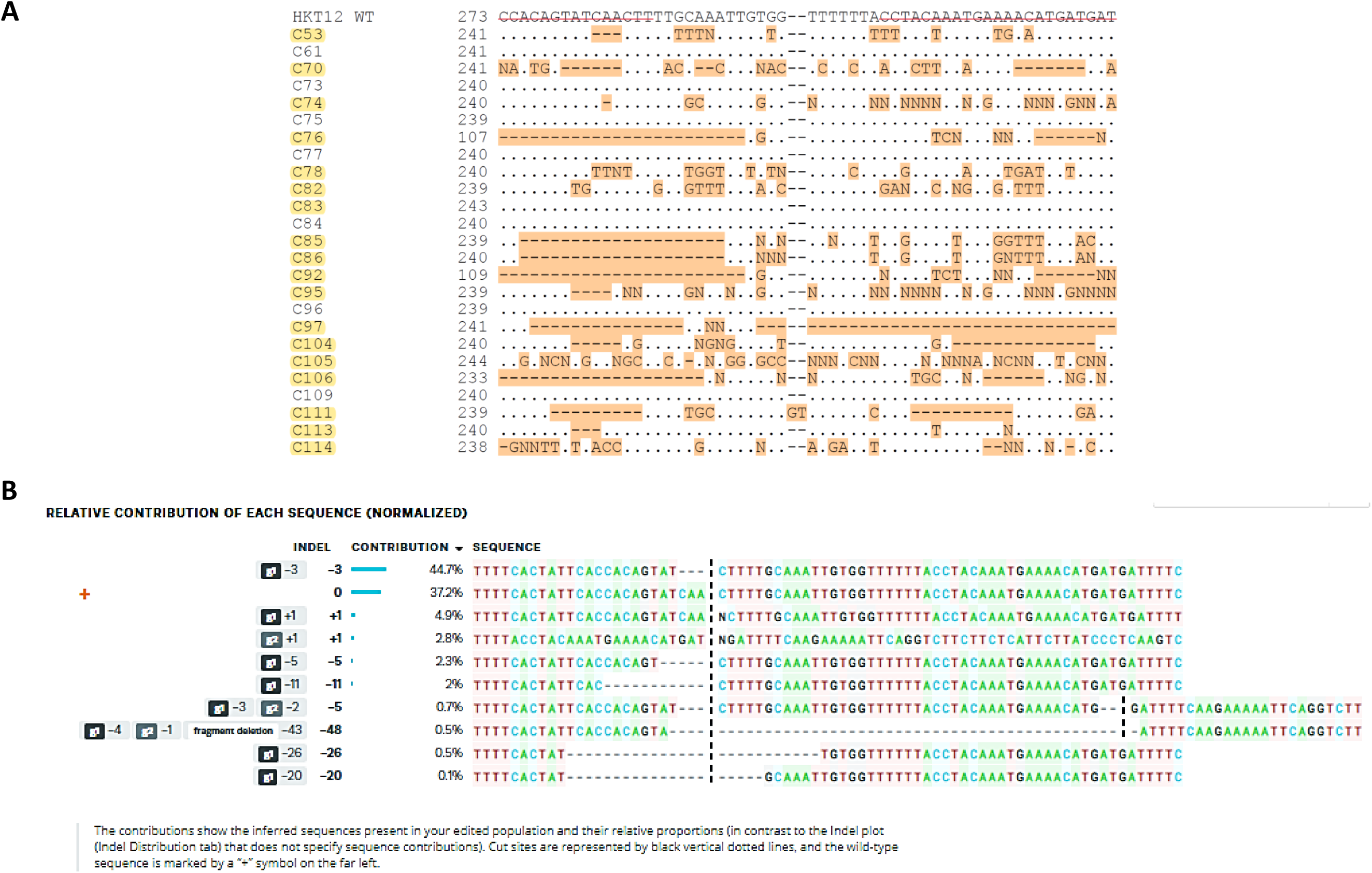
Analyses of indel mutations in HKT12 events. (A) Alignment of raw sequences obtained from Sanger sequencing. 18/25 events (highlighted in yellow) showed strong double peaks indicating single/biallelic mutations. Six out of 25 events showed clear biallelic mutations. C77 showed weak (30%) double peaks. C83 and C105 showed large truncations. (B) Decomposed sequence of event #C53 obtained with ICE Synthego software.

**Supplemental Figure 14.**
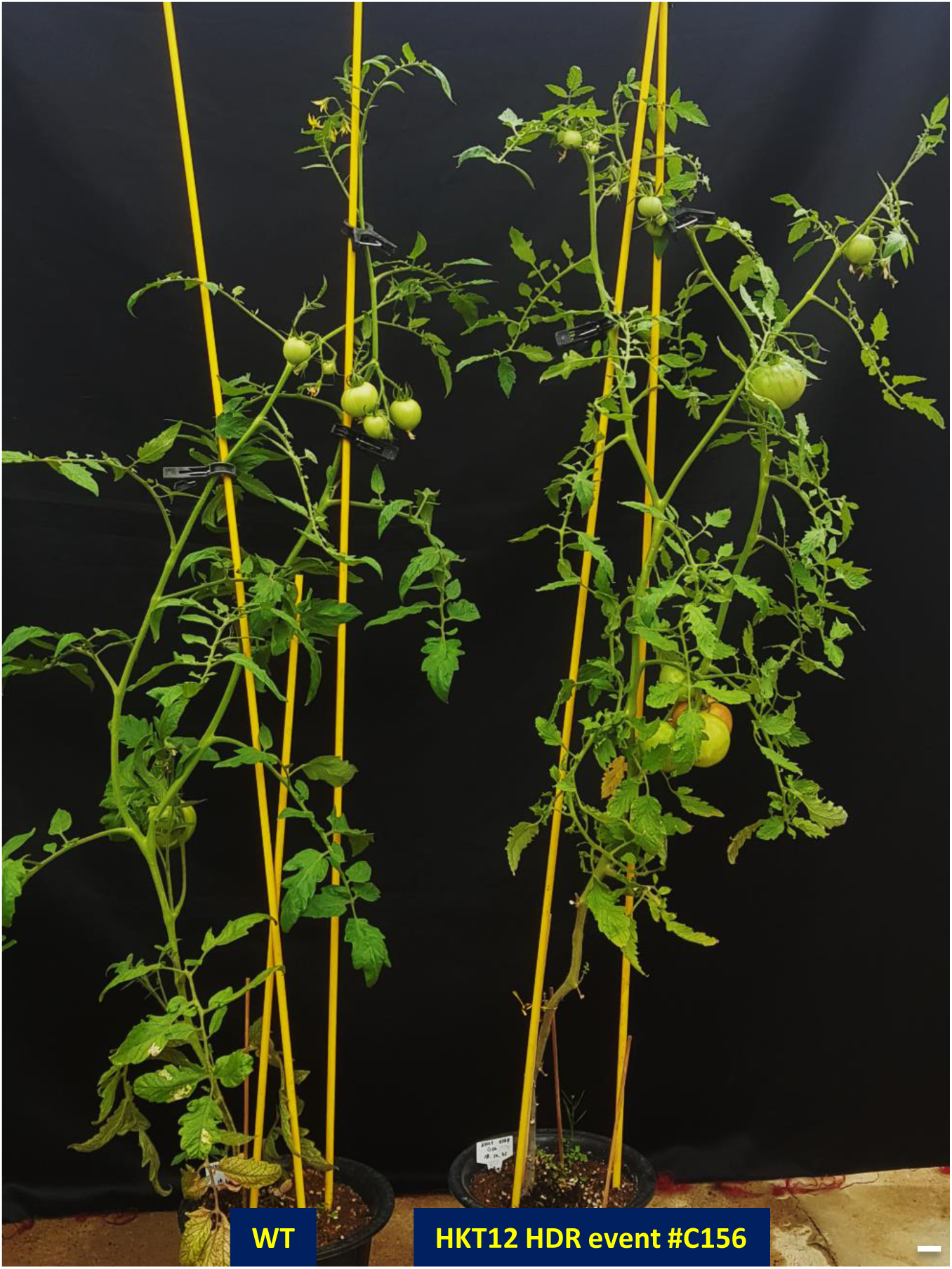
Morphology of the heterozygous HKT12^N217D^ event in a mature stage. A plant resulting from the HKT12^N217D^ event (right) shows a normal morphology and fruit setting compared to the parental plant (left). Scale bars = 2 cm.

**Supplemental Figure 15.**
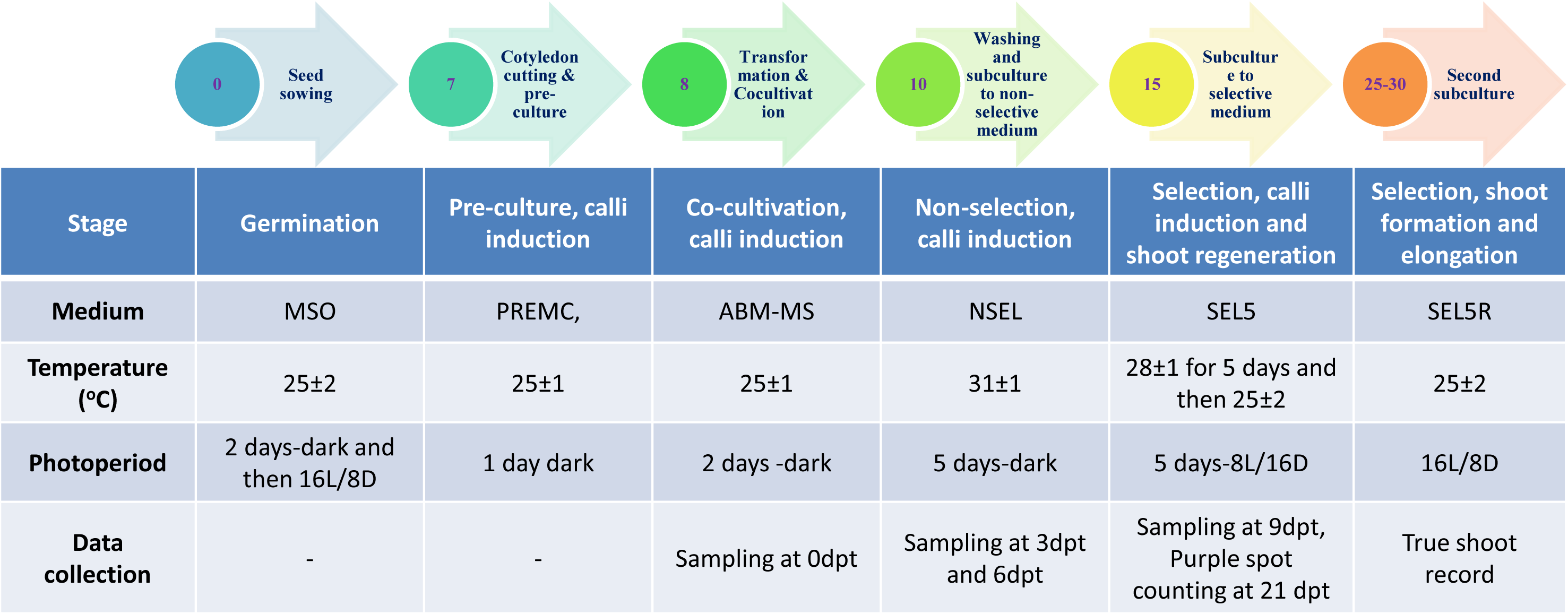
Timeline and contents of the *Agro*-mediated transformation protocol used in this work. The step-by-step protocol is presented with each number in the circles indicating the number of days after seed sowing (upper panel), and the treatments used in each step are shown in the lower panel.

**Supplemental Table 1A.**
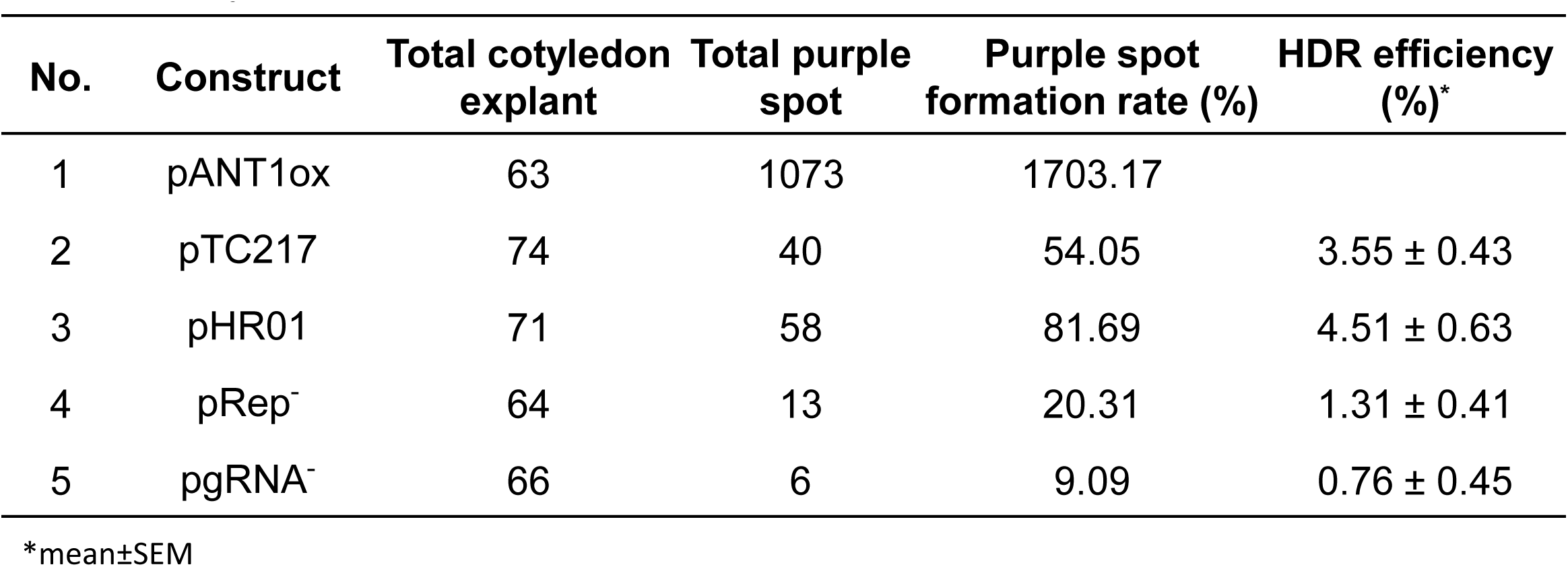
Purple spot data collected in the experiment for comparison of HDR efficiency between different constructs

**Supplemental Table 1B.**
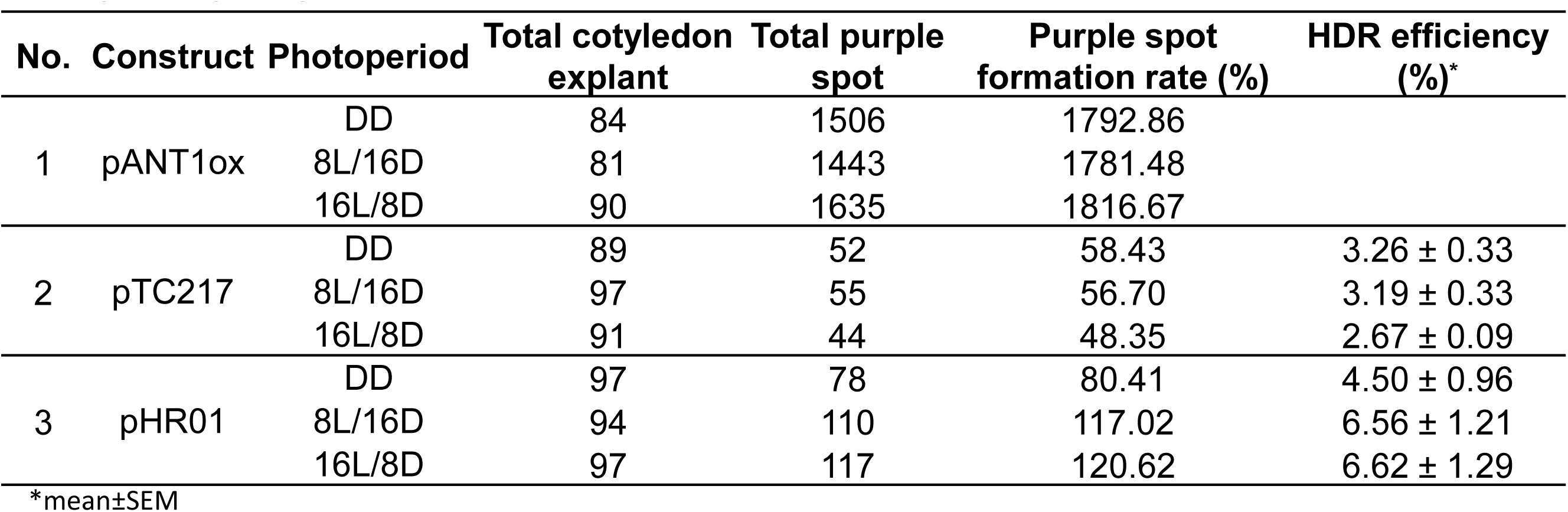
Purple spot data collected in the experiment for assessment of Impact of photoperiod on HDR

**Supplemental Table 1C.**
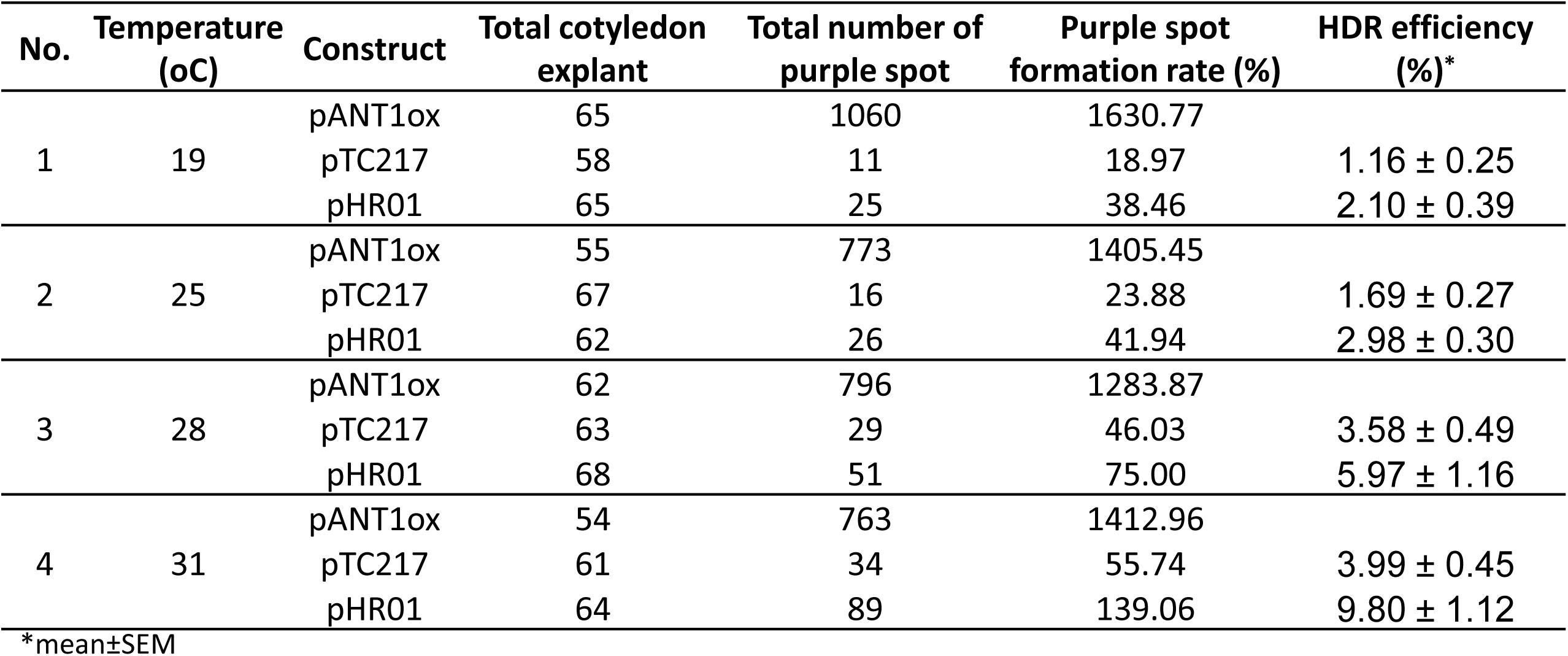
Purple spot data collected in the experiment for assessment of Impact of photoperiod on HDR

**Supplemental Table 2A.**
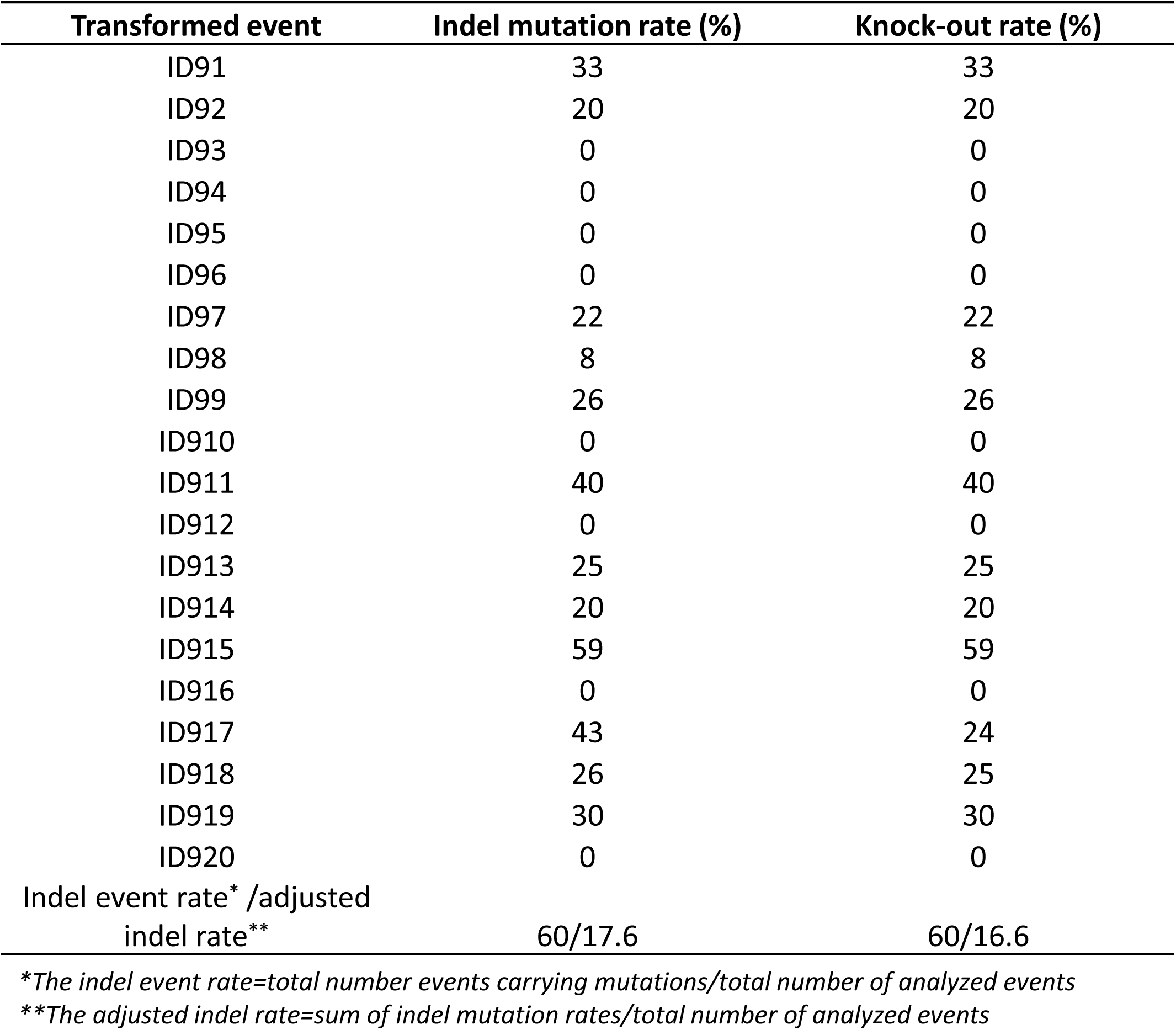
SlANT1 locus mutation rates observed from transformed events of pTC217

**Supplemental Table 2B.**
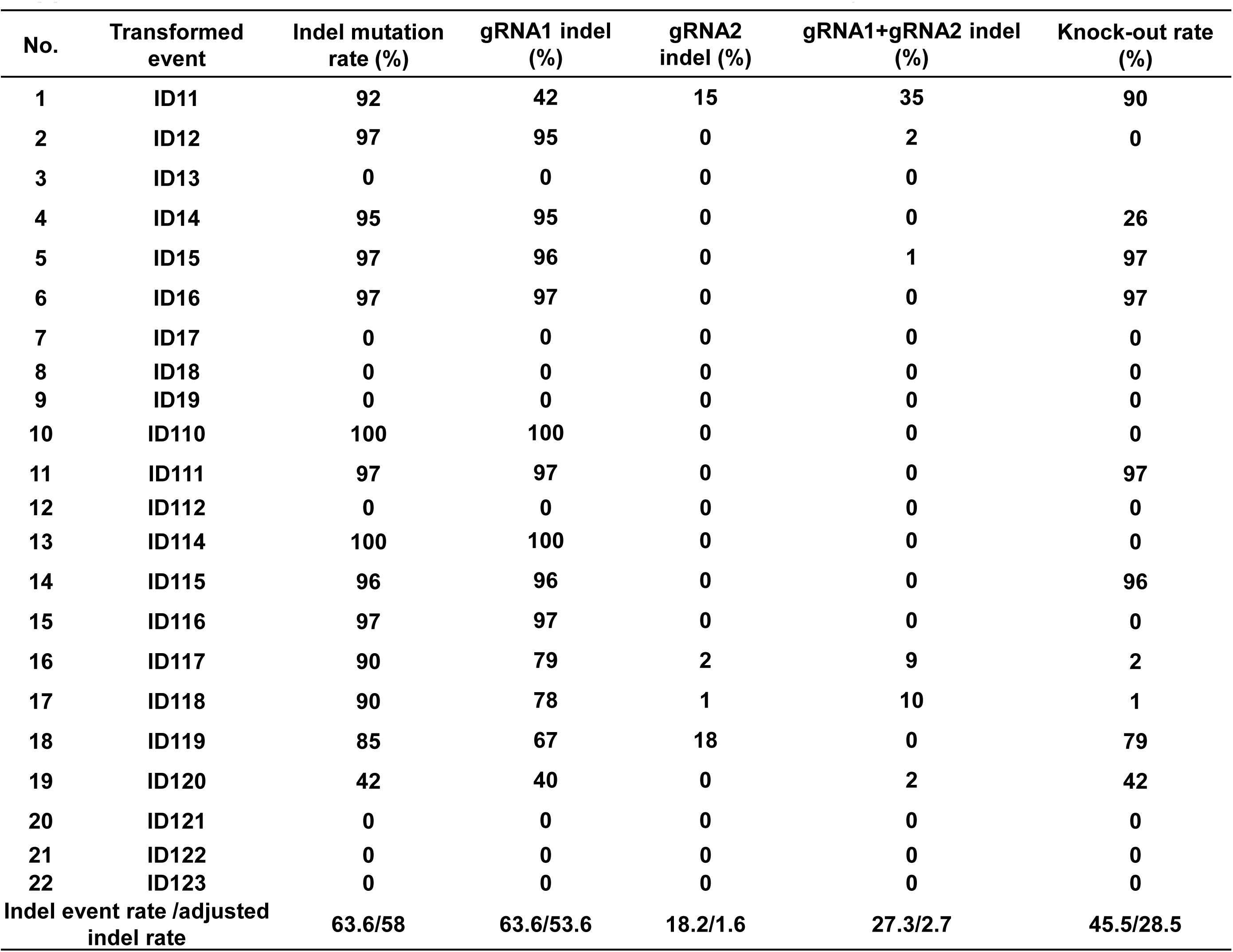
Indel mutation rates observed from transformed pHR01 events at SlANT1 sites

**Supplemental Table 3.**
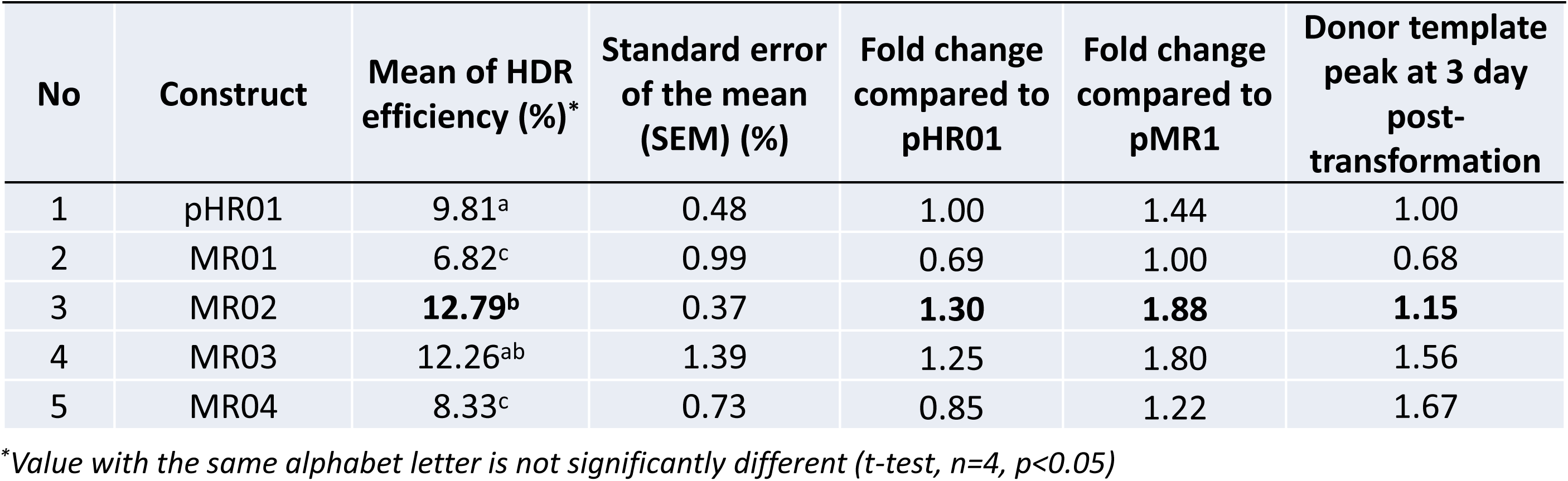
The increase in HDR by multi-replicon systems

**Supplemental Table 4.**
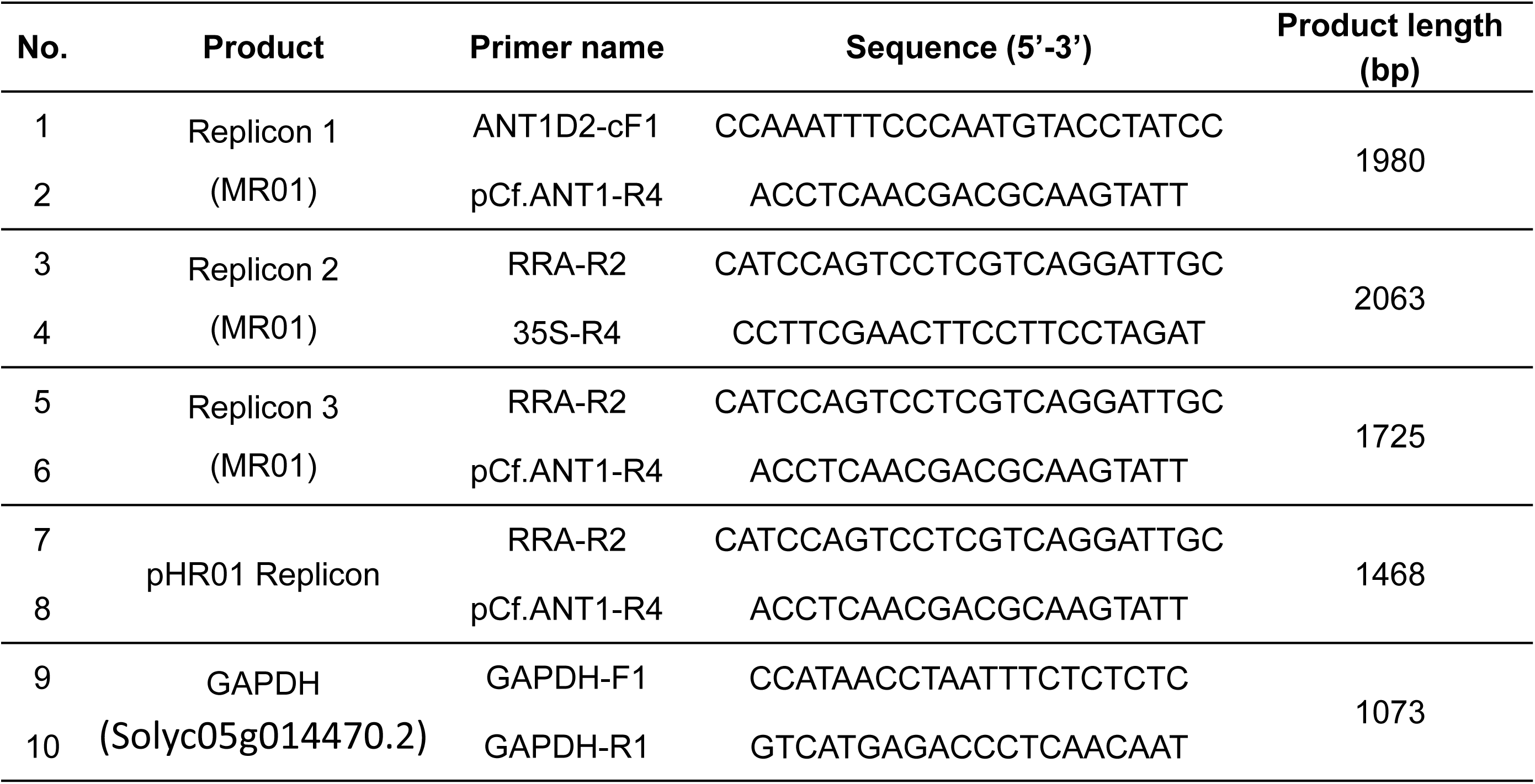
Primers for detecting circularized replicons released by MR01 and pHR01

**Supplemental Table 5.**
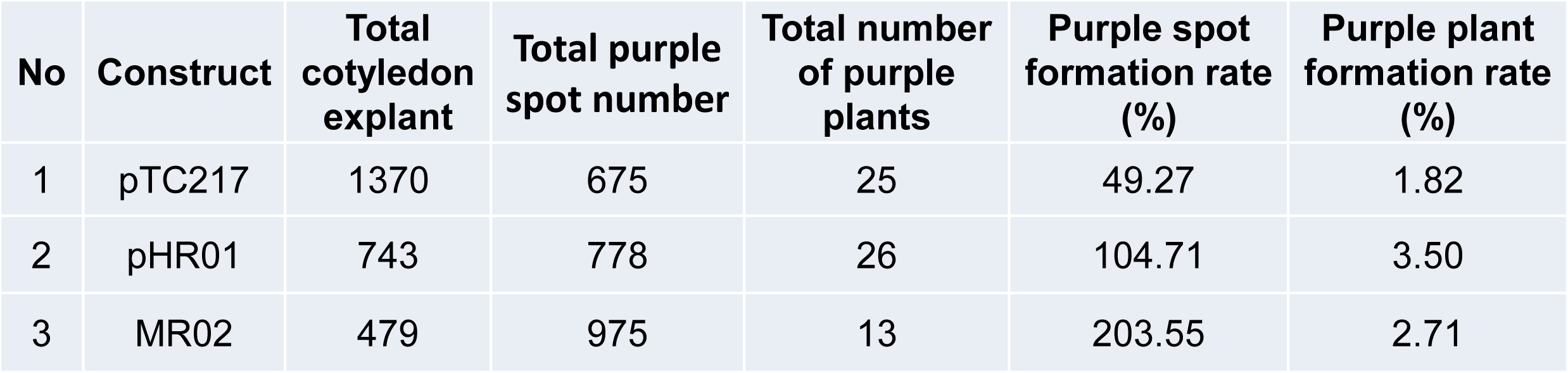
ANT1 HDR events derived from three main HDR constructs used in the study

**Supplemental Table 6.**
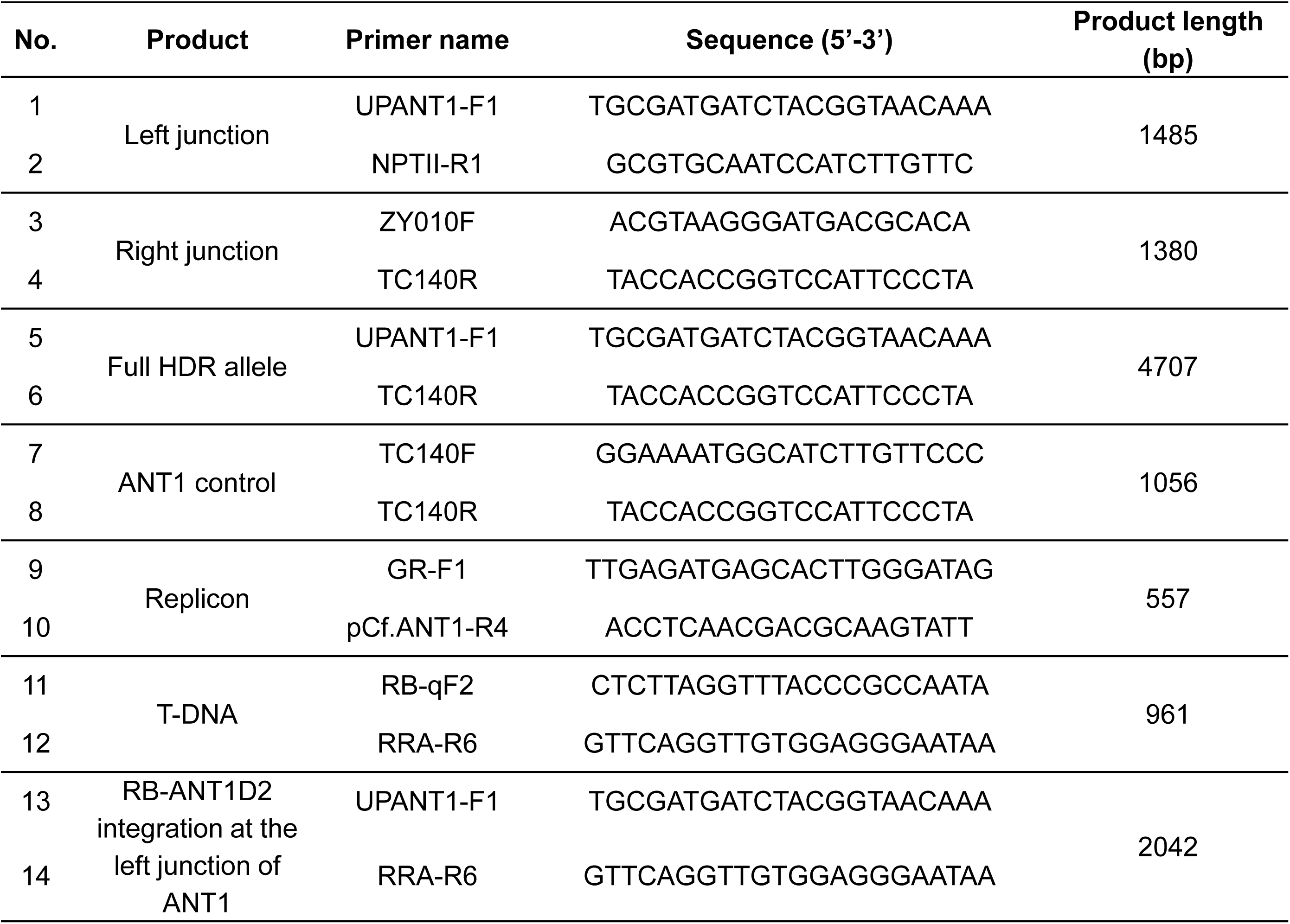
Primers for LbCpf1-based HR event analyses

**Supplemental Table 7.**
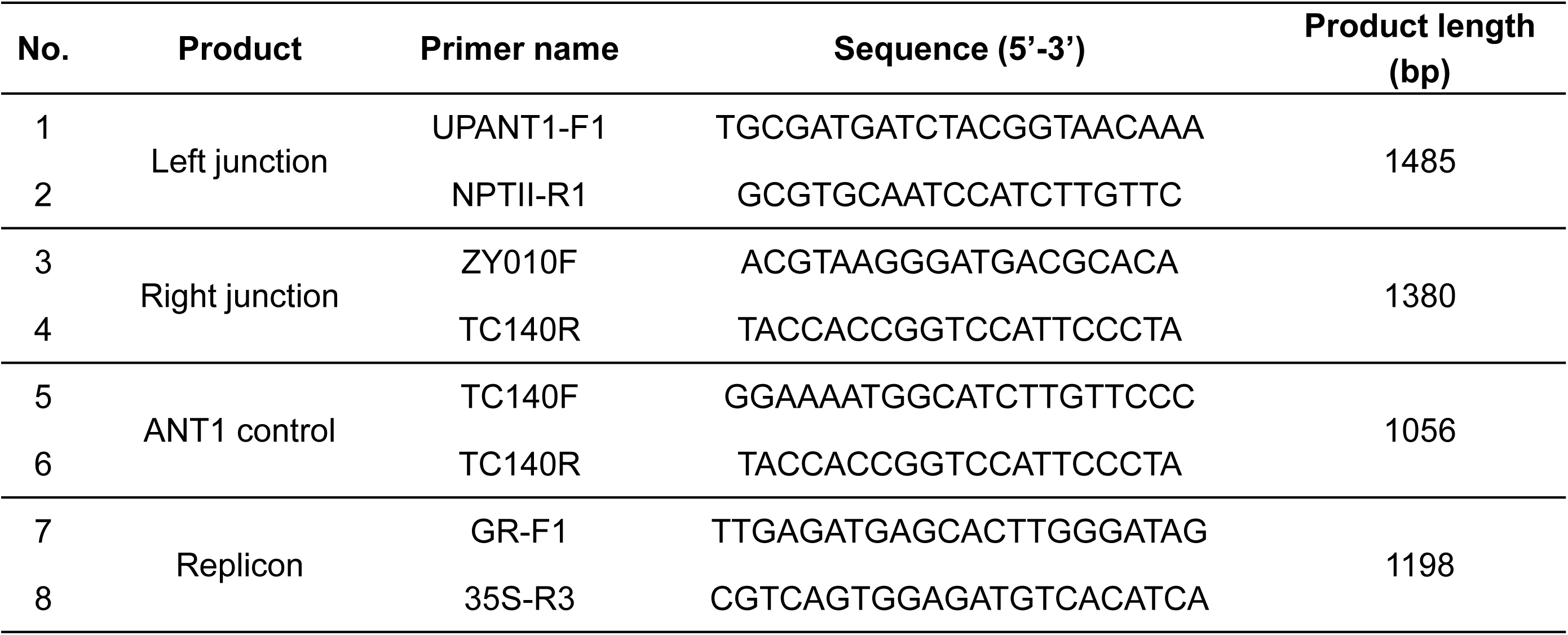
Primers for SpCas9-based HR event analyses

**Supplemental Table 8.**
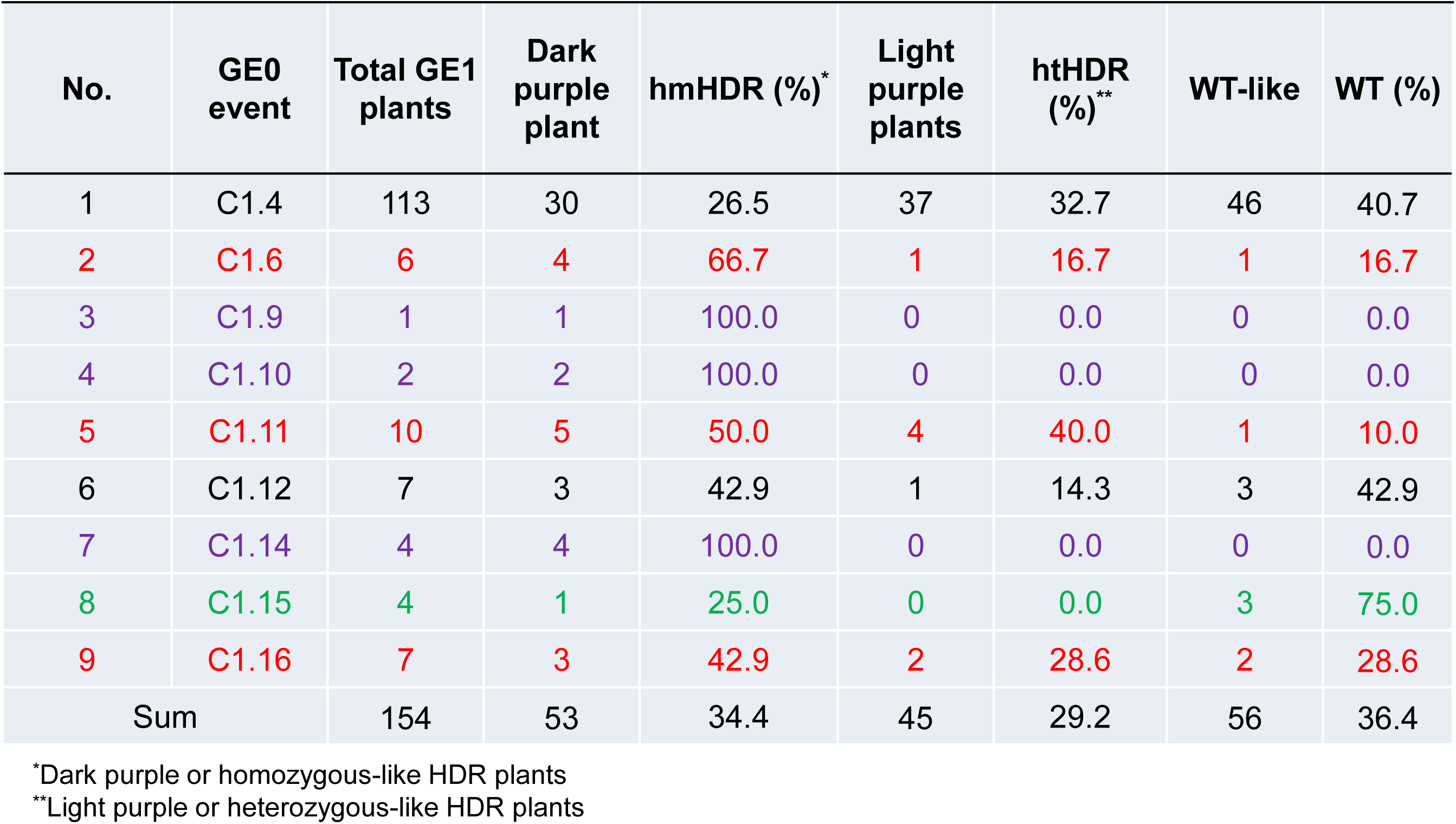
Phenotypic segregation of self-pollinated offspring resulting from LbCpf1-based HDR events.

**Supplemental Table 9.**
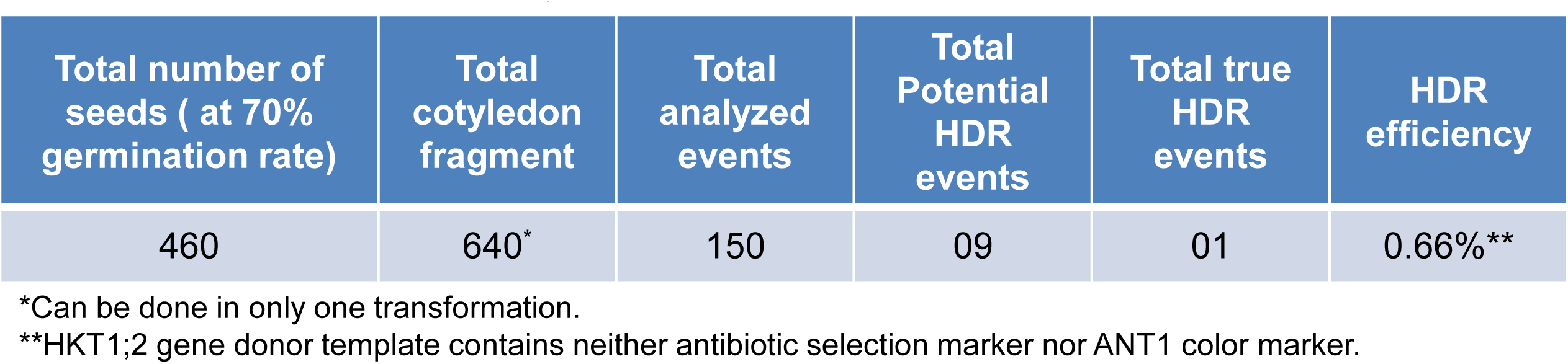
Summary of the *SlHKT1;2* HDR experiment.

**Supplemental Table 10.**
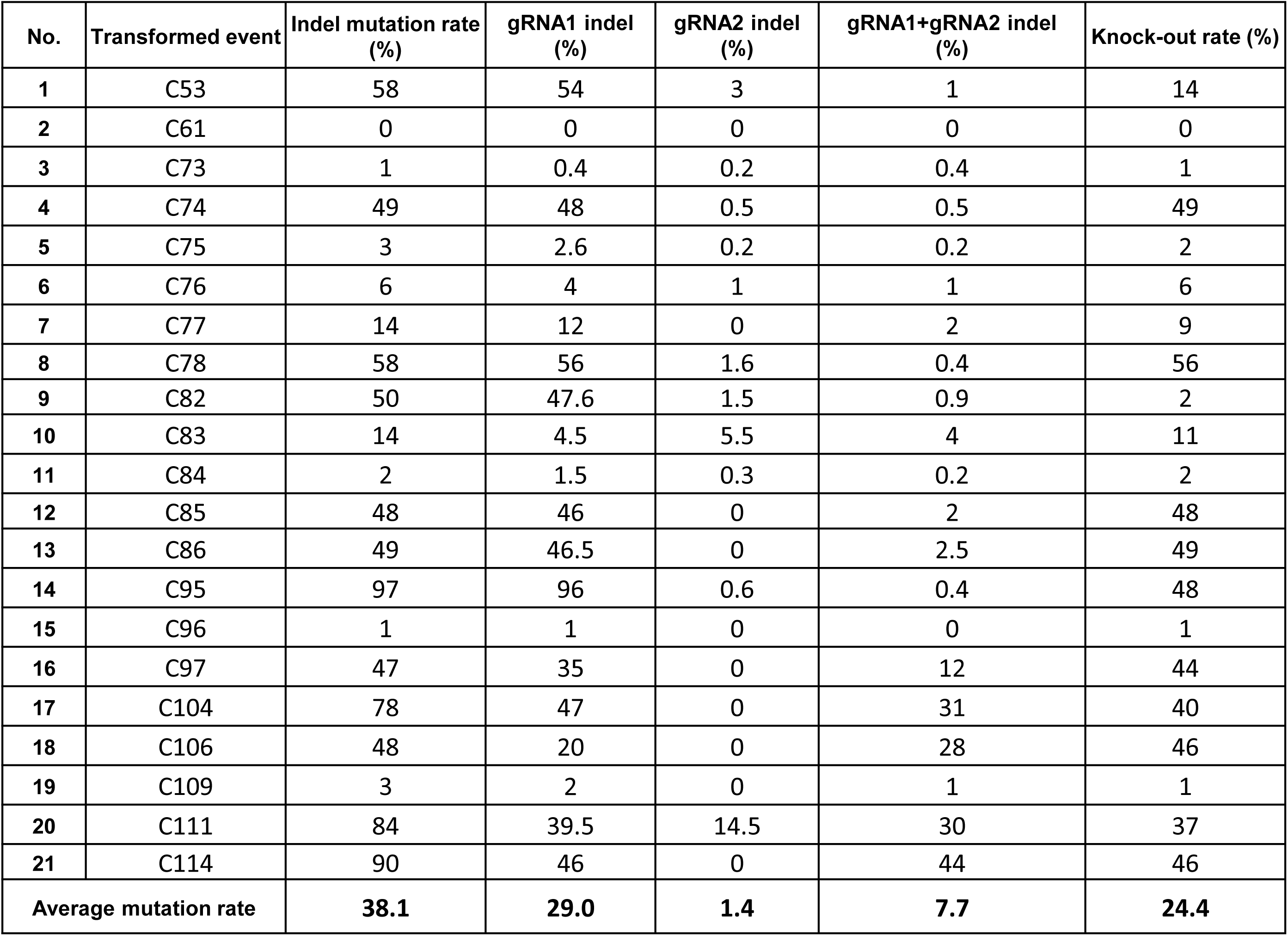
Indel mutation rates among *HKT12* samples decomposed by ICE Synthego software

## LIST OF SUPPLEMENTAL FILES

Data S1_sequences used in the study.

Data S2_Analysis of guide RNA activity.

Data S3_Potential off-targets of LbCpf1_gRNA1.

Data S4_qRT-PCR analyses of SlRAD51 and SlRAD54 mRNA levels.

Data S5_Southern blot analysis of GE1 plants.

## SEQUENCES USED IN THE STUDY

### Single replicon system with Golden gate level 2 acceptor sites

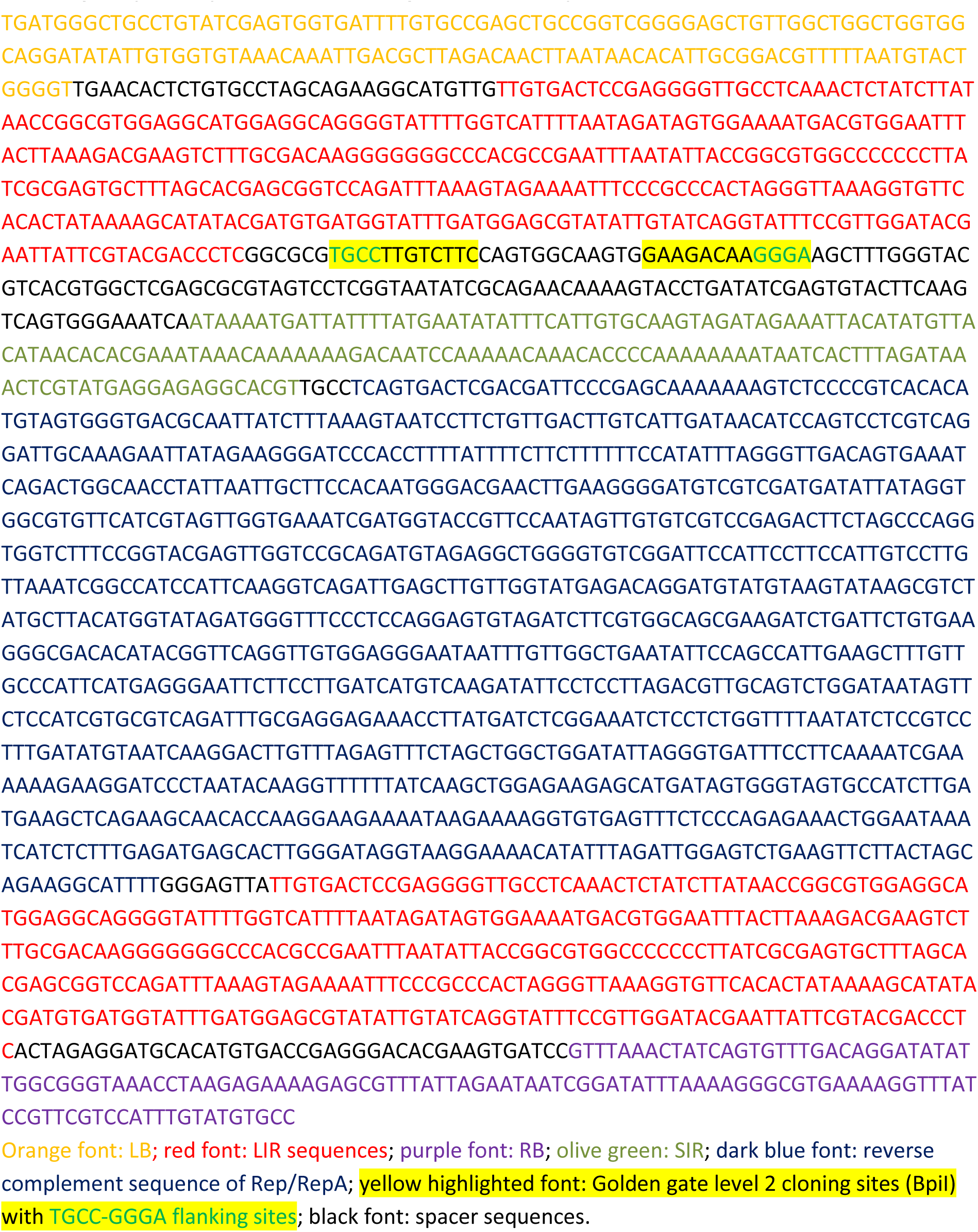

### In pHR01

#### The dual crRNA expression cassette

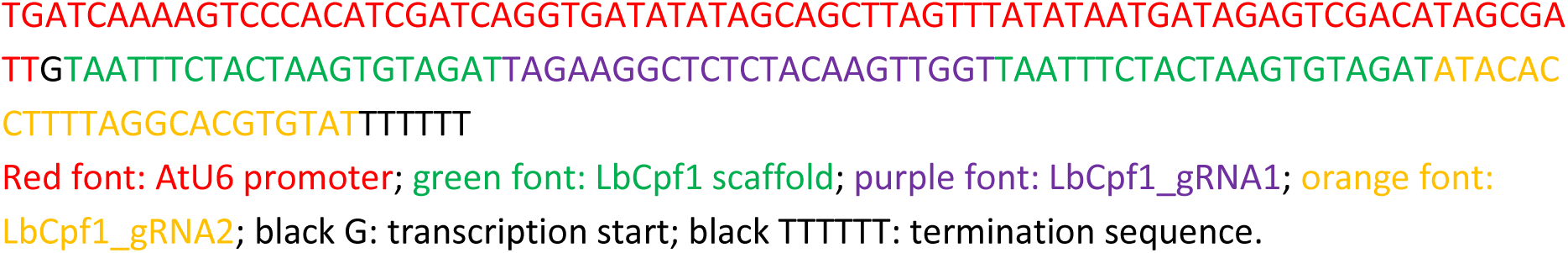

#### ANT1D2 donor sequence

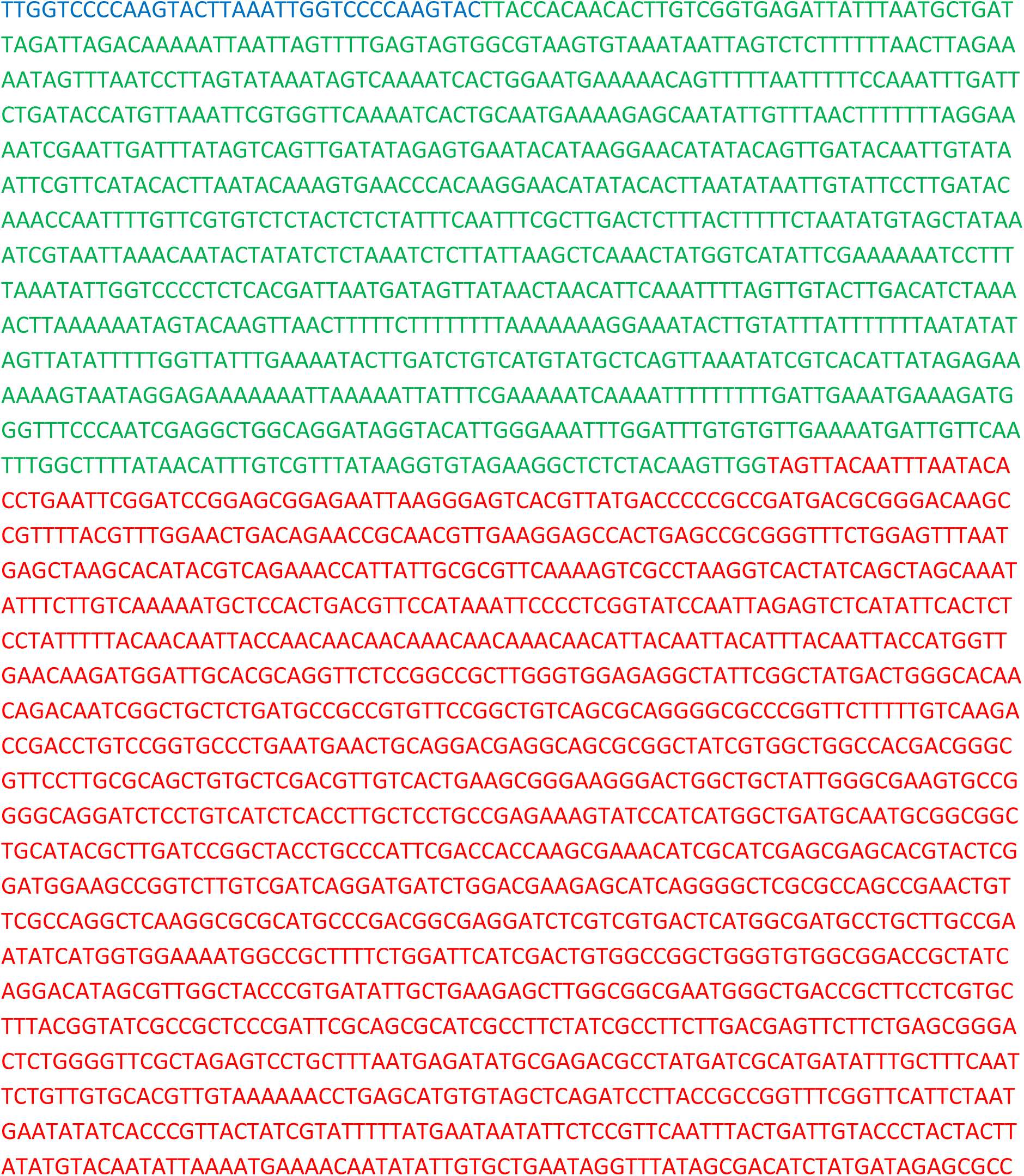
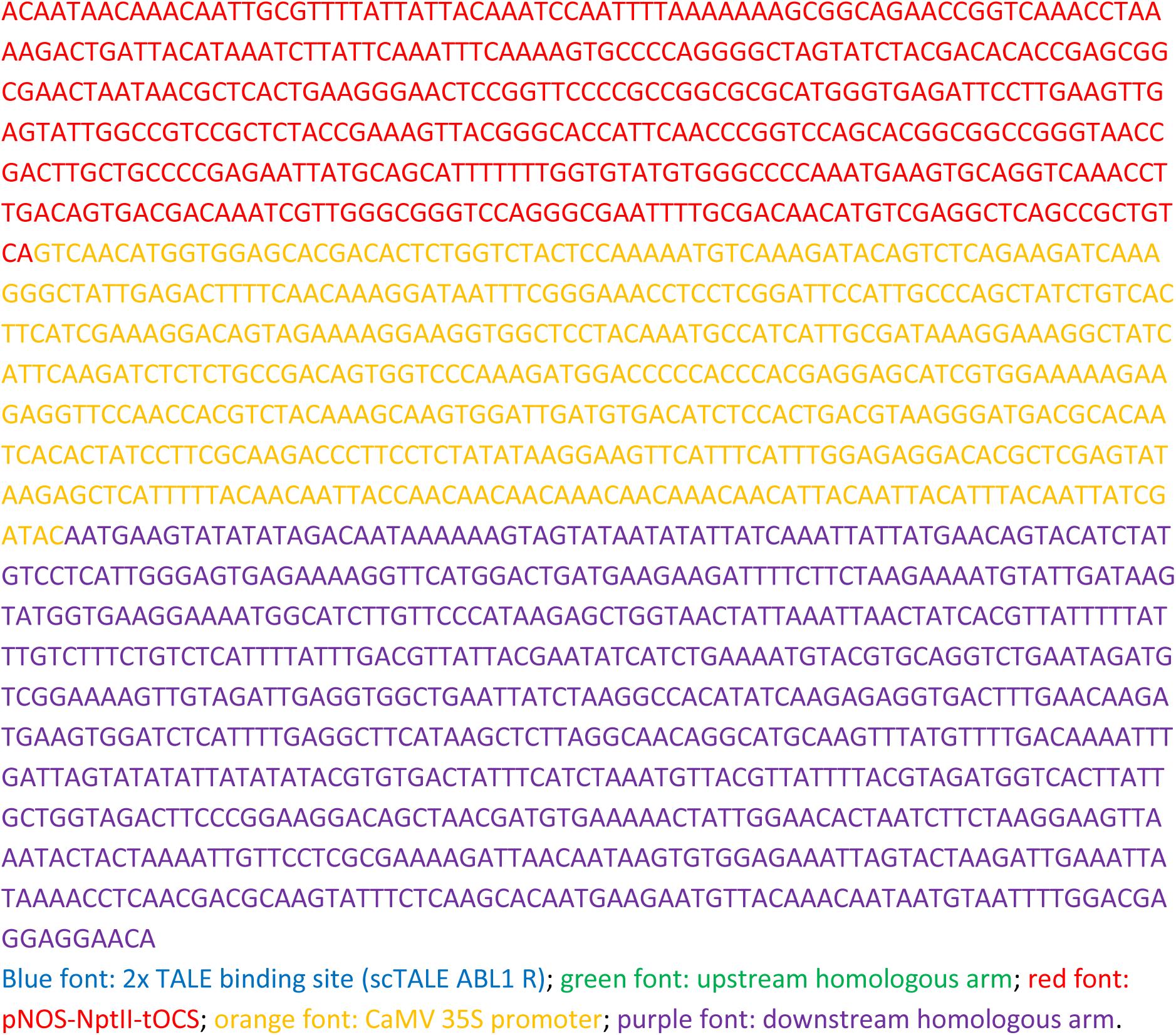

#### LbCpf1 expression cassette

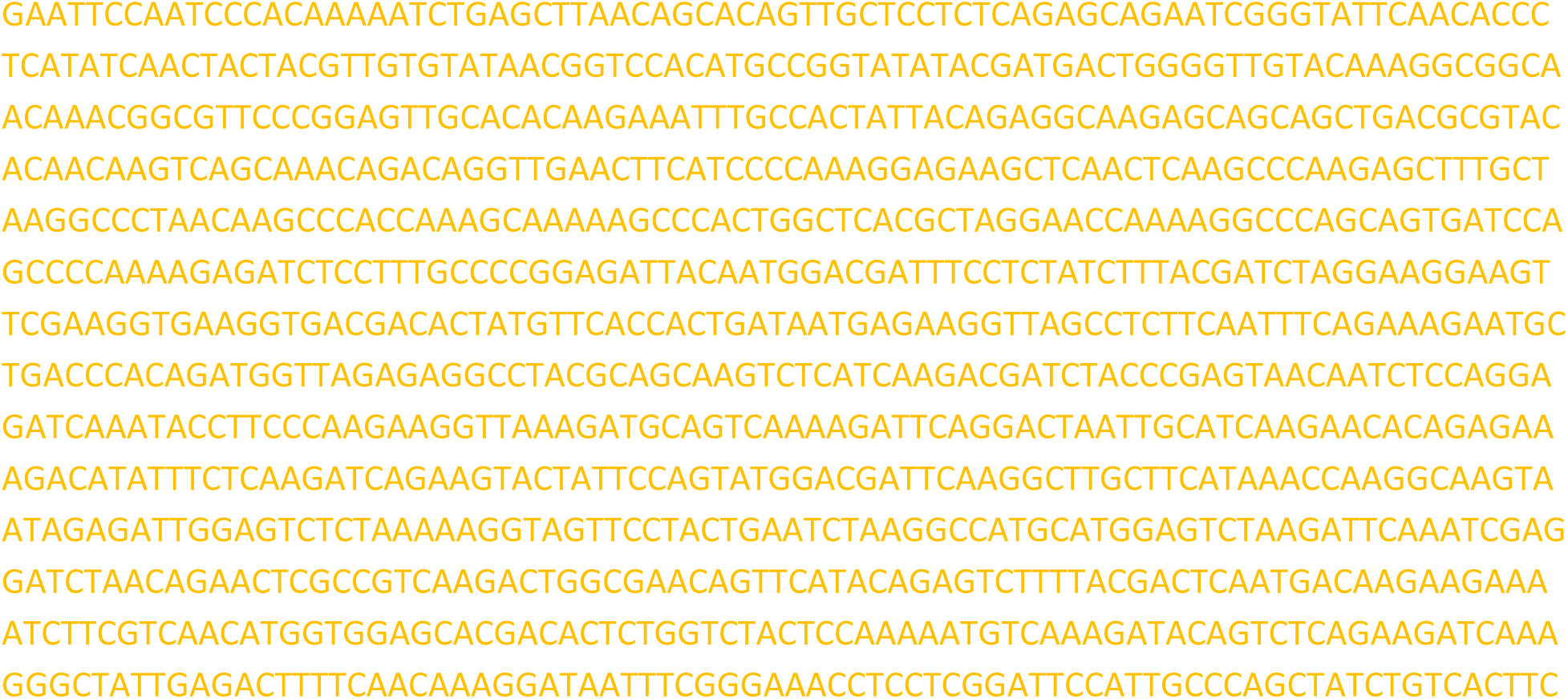
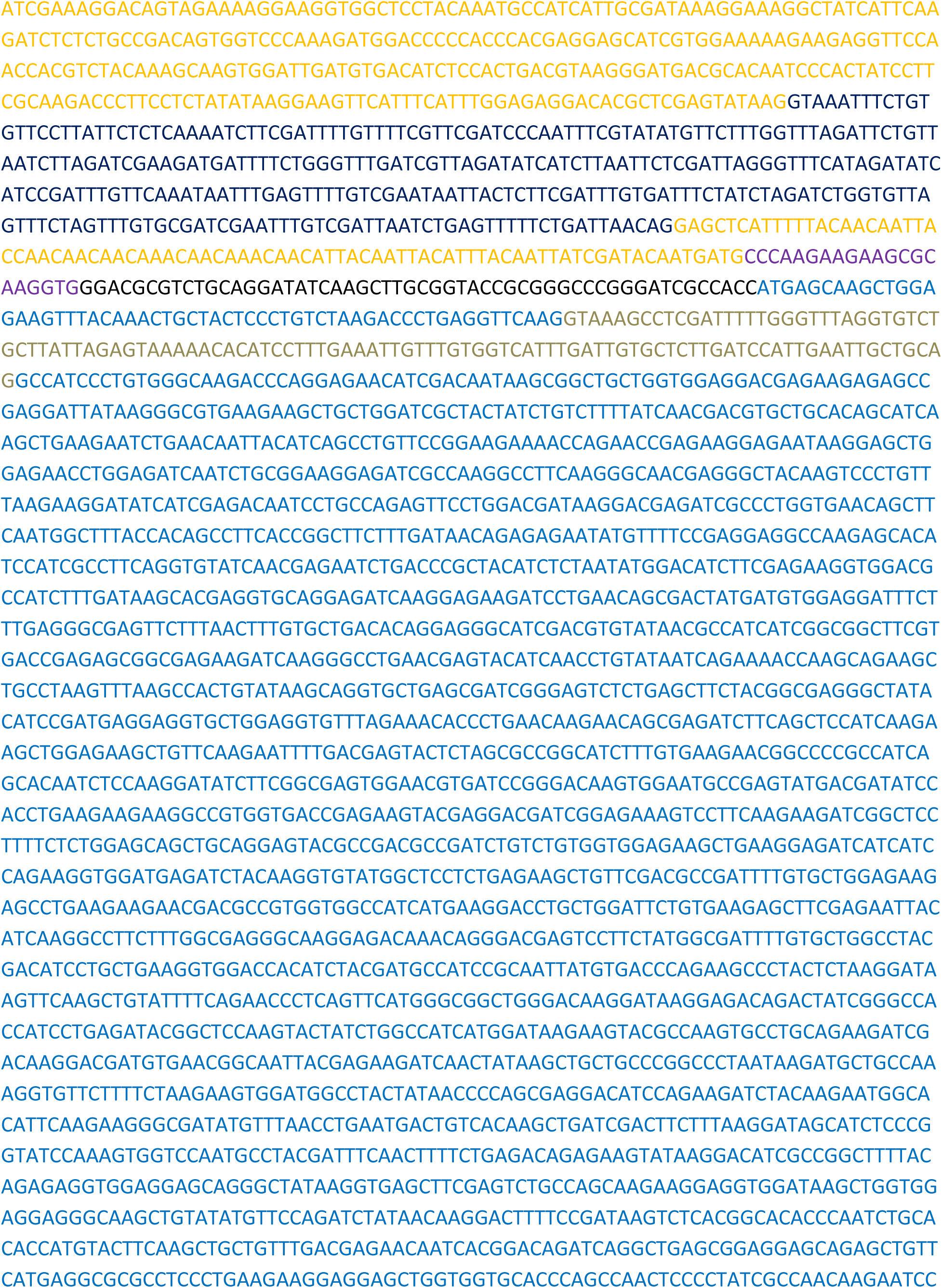
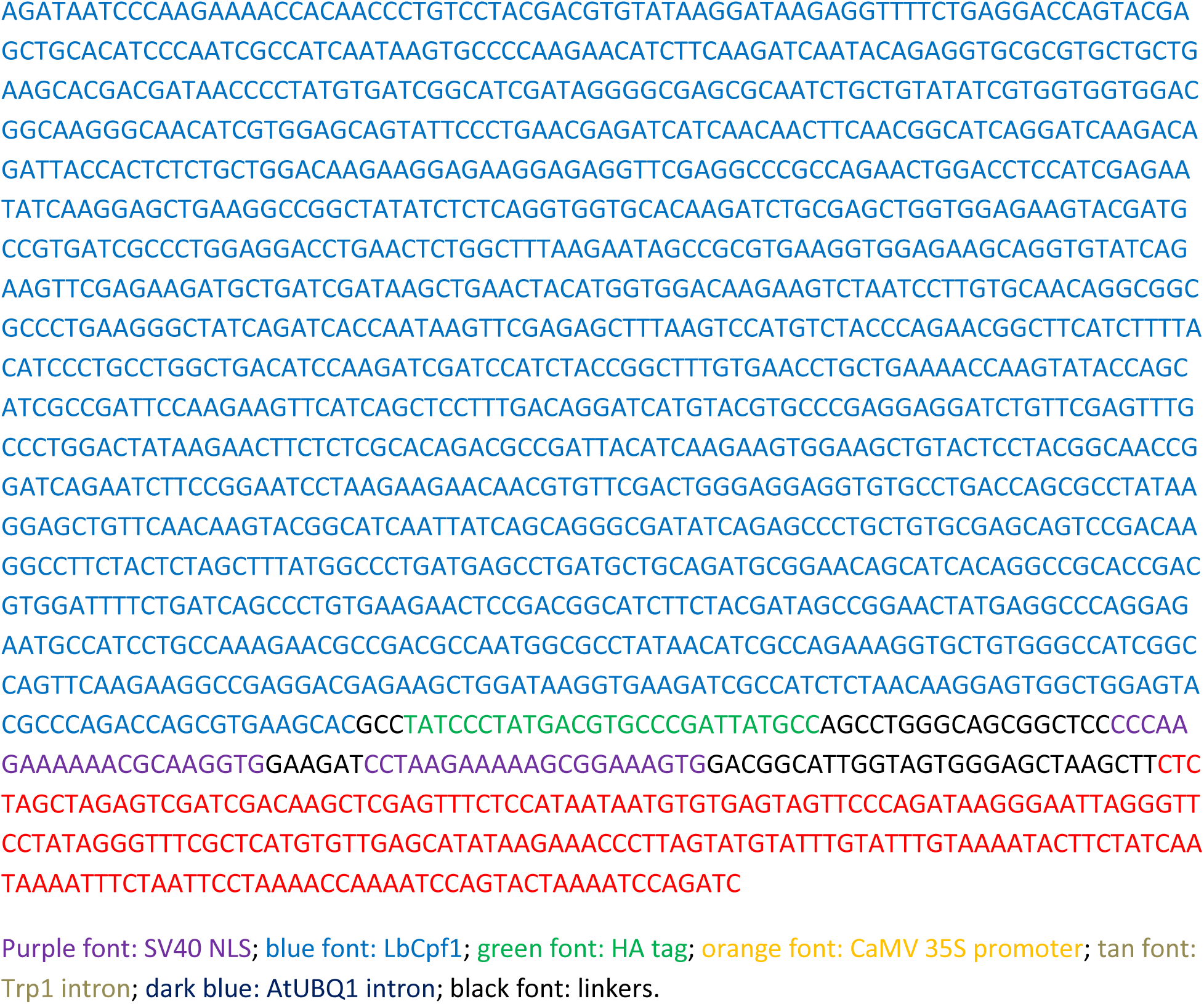

### Multiple replicon with Golden gate level 2 acceptor sites

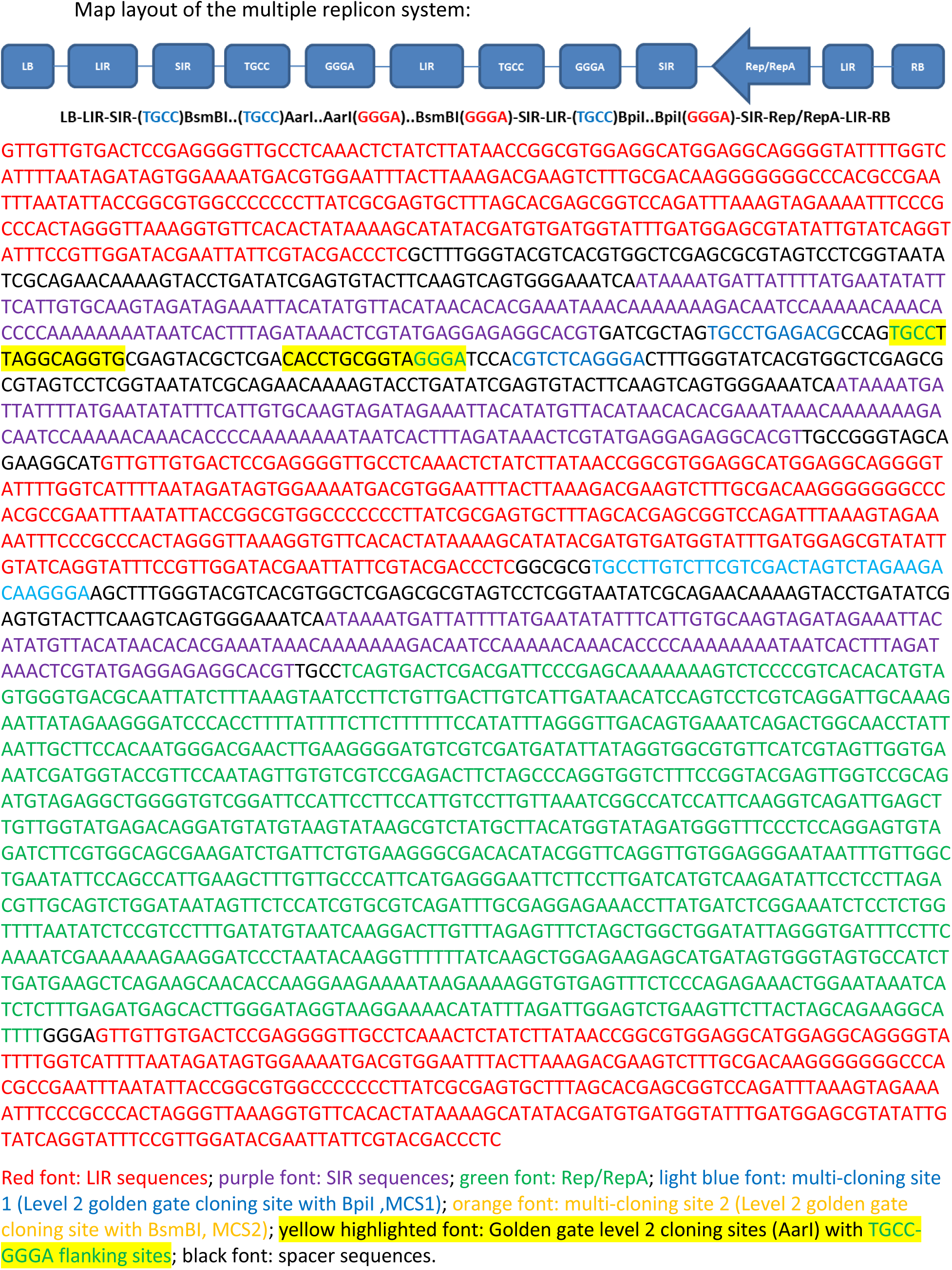

### In pHRHKT12.1 for editing SlHKT1;2 (Fig. 4B)

#### The dual KHT12 crRNA expression cassette

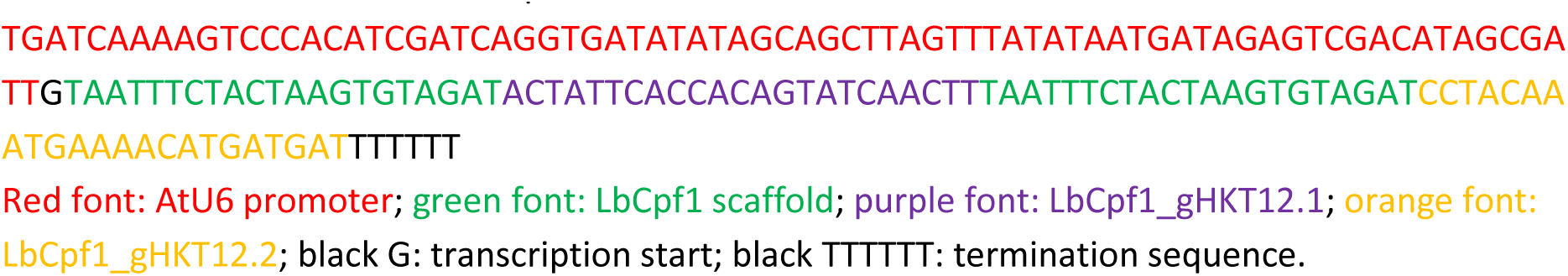

#### HKT1;2 donor sequence

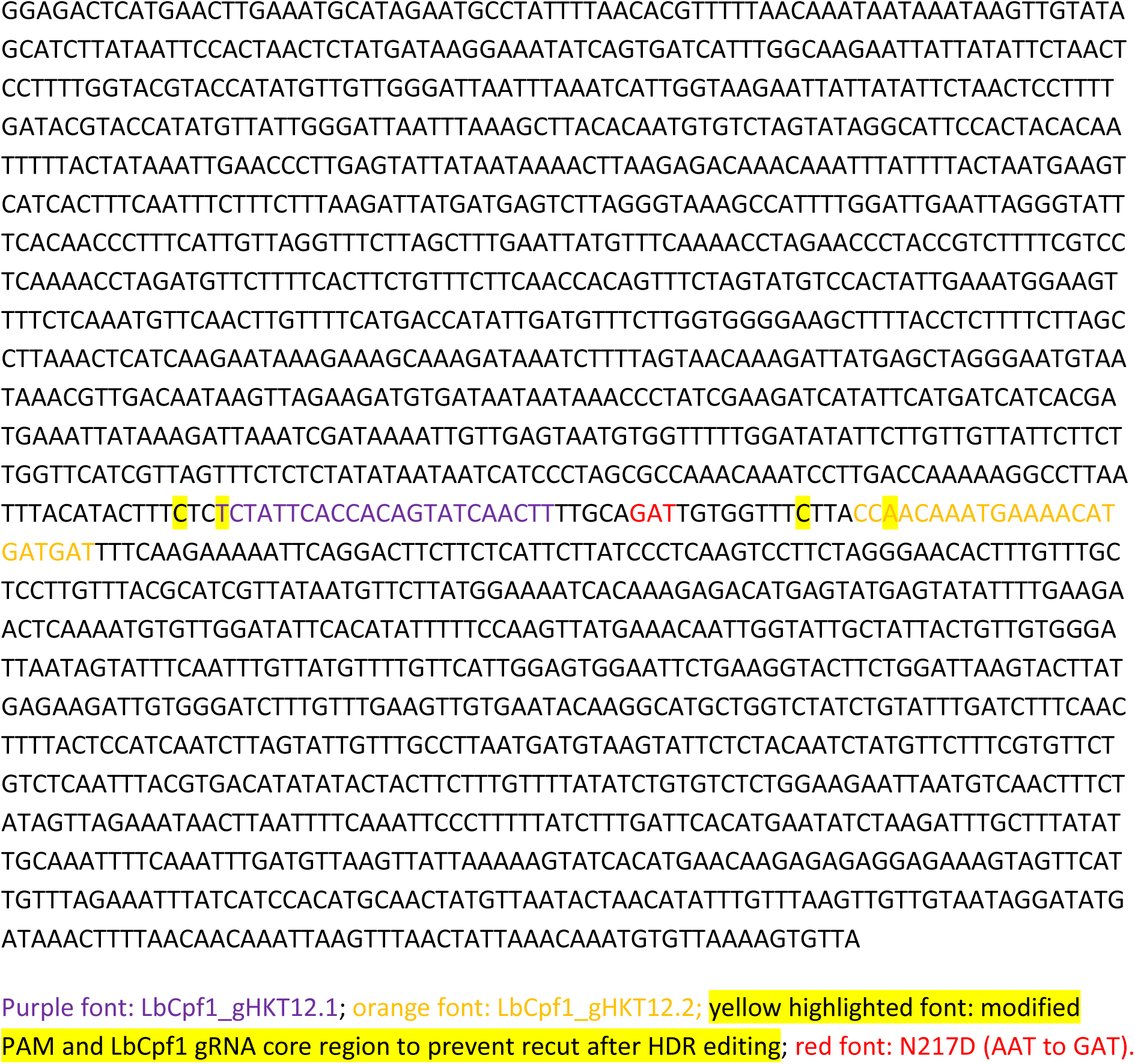

#### LbCpf1 expression cassette

Same as used in pHR01

#### Plant selection marker

pNOS-NptII-tOCS cloned from pICSL11024 (pICH47732::NOSp-NPTII-OCST) (Addgene Plasmid #51144).

Primers used for analyzing SlHKT1;2 events

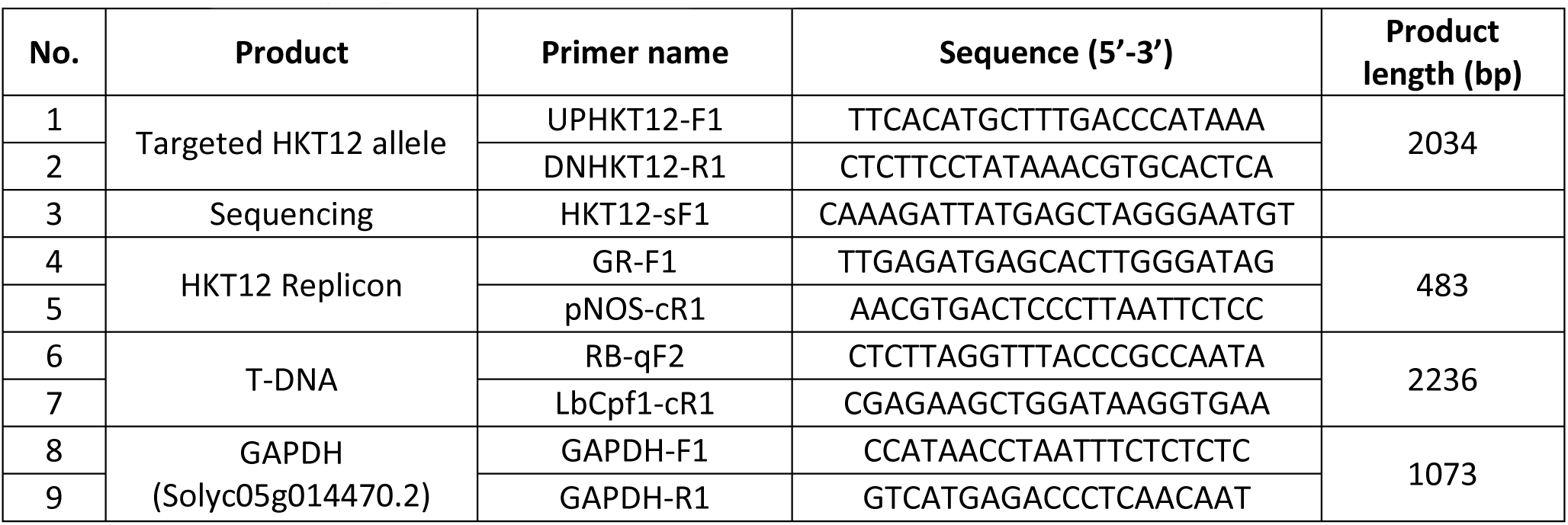

### pANT1^ox^

#### SlANT1 overexpression cassette

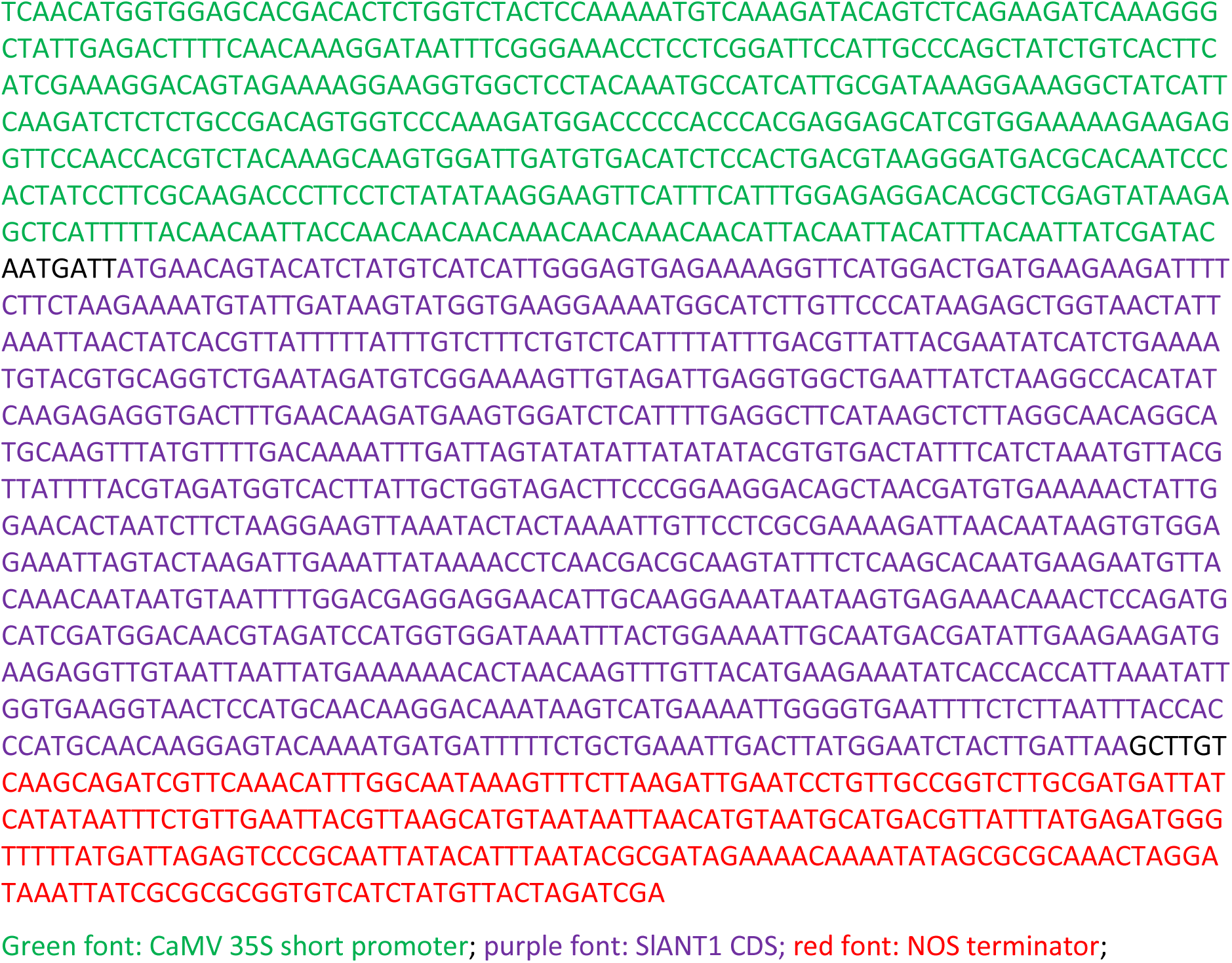

#### Plant selection marker

pNOS-NptII-tOCS cloned from pICSL11024 (pICH47732::NOSp-NPTII-OCST) (Addgene Plasmid #51144).

### SlRAD51 expression cassette of MR03

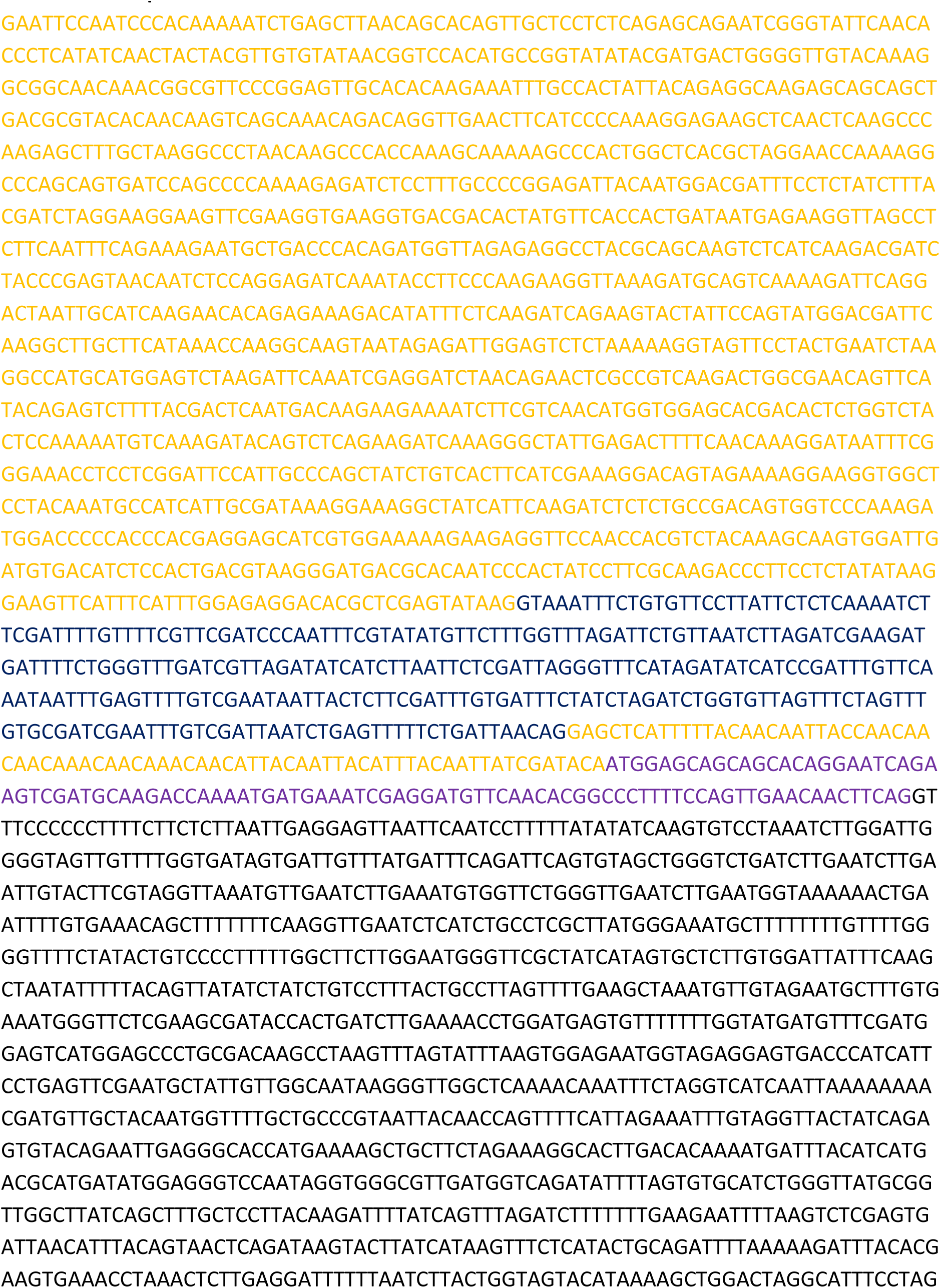
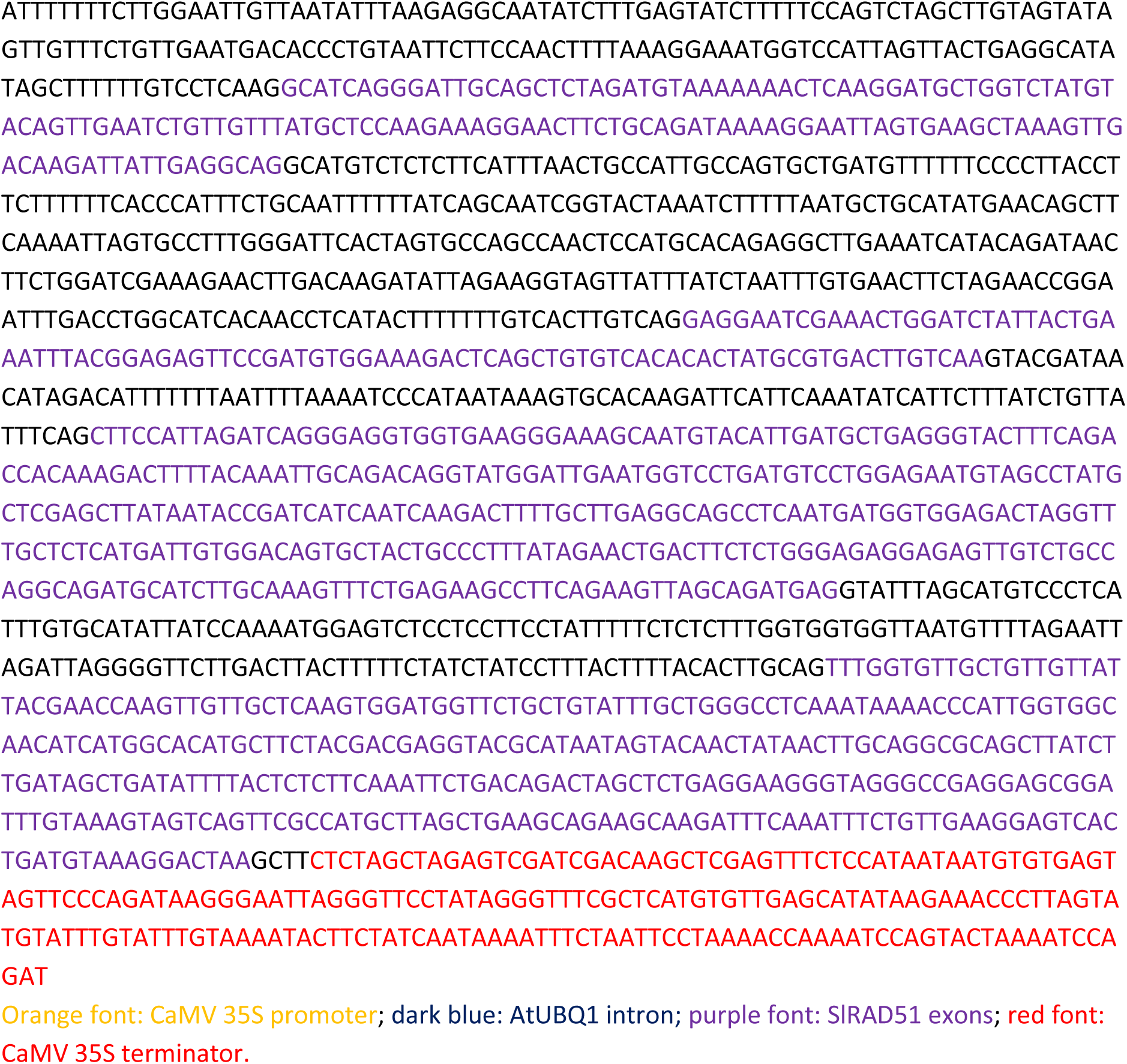

### SlRAD54 expression cassette of MR04

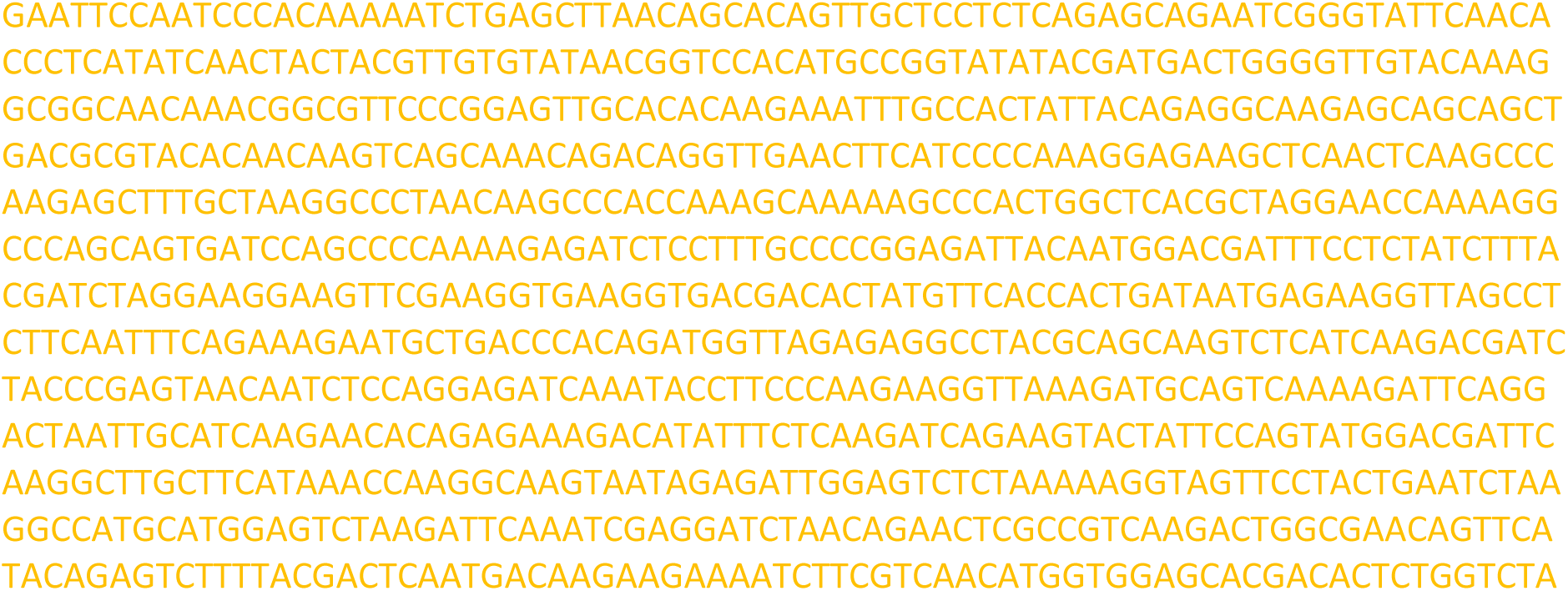
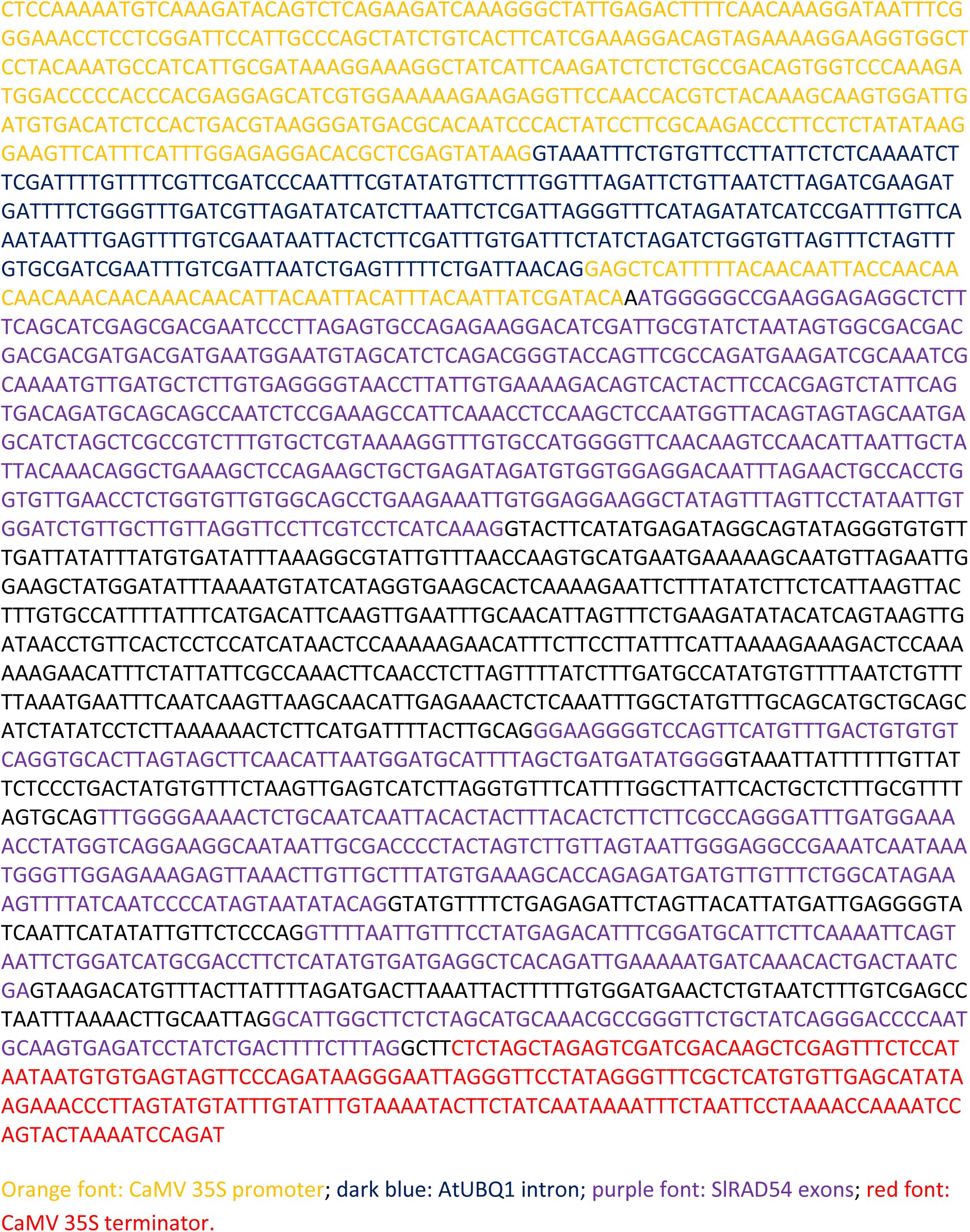

### Assessment of guide RNA activity via indel mutation traces of PCR products flanking targeted sites decomposed by ICE Synthego software

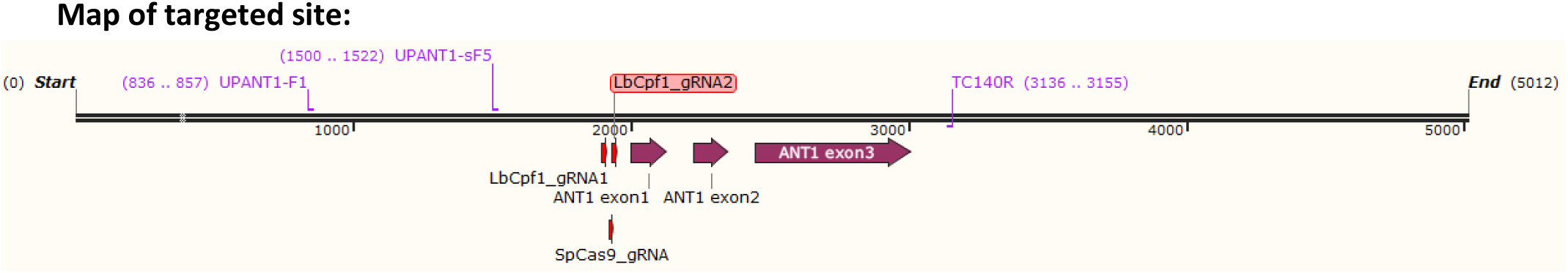

#### Sequences of the guide RNAs

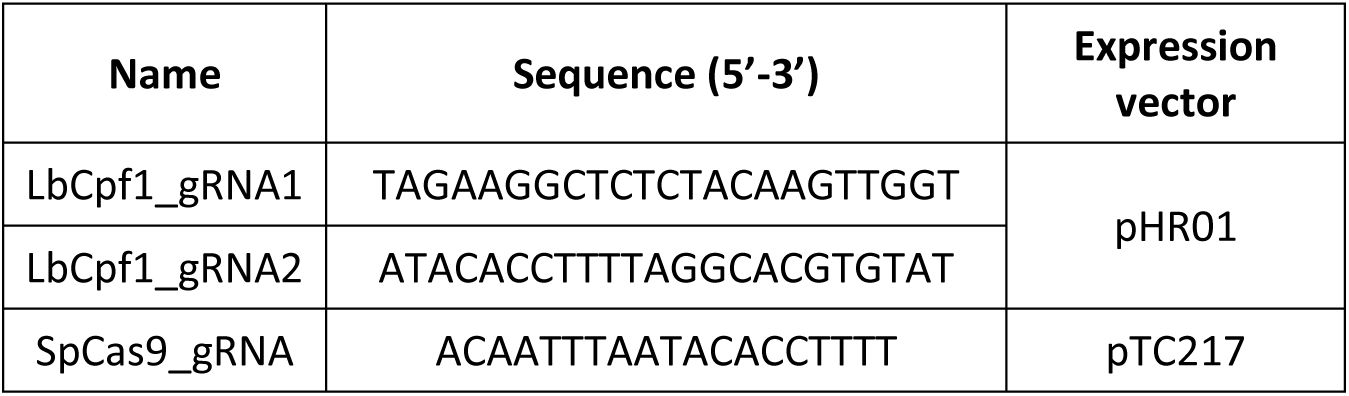

### Method used for assessment of activity of the guide RNAs

#### Decomposition of Sanger sequencing data using ICE Synthego software

Method: PCR products amplified using the primers (see table below) flanking the targeted sites of SlANT1 in pHR01 (ID11 to ID121) and pTC217 (ID91 to ID920)-transformed events and WT were purified on 0.8% agarose gel and subjected to Sanger sequencing and the sequencing data files (.ab1 extension) were decomposed using ICE Synthego (Hsiau et al., 2019). The WT sequencing ab1 file was used as the reference sequence for assessment of any DNA modification at the flanking site of that of the ANT1 events (Figure A and B). The results are summarized in Table A and B.

#### Primers used for amplifying flanking region of targeted sites

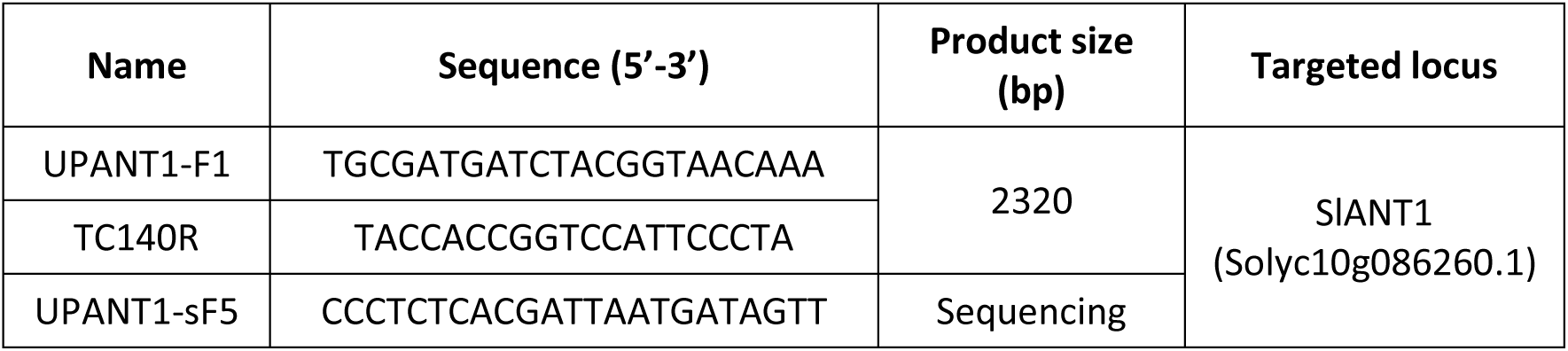

**Fig. A.**
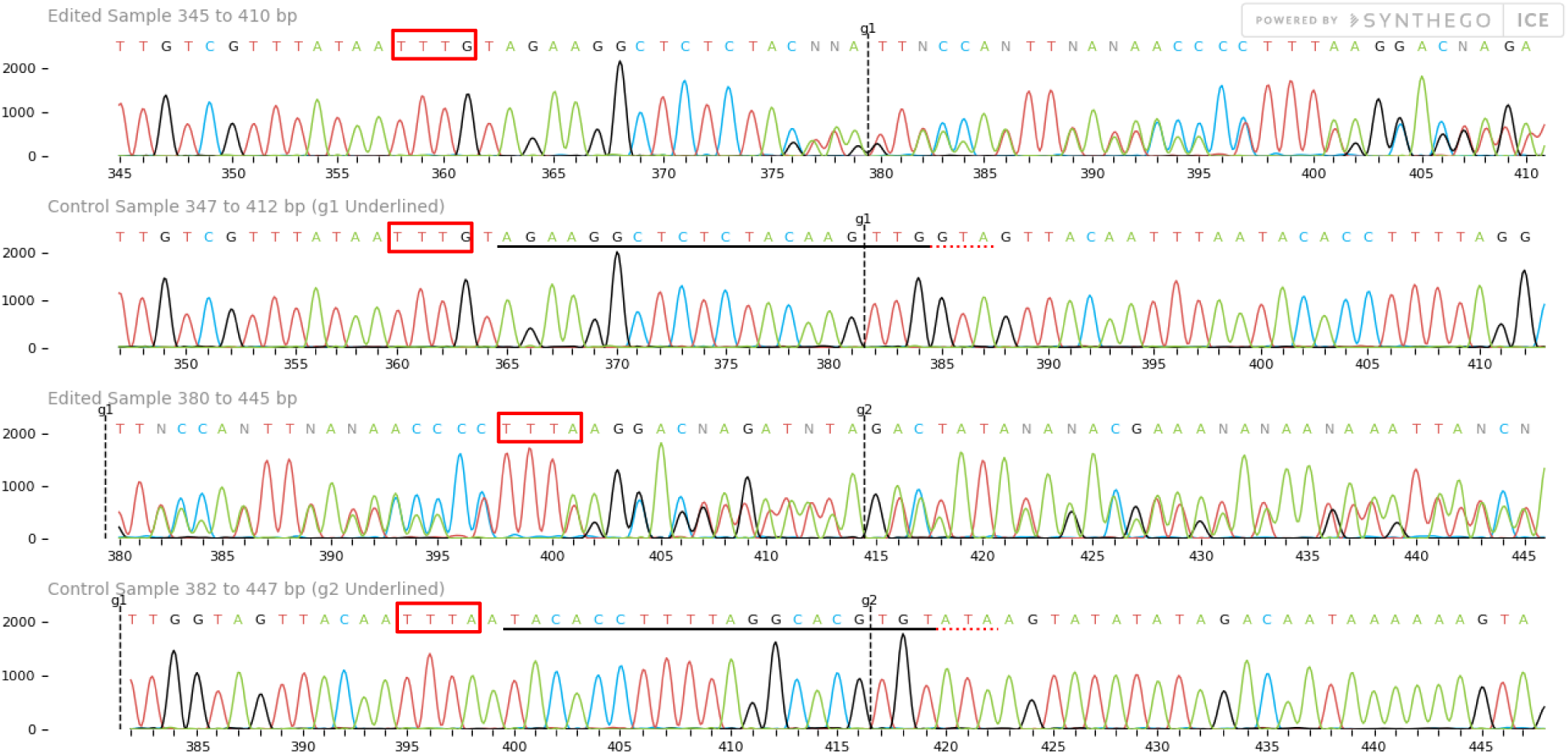
A representative diagram showing ICE Synthego sequence comparison of ID11 event (the first row for LbCpf1_gRNA1 and the third row for LbCpf1_gRNA2) and control sequence (the second and fourth rows). The vertical discontinuous lines denote hypothetical cutting site at 18bp downstream of the PAM sites (red boxes)

**Fig. B.**
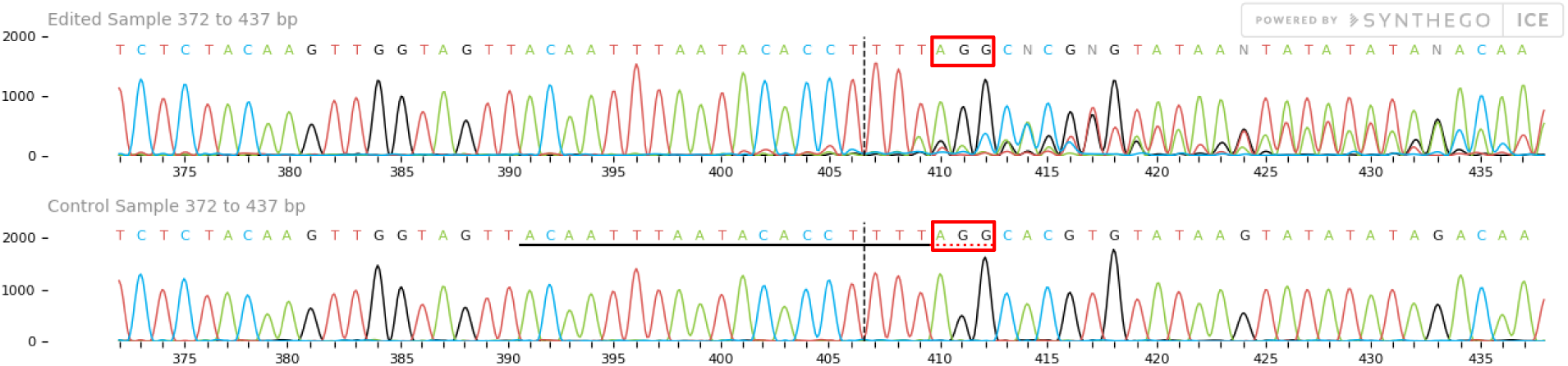
A representative diagram showing ICE Synthego sequence comparison of ID91 event (upper row) and control sequence. The vertical discontinuous lines denote hypothetical cutting site at 3 bp upstream of the PAM sites (red boxes).

**Table A:**
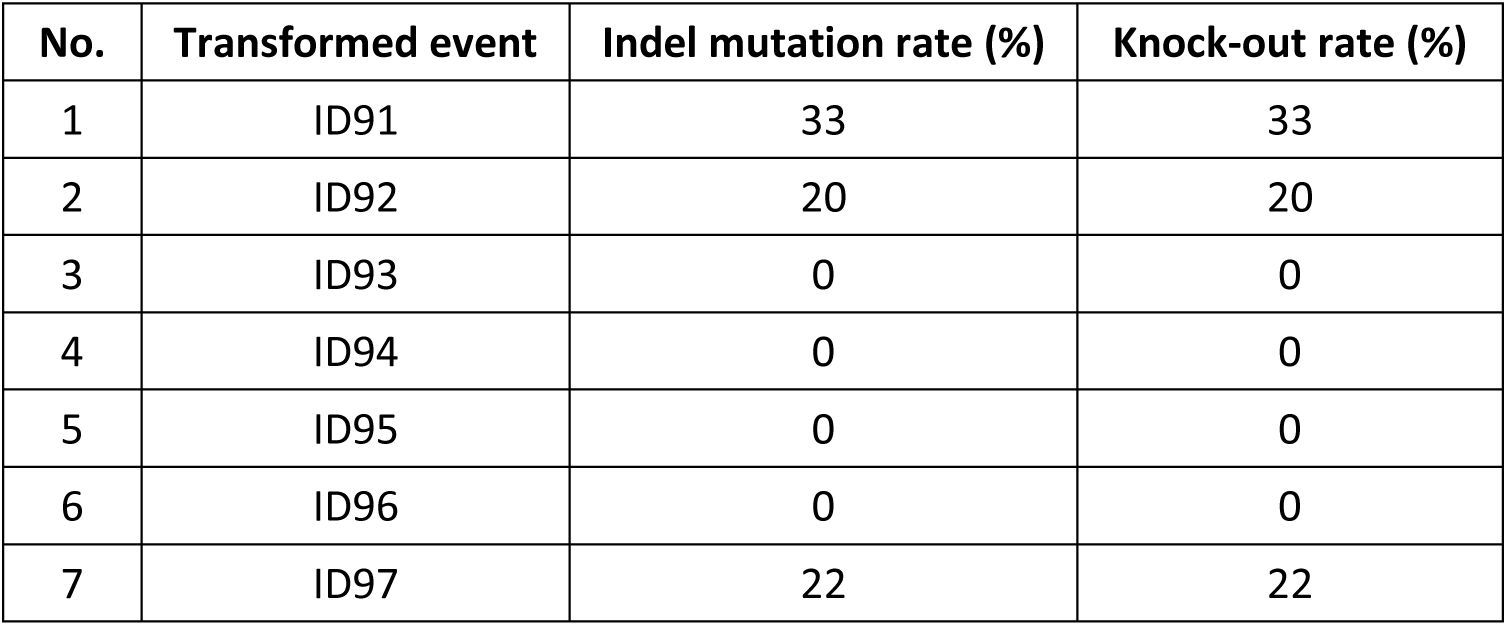
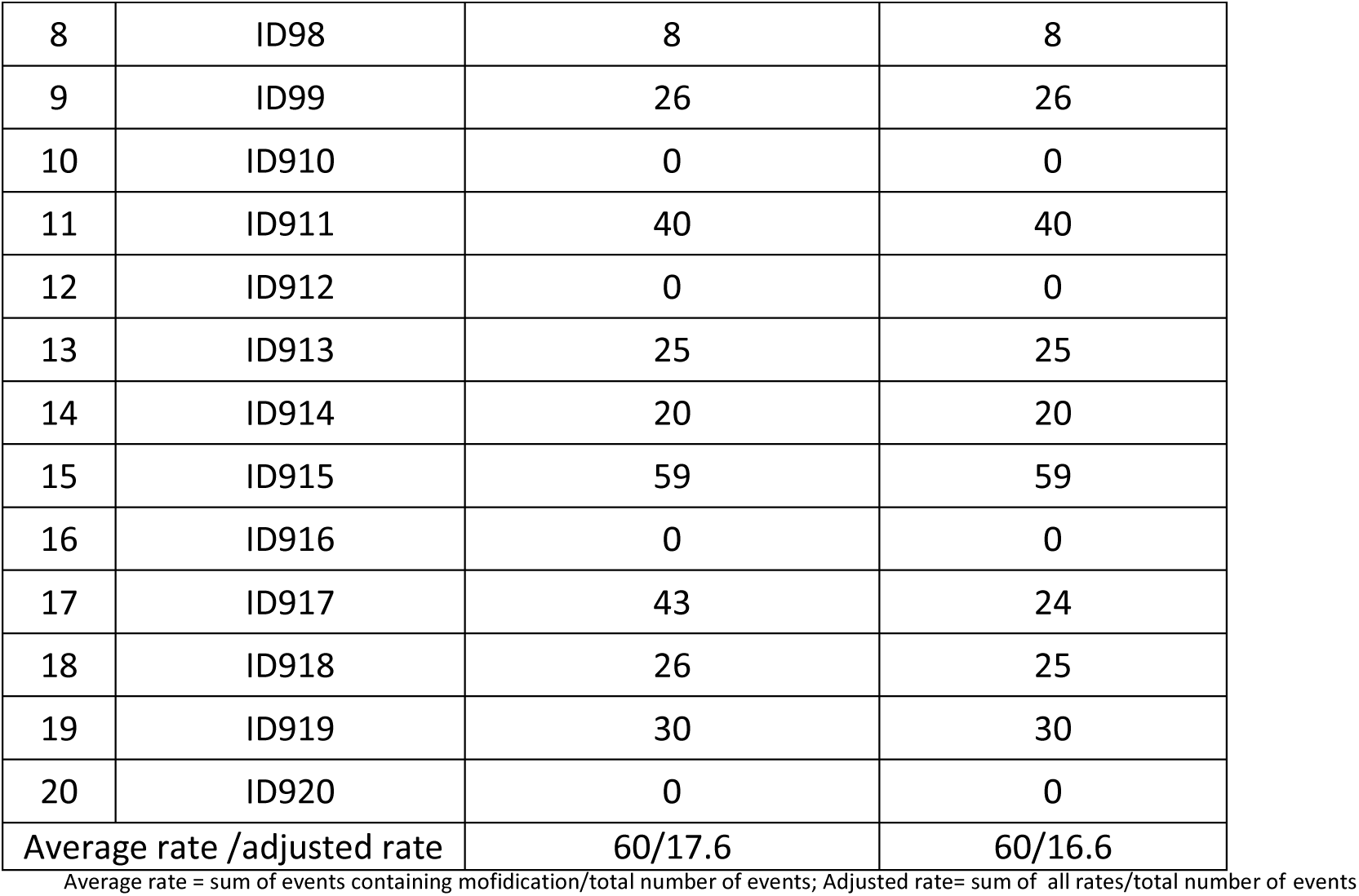
Indel mutation rates among SpCas9-based samples decomposed by ICE Synthego software

**Table B:**
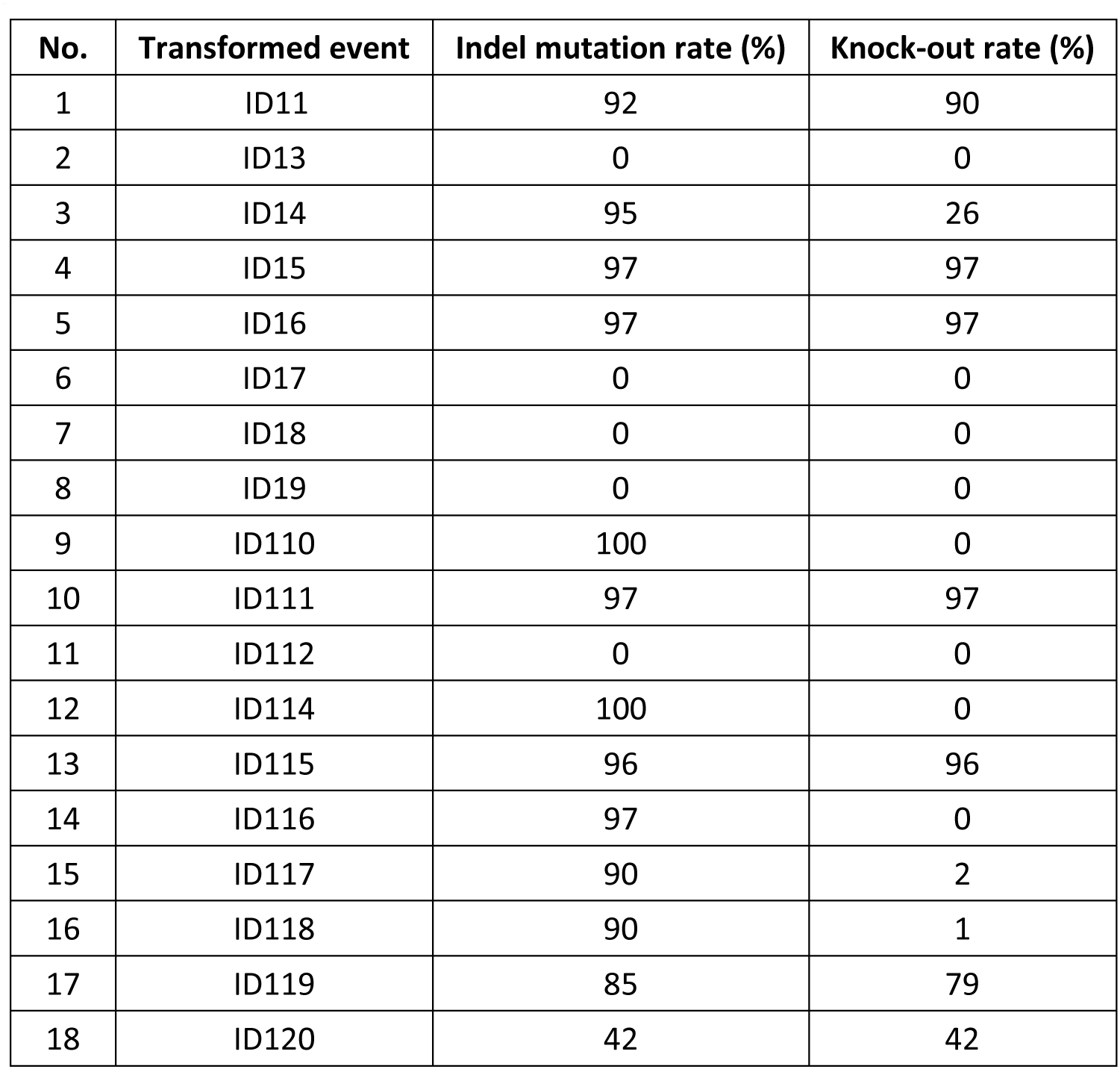
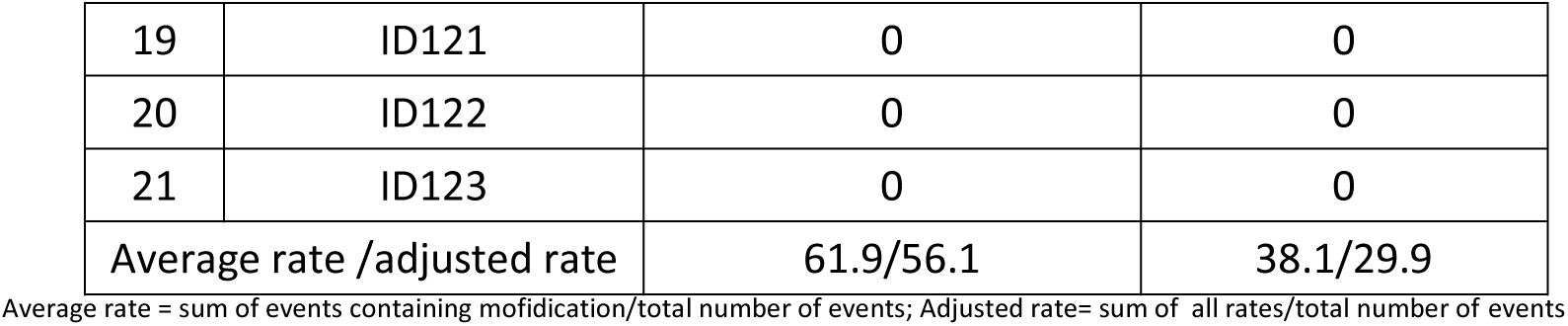
Indel mutation rates at SlANT1 locus recorded from pHR01 transformed events reported by ICE Synthego

## Off-target analysis for LbCpf1_gRNA1 and LbCpf1_gRNA2 using Cas-OFFinder (Bae et al., 2014)

Critetia for searching for off-target: 5’-TTTV-3’ PAM; maximum 4 mismatches; Tomato genome database: *Solanum lycopersicum* (SL2.4)

For LbCpf1_gRNA1:

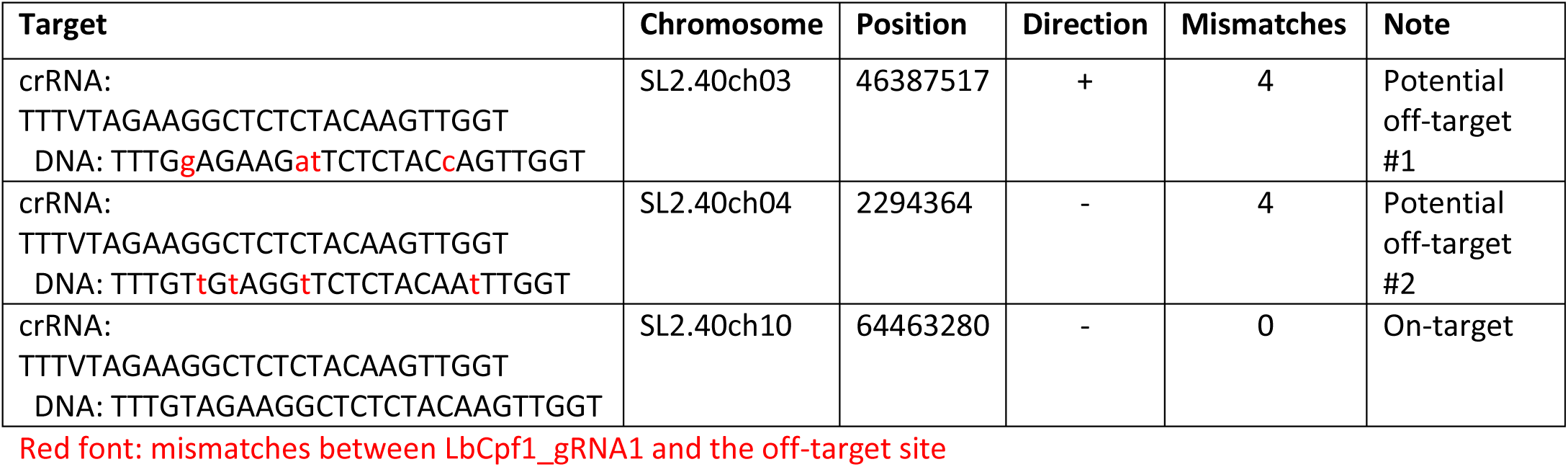

For LbCpf1_gRNA2:

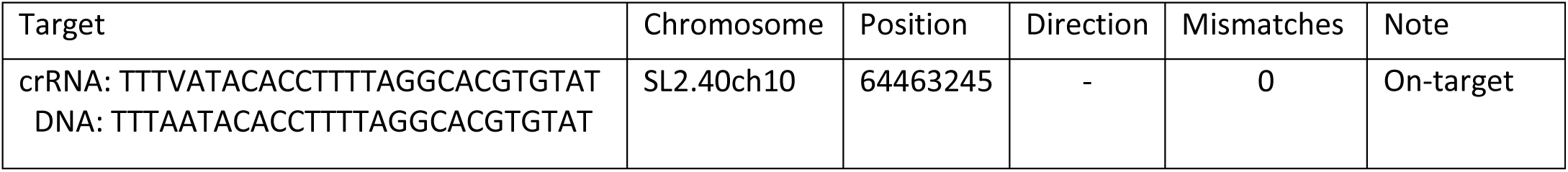

## Potential off-target #1 of LbCpf1_gRNA1: Tomato build SL2.40-SL2.40ch03-46386519..46388519

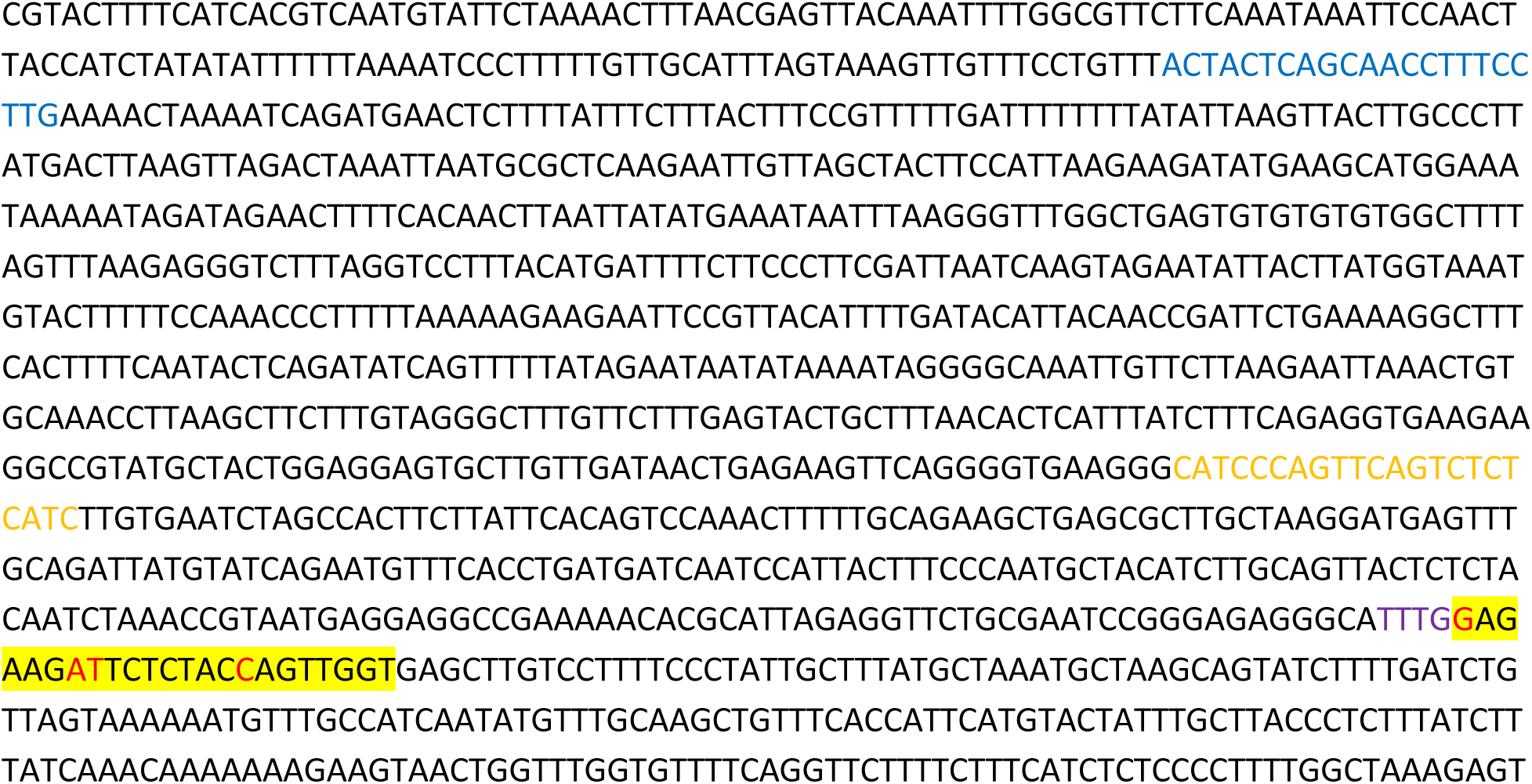
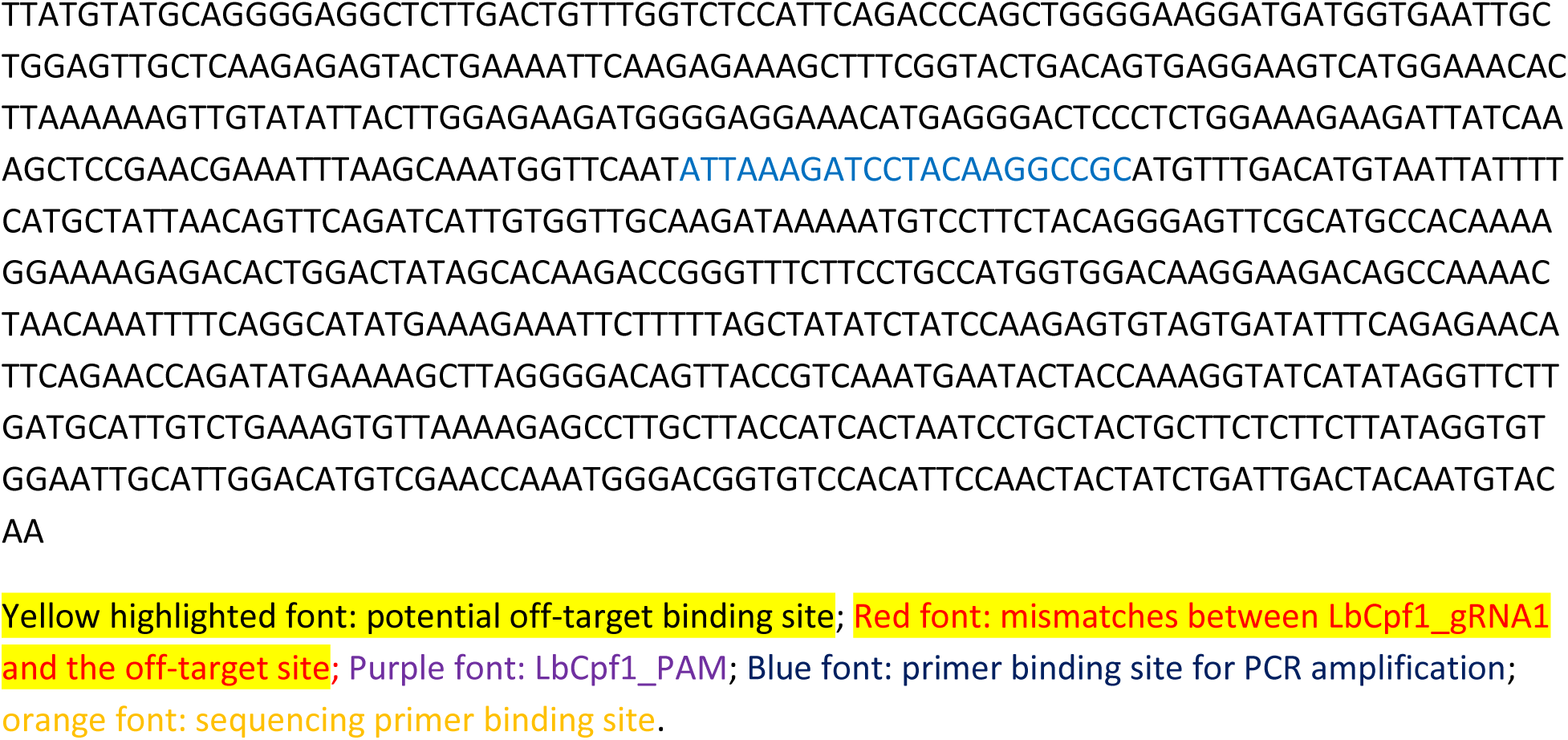

## Potential off-target #2 of LbCpf1_gRNA1: Tomato build SL2.40-SL2.40ch03-46386519..46388519

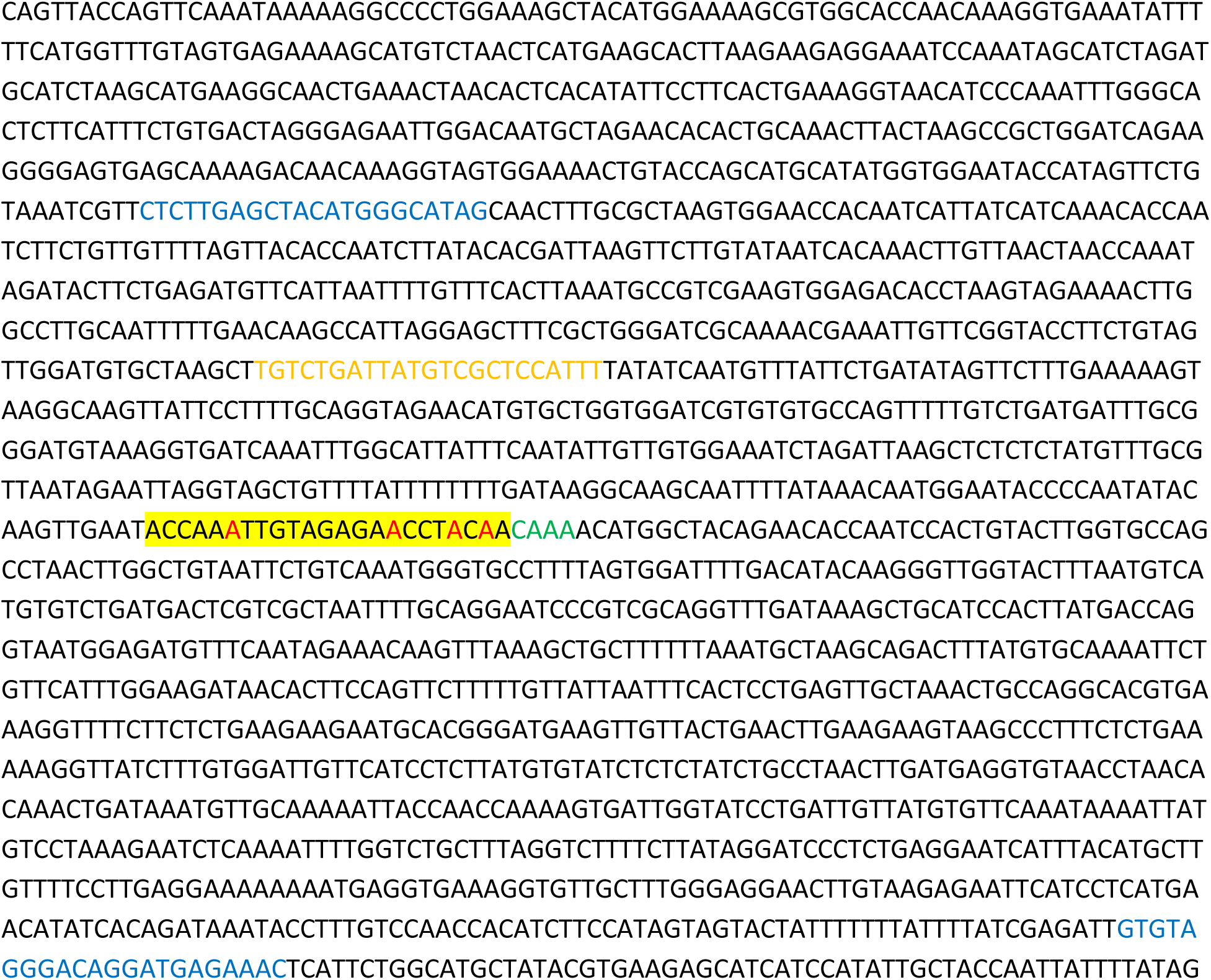
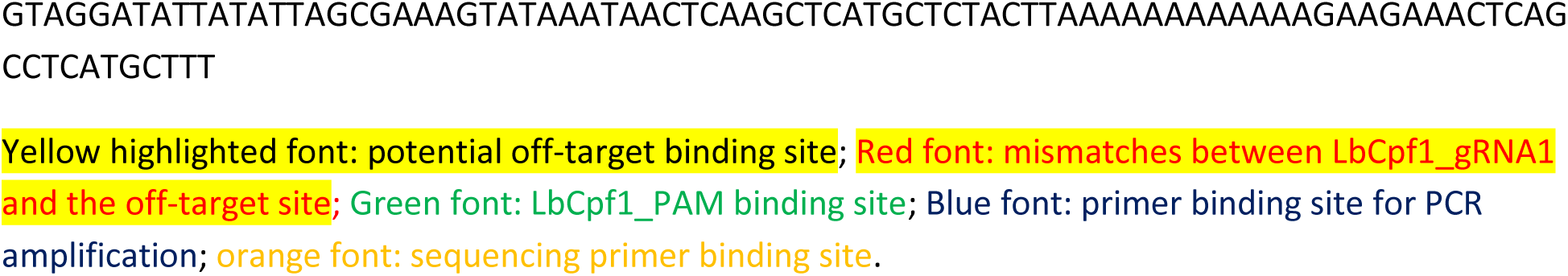

## Primers used for potential off-target analysis in this study

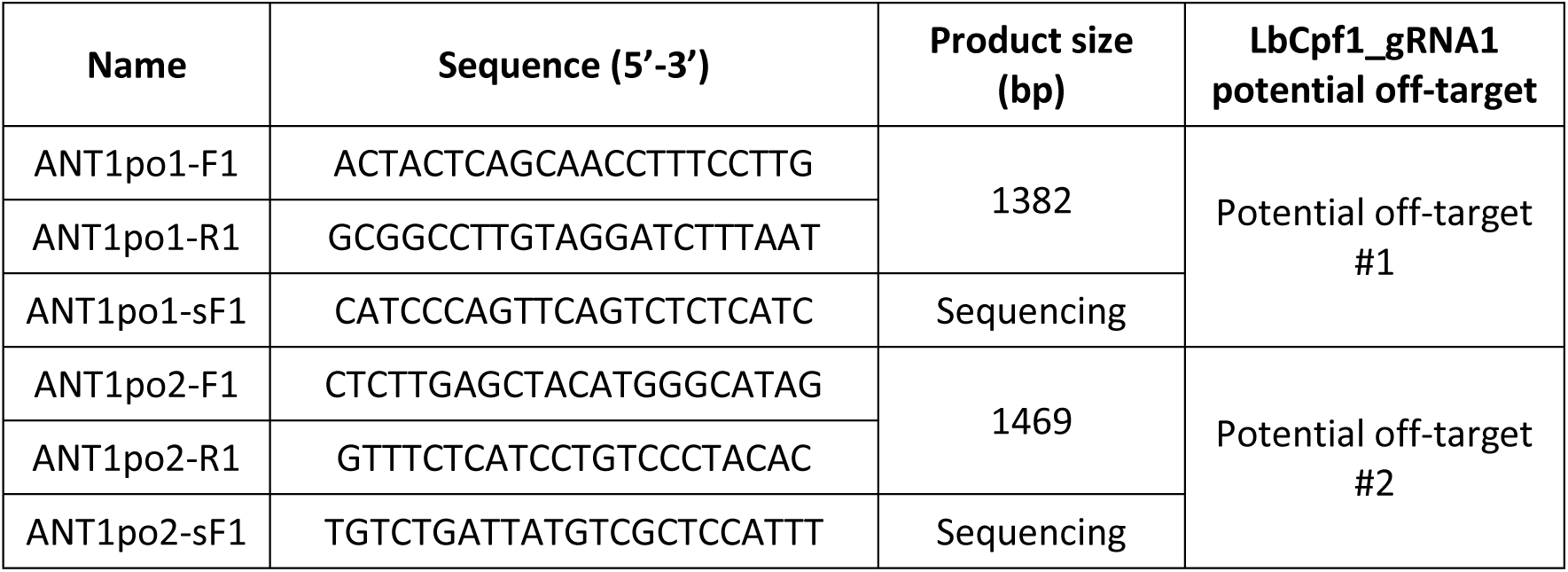

## Decomposition of Sanger sequencing data using ICE Synthego software

Method: PCR products (only single band at the expected size was observed) amplified using the primers with the ANT1 HDR events #C11; C12; C13; C141; C142; C18; C19; C110; C111; C112; C113; C114; C115; C116; C117 and WT were purified on 0.8% agarose gel and subjected to Sanger sequencing and the sequencing data files (.ab1 extension) were decomposed using ICE Synthego (Hsiau et al., 2019). The WT sequencing ab1 file was used as the reference sequence for assessment of any DNA modification at the flanking site of that of the ANT1 HDR events.

No off-targeted modification was found at both the potential off-target sites in all the tested samples:

**Fig. A.**
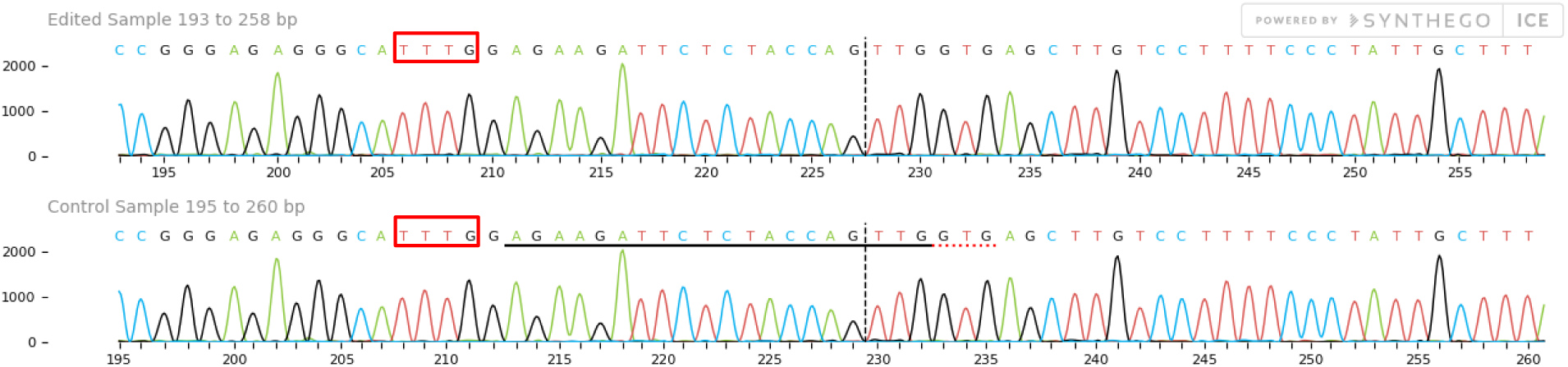
A representative diagram showing ICE Synthego sequence comparison of C11 event (upper row) and control sequence at the potential off-target site #1 of LbCpf1_gRNA1. The vertical discontinuous lines denote hypothetical cutting site at 18bp downstream of the sequence containing PAM sites (red boxes).

**Fig. B.**
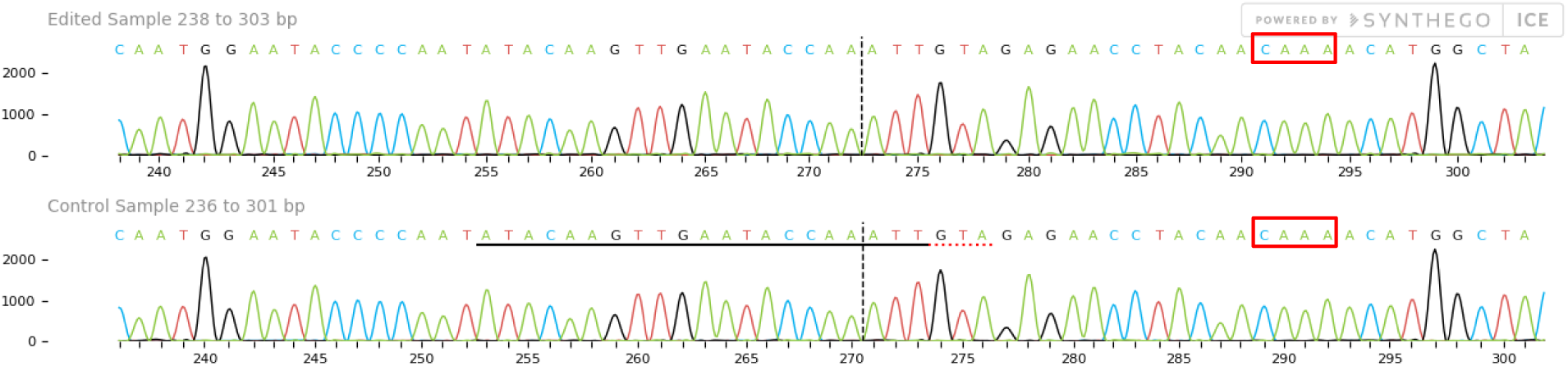
A representative diagram showing ICE Synthego sequence comparison of C11 event (upper row) and control sequence at the potential off-target site #1 of LbCpf1_gRNA1. The vertical discontinuous lines denote hypothetical cutting site at 18bp downstream of the sequence containing PAM sites (red boxes)

**Table A:**
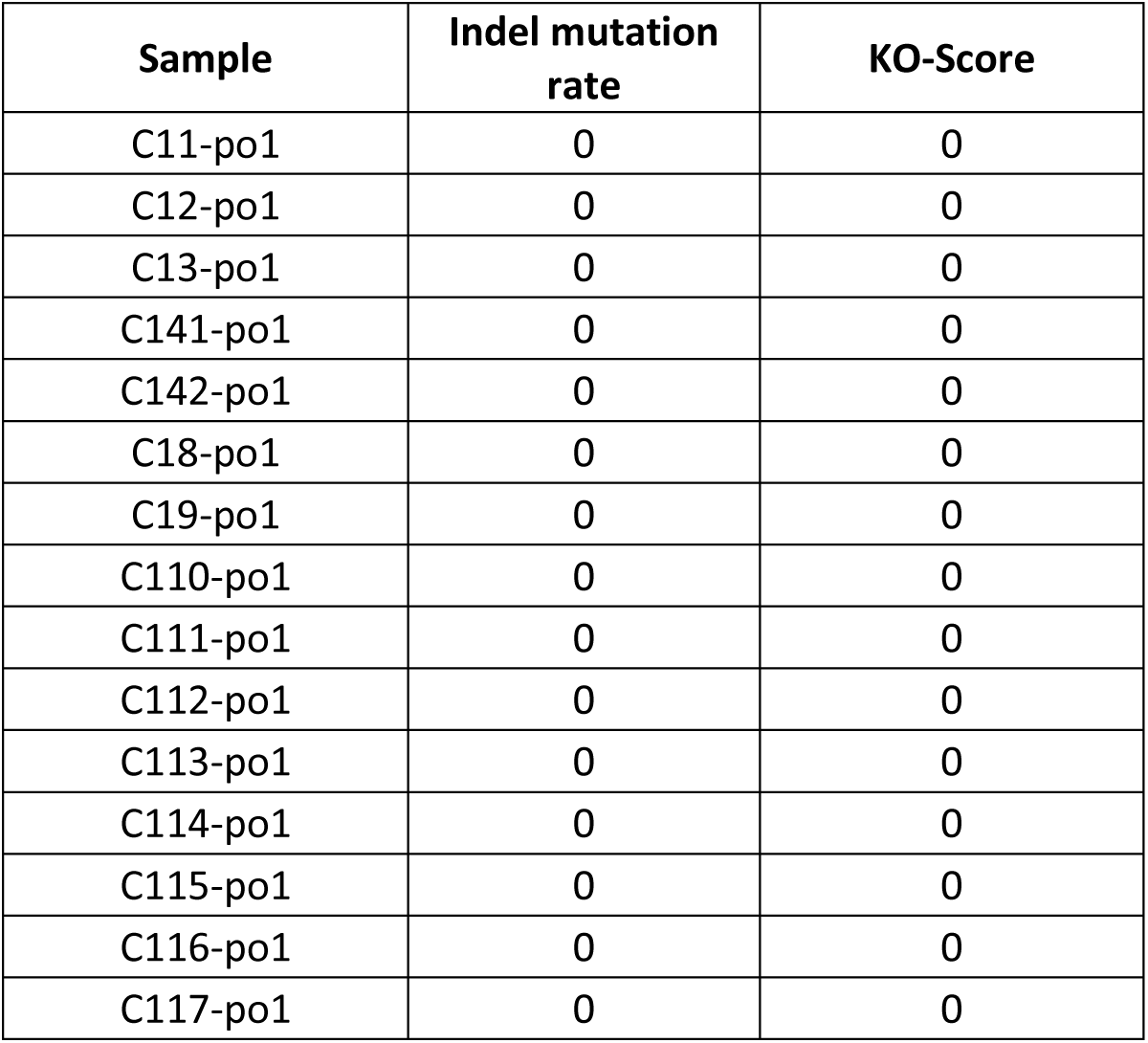
Knock-out scores of the potential off-target site #1 reported by ICE Synthego

**Table B:**
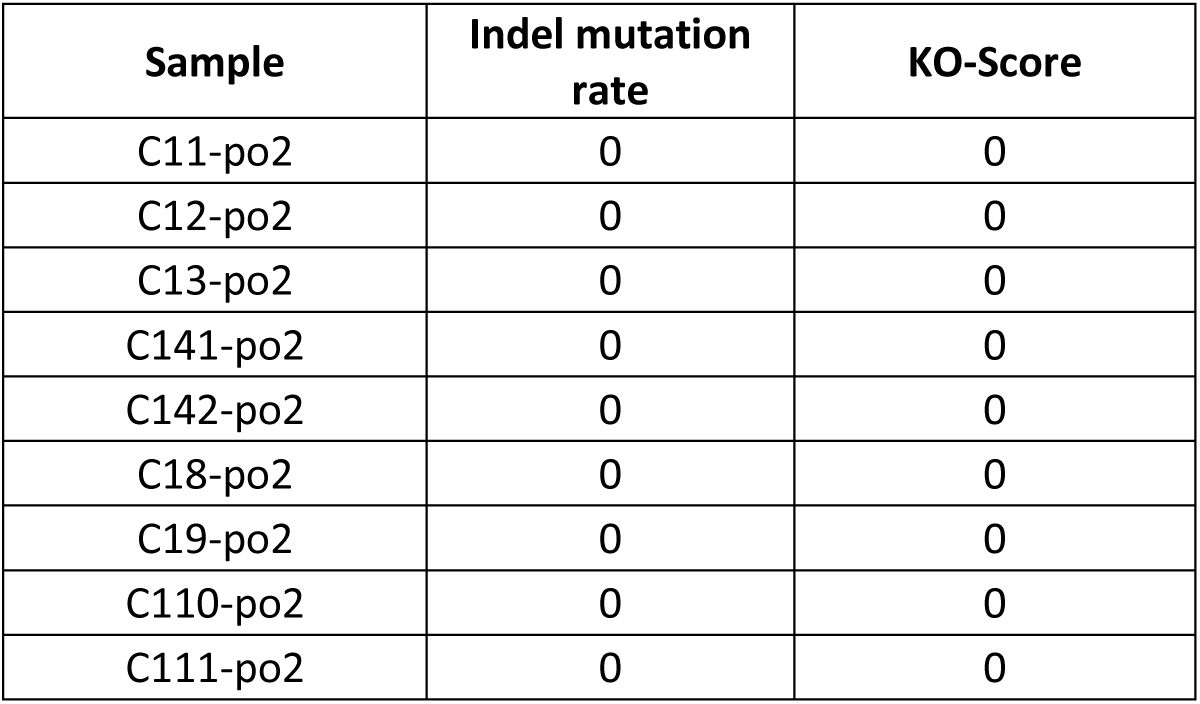
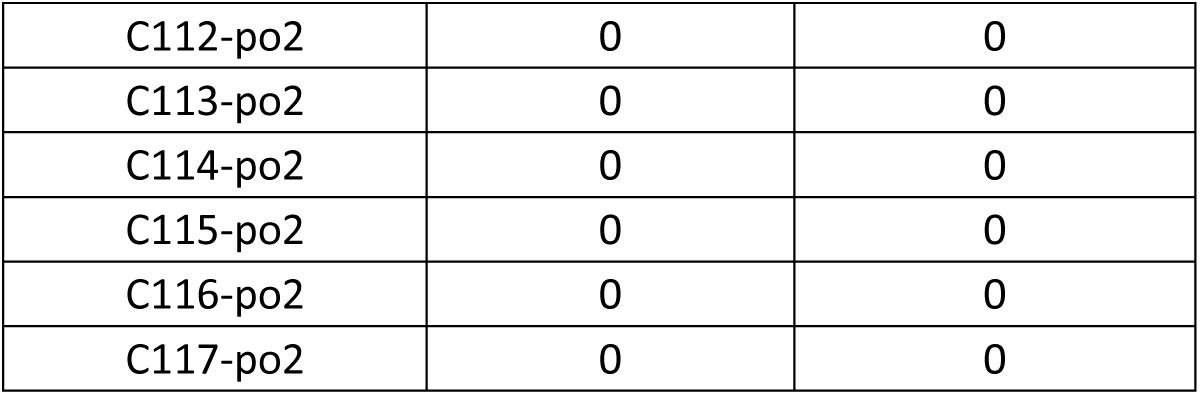
Knock-out scores of the potential off-target site #2 reported by ICE Synthego

## Analysis of RNA transcript levels of RAD51 and RAD54 expressed in pMR03 or pMR04 transformed events by qRT-PCRs

### Linearized maps of SlRAD51 and SlRAD54 loci and oligos used for qRT-PCRs

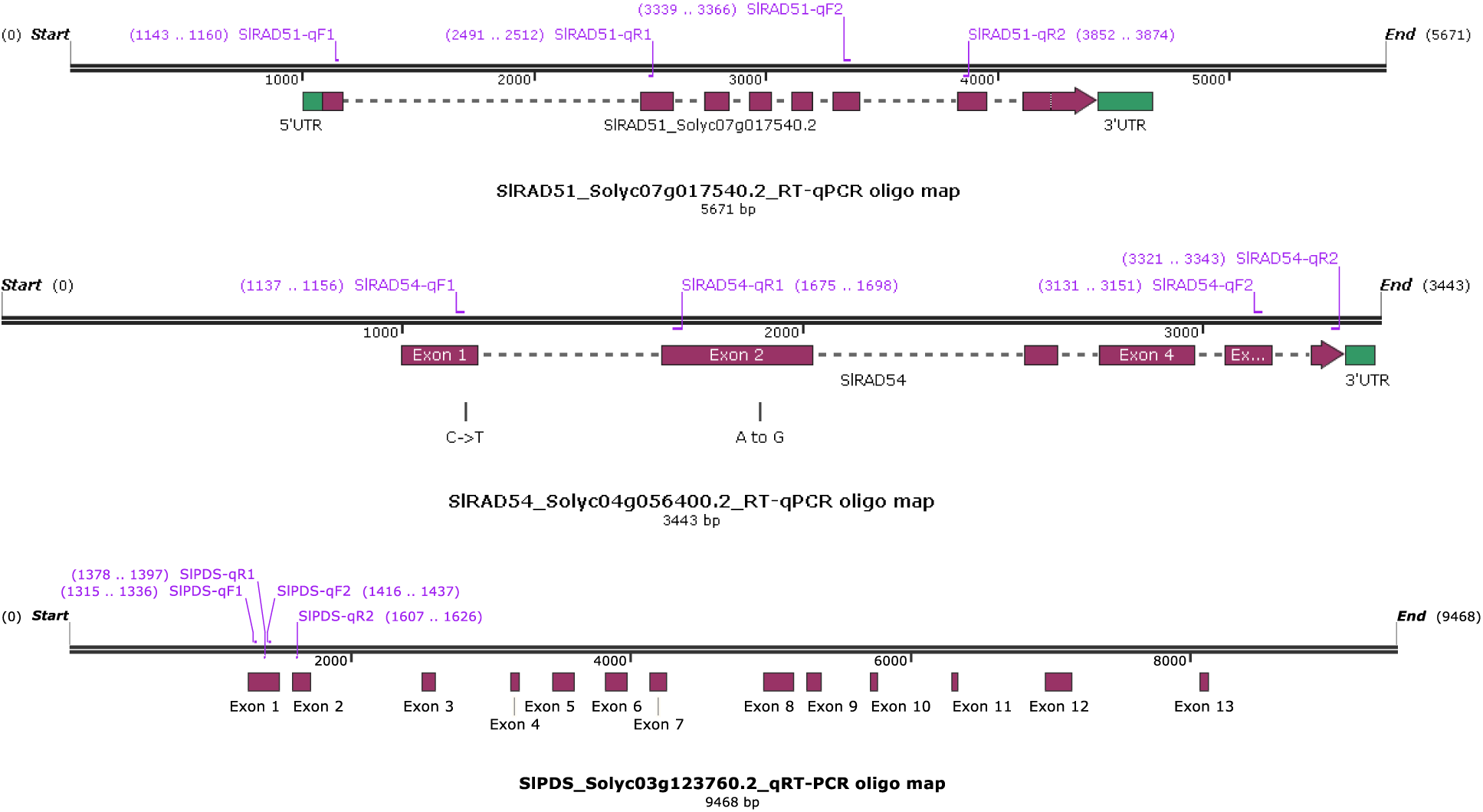

- We designed and evaluated two pairs of primers for each of the locus that helped amplifying exon region.

### Primers used in this analysis

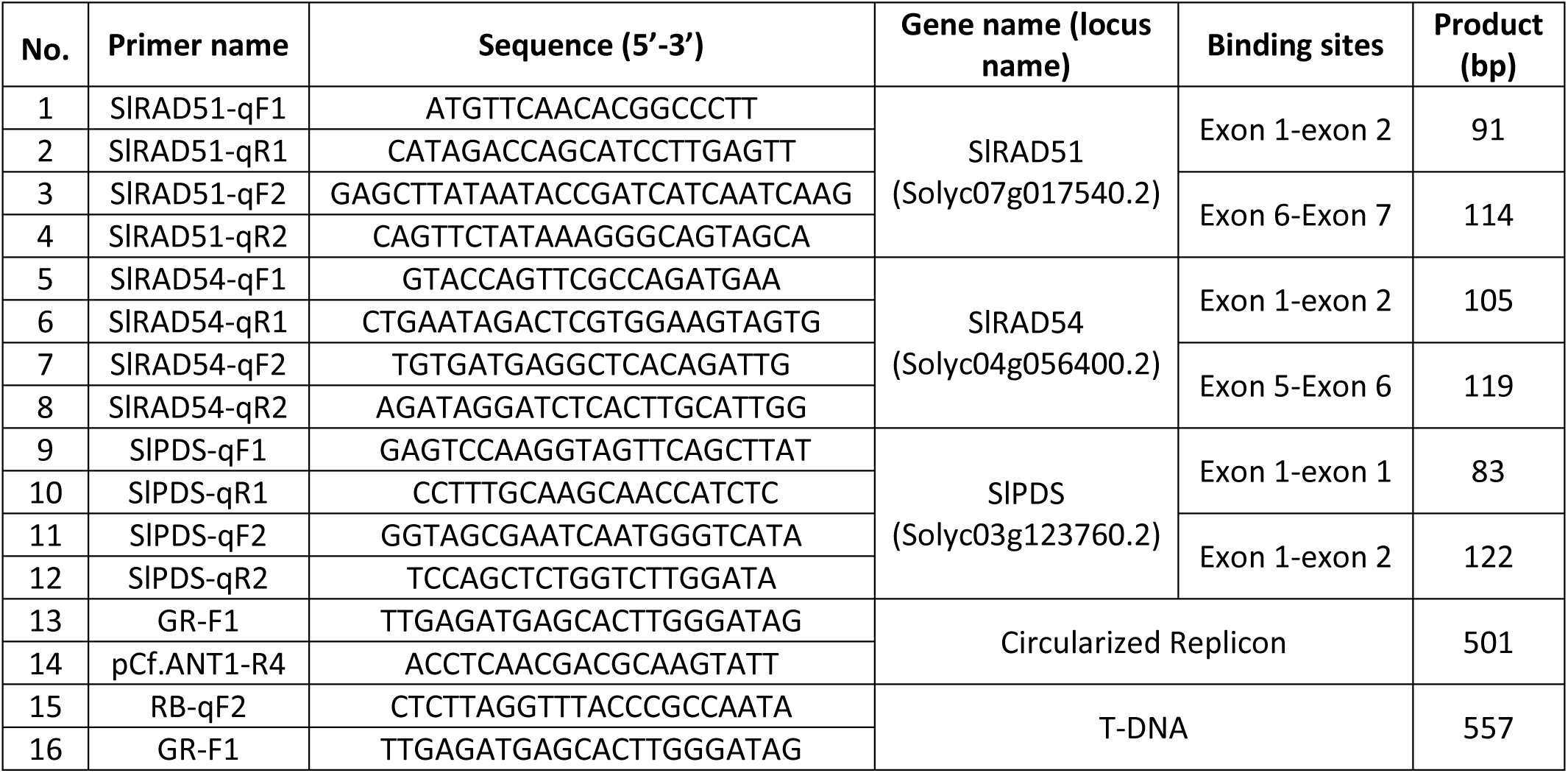
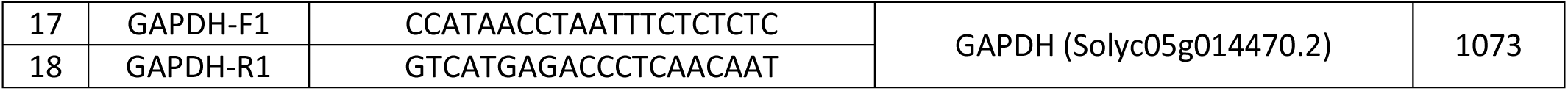

### qRT-PCR procedure and analysis of data

All of the qPCR and qRT-PCR analyses were performed following the MIQE guideline (Bustin et al., 2009). Briefly total RNAs were isolated from plant tissues using RNeasy mini Qiagen kits (cat. no. 74104, Qiagen, USA) and subjected to reverse transcription for synthesizing 1^st^ cDNA strands using QuantiTect Reverse Transcription Kit and protocol (cat. no. 205311, Qiagen, USA). At least two pairs of primers were tested for each target gene/cDNA for evaluating their efficiencies and primers pairs with ∼ 100% efficiency were ultimately used for qPCR/qRT-PCR. The similar assessment was also applied for the internal genes for normalizing the amplicon levels. qPCR/qRT-PCRs were performed using intercalating dyes (KAPA SYBR FAST Universal, cat. No. KK4601, Sigma, USA) for detecting products. Thermocycling was conducted with Illumina Eco Real-Time PCR System (Illumina, USA). Analyses of amplicon levels were performed using delta delta Cq method (Livak and Schmittgen, 2001) with internal gene/transcript of SlPDS and were plotted using Excel software (Microsoft, USA).

### SlRAD51 or SlRAD54 transcript levels expressed in transgenic pMR03 or pMR04 events, respectively. pMR02 samples were used as references

**Table A:**
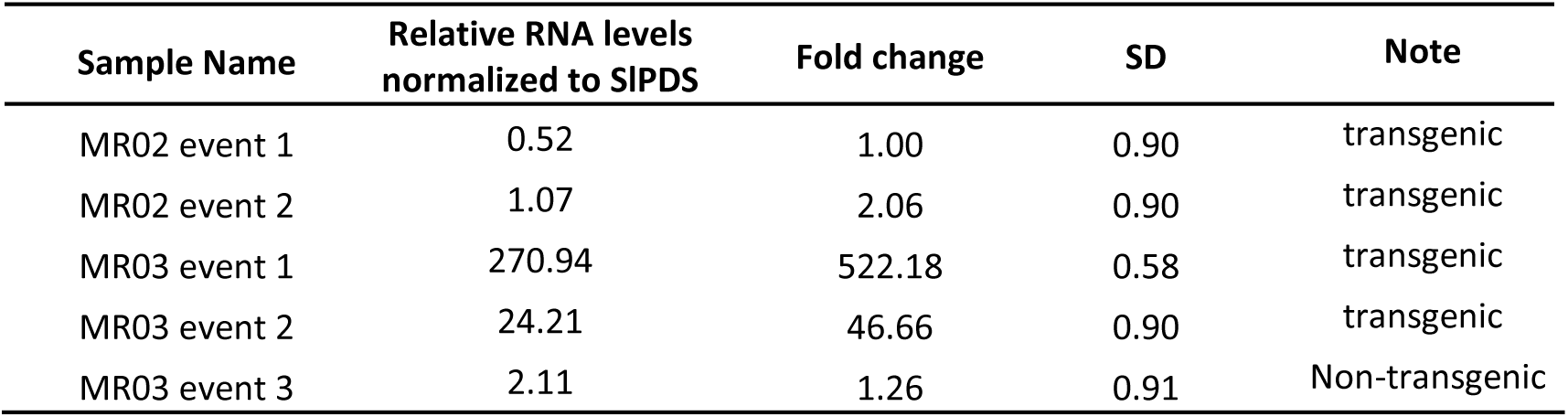
Relative transcript levels of SlRAD51 measured in the transformed events of MR03

**Table B:**
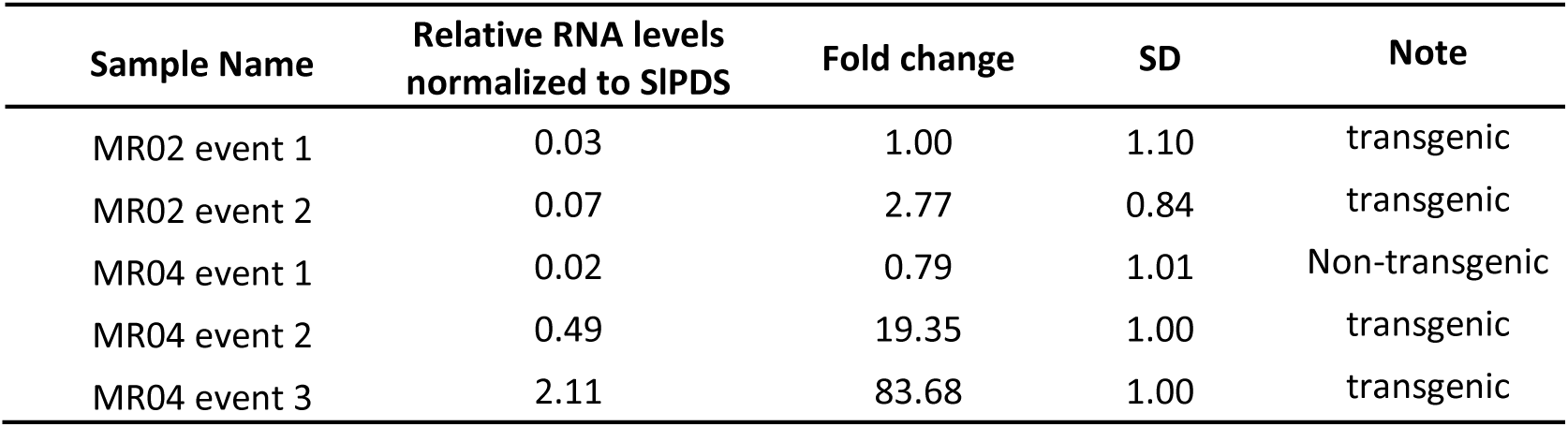
Relative transcript levels of SlRAD54 measured in the transformed events of MR04

## Southern analysis for ANT1 HDR GE1 plants

The Southern blot technique was implemented using MSU Potato Lab’s protocol (https:/msu.edu/course/css/451/LabProtocols/09SOUTHERNprocedit.pdf) with minor modification. Essentially, genomic DNAs (gDNAs) are isolated from tomato leaves using DNeasy Plant Maxi Kit (cat. no. 68163, Qiagen, Gemany). Approximately 800 µl of gDNA was eluted from column and subjected to precipitation by adding 2.5 volume of absolute ethanol and 1/30 volume of sodium acetate 3M, incubating at −80oC for 2h, centrifuging at 13rpm for 20 minutes and drying for 30 minutes in a 37oC incubator to completely remove residual ethanol. The final eluted volume was 60 µl of AE buffer with ∼500-1000 ng/µl.

Twenty micrograms of the gDNAs was digested overnight (∼18 h) at 37oC with NsiI (NEB, USA) and inactivated at 65oC for 20 minutes. The digested product was precipitated by adding 2.5 volume of absolute ethanol and 1/30 volume of sodium acetate 3M, incubating at −80oC for 2h, centrifuging at 13rpm for 20 minutes and drying for 30 minutes in a 37oC incubator to completely remove residual ethanol. The dried DNA was suspended in 30 ul of EB buffer and 6 ul of DNA loading dye 6x was added for agarose gel running. The digested product was loaded in a 0.8% agarose gel and resolved by running overnight (∼18h) at low voltage (30V). Resolved DNA bands were overnight transferred onto a Hybond N+ membrane (GE Healthcare, USA) by capillary transferring system. The blotted membrane was treated two times with high energy UV (1200×100 µJ/cm2) for crosslinking the DNAs to the membrane.

The membrane was then pre-hybridized and overnight hybridized with DIG-labeled Probe specific for the downstream homologous arm (Supplemental Figure 2) in a plastic bag. The probe was amplified by PCR using primers flanking the downstream homologous arm (see ptimer table below). Probes were DIG-labeled by random priming using a Random Primed DNA Labeling Kit (cat.no. 11004760001, Roche, Switzerland) following the manufacture’s protocol. The probed membrane was washed stringently and bound DIG probe was detected using DIG Nucleic Acid Detection Kit (cat.no. 11175041910, Roche, Switzerland) following the manufacture’s protocol. Below are details of each part conducted in the analysis.

**Primers used for making Probe**

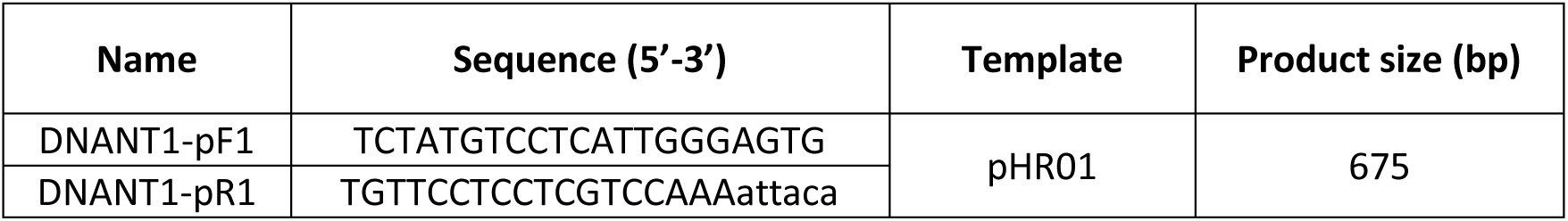

### Setting up digestion reactions

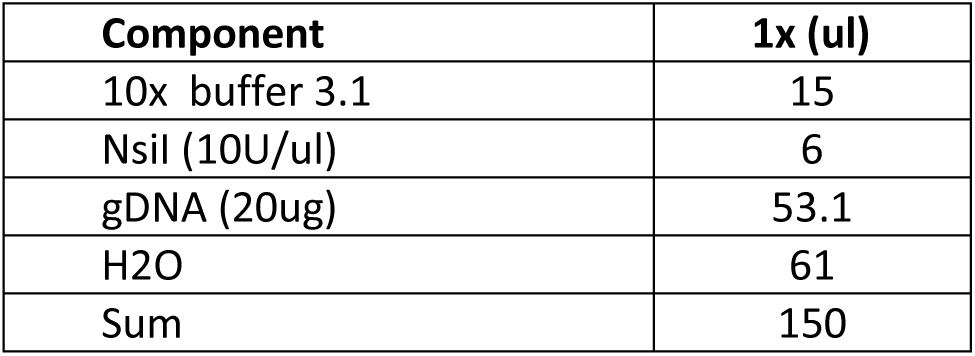

- Incubated the reaction tubes in a water-bath at 37°C for 18h.

- Inactivated the enzymes at 80°C for 20 minutes.

- Precipitated the digested gDNA with 1/30 volumes of NaOAc 3M and 2.5 volumes (250ul) of ethanol 100% (molecular biology grade) and incubated at −80°C for 4 hour.

- Centrifuged the tubes at 13rpm for 20’, 4°C and discarded supernatant.

- Dried DNA pellets at 37°C for 2h and add 30ul EB.

- Added 6ul of 6x loading dye for loading.

### Agarose gel running

- Gel size: 14.3cmx21.1cm; need to prepare ∼250ml TAE 1x, 0.8% agarose

- Run at 30V overnight (18h)

### Capillary Transferring of DNAs

- Depurination in 0.25N HCl: 1 time, 20’; agitation 30rpm.

- Rinsed briefly 2 times with H2O.

- Denaturation in SOLUTION D: 2 times, 15’ each; agitation 30rpm.

- Rinsed briefly 2 times with H2O.

- Neutralization in SOLUTION N: 2 times, 15’ each; agitation 30rpm.

- Pre-wetted Whatman paper wick with 20x SSC and put in a plastic tank. Removed air bubbles using plastic pipet.

- Pre-wetted 3 sheets of gel-sized Whatman paper and placed on the wick.

- Pre-wetted gel in 20x SSC and placed upside down on the whatman paper.

- Pre-wetted gel-sized Hybond N+ membrane in 20x SSC and placed on the gel.

- Pre-wetted 3 sheets of gel-sized Whatman paper and placed on the membrane.

- Placed paper towels (∼15cm) on the Whatman paper.

- Placed the gel casting tray on the top of the paper towel.

- Placed ∼800 g Schott bottle on the top

### Pre-hybridization

- Put the membrane in a clean plastic bag. Use gloved hands and blunt ended forceps.

- Warm 40mls of prehybridization solution to 42°C.

- Boil 1000ul of Salmon Testes DNA (10mg/ml) for 10min and then place on ice for 2 min. Add the 1000ul of DNA to the 40mls of prehybridization solution. The final concentration of DNA is 250ng/ml. This is used to block nonspecific binding sites on the blot.

- Add the prehybridization solution to the blot in the bag. Seal and place the bag in hybridization oven that has been warmed to 42°C.

- Prehybridize blot for at least 3h30 at 42°C. Longer times are possible.

### Hybridization

- Prepare hybridization solution for hybridization as follows. Hybridize a 20×30cm blot with 40mls of the buffer. The amount varies depending on the amount of stock probe that you have. Warm the Hybridization solution to 42°C.

- Reused the hypridization buffer of the first blot by warming up at 65°C for 20’

- Add 6 ul of the DIG-labelled Probe to the warm hybridization solution and mix by shaking well.

- Remove prehybridization solution from hybridization tube. (The prehybridization solution can be reused 2 times. Store at −20°C. To reuse, the Salmon DNA in the solution must be denatured. Place the solution in a 65°C water bath for 15min. The flashpoint of pure formamide is 68°C therefore <u>do not boil</u> the solution.)

- Do not let blot dry in the bag. Immediately add the hybridization solution and place in hybridization oven. Make sure the tube is balanced! Incubate overnight at 37°C.

### DETECTION

- After hybridization and stringency washes, rinse membrane briefly with WASHING BUFFER about 2-5 min in a plastic tray with the DNA side up.

- Incubate membrane in 100mls of BUFFER 2 for 30 min.

- Dilute anti-DIG-AP conjugate 150 mU/ml (1:5,000 dilution) =10ul in 50ml of fresh BUFFER 2.

- Incubate membrane for 30 min in the antibody solution on shaker either in a plastic tray. Ensure that the solution is covering the entire blot with gentle agitation (30rpm).

- In a plastic tray wash the membrane 2X 15 min. with 100ml of WASHING BUFFER. Apply gentle agitation (30rpm).

- Equilibrate membrane 2-5 min. in 20 ml BUFFER 3.

- Add 400 μl of NBT/BCIP stock solution (vial 4) to 20 ml of Detection buffer. Note: Store protected from light!

- Add the solution to the membrane for color development and keep in dark up to 16h.

- Taking photograph of the blot.

### Southern blot buffers

**<u>A.</u> <u>Agrose gel treatment</u>**

1) 0.25N

2) Solution D, Denature solution: 1.5M NaCl and 0.5M NaOH

3) Solution N, Neutralizing solution: 0.5M Tris, 1.5M NaCl and 1mM EDTA; **pH to 7.5**

4) 20 X SSC: 3.0M NaCl, <u>0.3M NaCitrate;</u> **pH to 7.0**

**<u>B. Hybridization and washing</u>**

1. Prehybridization and Hybridization solution:

5X SSC, 2% Block solution, 0.1% N-lauroylsarcosine, 0.2% SDS

0.5 volumes of <u>Pure</u> Formamide (deionized)

Aliquot in 50ml tubes and store at −20°C until use.

Salmon Testes DNA and Probes are added just before use.

2. 2X Wash Solution: 2X SSC, 0.1% SDS

3) 0.5X Wash Solution: 0.5X SSC, 0.1% SDS

**<u>C. Detection</u>**

**Solutions:**

**1) Maleic Acid Buffer**

0.1M Maleic acid, <u>0.15M NaCl,</u> pH to 7.5 using NaOH pellets.

**2) WASHING BUFFER**

Maleic Acid Buffer, Make the same as above 0.3% Tween 20

**3) Blocking stock solution 10x conc**.

Blocking reagent from BMB, 10% (w/v) in maleic acid buffer. Dissolve blocking reagent by constantly stirring on a heating block at (65°C). Do not boil the solution. It is difficult to get into solution and may take several hours. Be sure that all of it has dissolved and then autoclave. Store at 4°C. The solution is opaque.

**4) BUFFER 2** (make fresh for each use)

1% Blocking Buffer in Maleic Acid Buffer

**5) BUFFER 3**

0.1M Tris-HCl, 0.1M NaCl, 50mM MgCl2; pH to 9.5

**7) Random Primed DNA Labeling Kit** (cat.no. 11004760001, Roche, Switzerland)

**8) DIG Nucleic Acid Detection Kit** (cat.no. 11175041910, Roche, Switzerland)

